# Rapid selection of transgenic mammalian cells via diphtheria toxin resistance

**DOI:** 10.1101/2024.08.26.609662

**Authors:** David Scherer, Steffen Honrath, Jean-Christophe Leroux, Michael Burger

## Abstract

The ability to generate stable transgenic mammalian cell lines is crucial to the investigation of gene functions and the production of recombinant proteins. Currently, mammalian cells can be readily transfected in cell culture settings via both viral and nonviral vectors to induce transgene expression. However, there is an unmet need for efficient selection of transfected cells, since current methods involve rather inefficient antibiotic selection protocols or require the coexpression of fluorescent marker proteins, followed by laborious cell sorting procedures. Our aim was to implement a rapid and efficient in situ selection approach for transgene-expressing human cells, using an engineered diphtheria toxin (DT) resistance-based selection, referred to as selecDT. We demonstrated that selecDT is expressed on the surface of modified human cells, provides efficient protection from DT by inactivating its uptake receptor, and, therefore, enables selection. Current antibiotic-based methods require selection periods of more than a week and often achieve only limited cell enrichment. With selecDT, one day of selection is sufficient to obtain nearly 100% enrichment. The DT resistance described herein may thus positively impact biotechnological processes and biomedical research.

---

Currently, recombinant protein production in bacteria is a relatively simple process. The bacteria are first transformed with plasmids containing an expression cassette for the protein of interest and a specific antibiotic resistance gene. In the presence of the antibiotic, untransformed bacteria are rapidly eliminated, while the expression of the resistance gene efficiently protects the transformed cells. Unfortunately, comparable selection tools for mammalian cells are currently not available. Although antibiotics are commonly employed, selection efficiencies are generally low, vary substantially among cell lines, and necessitate prolonged selection periods ranging from one to four weeks. For example, HeLa cells transfected with plasmids containing the neomycin resistance gene (neoR) are typically selected at antibiotic (G418) concentrations of up to 2 mg/mL. While the applied antibiotic concentrations are toxic to ‘nonresistant’ cells, near-immediate cell death is not achieved, necessitating prolonged application of selection pressure for more than a week. High antibiotic concentrations and prolonged treatment also decrease the viability of the ‘resistant’ cell population, further resulting in undesired delays in the selection process^1^. Moreover, cytotoxicity varies widely depending on the cell line and antibiotic, necessitating laborious optimization for each antibiotic–cell combination.

In this study, we present a selection system based on a recombinant cell surface protein that renders cells highly resistant to diphtheria toxin (DT). DT, a 535-amino acid exotoxin secreted by *Corynebacterium diphtheriae*, comprises two disulfide-linked fragments responsible for cell entry and toxicity. Fragment A inhibits ribosomal protein synthesis, whereas fragment B is responsible for binding to cellular receptors to induce endocytosis and subsequent endosomal escape^2^. In the cytoplasm, only a few copies of fragment A are sufficient to completely inhibit ribosomal protein synthesis, leading to cell death. The inhibition of protein synthesis occurs when DT conjugates ADP1ribose to a modified histidine residue (diphthamide) on elongation factor 2 (EF2) (Figure 1). Previous studies have attempted to introduce cellular resistance against DT by knocking down enzymes involved in diphthamide synthesis^2–4^. However, the diphthamide modification in EF2 is highly conserved and important for translational fidelity; therefore, its removal might interfere with cellular fitness^5–8^. An alternative approach to introducing DT resistance is the deactivation of the cellular DT receptor, the immature form of heparin-

**Figure 1:**
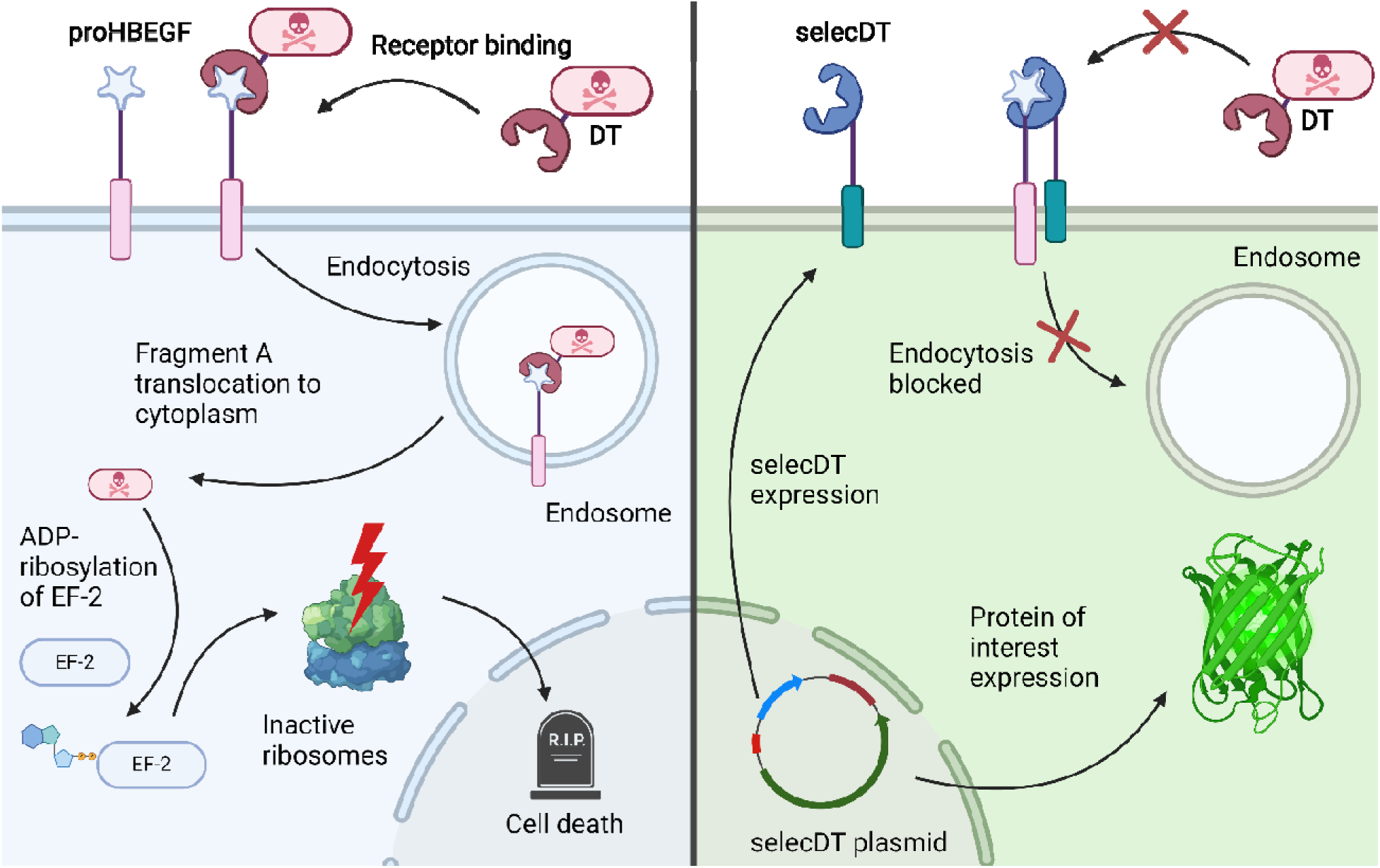
Schematic representation of the cellular response to DT exposure resulting in rapid cell death (left). Model of the response of selecDT cells to toxin exposure (right). Transgene expressing cells block proHBEGF on the ce ll surface with selecDT, inhibiting toxin endocytosis and cell death.

binding EGF-like growth factor (proHBEGF). While the mature form of HBEGF is essential for tissue development and regeneration in vivo, its deactivation is expected to have minimal adverse effects on cell viability in a cell culture setting. Because only proHBEGF is responsible for toxin endocytosis, inhibiting this interaction can make cells highly resistant. This can be observed in mice, which are naturally resistant to DT due to a point mutation (E141H) in the DT binding site of murine proHBEGF^9,10^. Consequently, mouse cells do not endocytose DT and are thus protected from its toxic effects. The fact that mouse cells are virtually immune to DT has long been exploited in vivo for the selective depletion of xenograft cells in murine models^11^. In a recent study, CRISPR-based genetic introduction of an E141K mutation in human proHBEGF (analogous to E141H in the mouse protein) resulted in DT resistance similar to that of mouse cells^12^.

Both approaches mentioned above rely on genomic modifications to make human cells resistant to DT. This is not ideal since genetic modifications become effective only once the wild-type proteins, i.e., proHBEGF or the diphthamide synthesis machinery, have been degraded. This process can take several days, depending on the turnover of the targeted proteins. Therefore, both protocols incorporate a lag period of 3 days posttransfection to render the cells DT resistant before beginning the selection process. These approaches are not substantially faster than traditional antibiotic-based selection protocols. Here, we describe a form of recombinant protein-based resistance that quickly inactivates proHBEGF on the cell surface to provide immediate protection against the toxin and does not necessitate the introduction of genomic modifications.

Studies have shown that the addition of high amounts of a soluble nontoxic DT mutant (K51E, E148K) or DT receptor-binding fragment can protect cells from the cytotoxicity of native DT^13^. To mimic this protection and create a resistance factor for selection, we genetically fused the receptor-binding subunit of DT to the transmembrane (TM) domain of CD70 (Figure 1). The TM domain localizes the recombinant protein to the plasma membrane, whereas the receptor-binding subunit binds proHBEGF. The interaction of these two membrane proteins sterically inhibits soluble toxin binding. Additionally, the binding of proHBEGF triggers receptor endocytosis, resulting in its clearance from the cell surface. Consequently, we established a novel selection marker, termed selecDT, which potently blocks and removes the DT uptake receptor from the cell surface and thus renders the cells DT resistant. By leveraging selecDT, we established a straightforward protocol to rapidly generate stable human producer cell lines. To keep the protocol simple and fast, we used two nonviral plasmid transfection systems. First, the TFAMoplex, which was developed by our group, allowed efficient HeLa cell transfection via human mitochondrial transcription factor A (TFAM)-based nanoparticles^14^. Second, the standard poly(ethylenimine) (PEI) transfection system was used to transfect HEK293 FreeStyle™ suspension cultures. Stable genome insertions were performed via a highly efficient DNA recombinase (cp36)^15^.

## Results

First, we aimed to confirm that selecDT localizes to the plasma membrane and interacts with the DT receptor proHBEGF. Therefore, proHBEGF and the selecDT protein were genetically fused to the fluorescent reporters eGFP and mScarlet3, respectively, and were transiently overexpressed in HeLa cells (Supplemental Information Fig. 1). Confocal microscopy revealed that selecDT-mScarlet3 localized primarily to the plasma membrane, with a pronounced punctate intracellular signal, and that recombinant proHBEGF-eGFP was located predominantly on the plasma membrane. When both labeled proteins were coexpressed in the same cells, the overall localization of the proteins was not obviously altered, with strong colocalization observed on the plasma membrane (Supplemental Information Fig. 1, bottom panel). Next, we investigated whether selecDT could indeed inhibit the uptake of soluble DT. To this end, we produced a recombinant probe composed of the receptor-binding subunit of DT genetically fused to an mCherry fluorophore (mCherry-DTrb), similar to Sugiman-Marangos et al.^13^ When HeLa cells were incubated with this fluorescent probe (10 nM), punctate staining of the cells was evident (Figure 2A), compared with the same cells without mCherry-DTrb (Supplemental Information Fig. 2). The punctate staining suggested that, upon binding, mCherry-DTrb is able to trigger endocytosis similar to the full-length toxin. When HeLa cells transiently overexpressing recombinant eGFP-labeled proHBEGF were incubated with mCherry-DTrb, the punctate staining was even more pronounced; in addition, we observed significant colocalization with labeled proHBEGF on the plasma membrane (Figure 2B). The colocalization of mCherry-DTrb with proHBEGF-eGFP indicated that our recombinant probe is able to bind the native DT receptor. On the other hand, almost no mCherry signal was detected in cells coexpressing selecDT and soluble eGFP as a marker protein, supporting our hypothesis that selecDT reduces the uptake of DT by blocking interaction with proHBEGF (Figure 2C and Supplemental Information Fig. 3). Note that untransfected neighboring cells retained the punctate mCherry-DTrb staining (yellow arrowheads in Figure 2C). These findings suggest that the effect of selecDT is confined to the transfected cells expressing the resistance-conferring gene, further validating the potential of selecDT as a selection marker.

**Figure 2:**
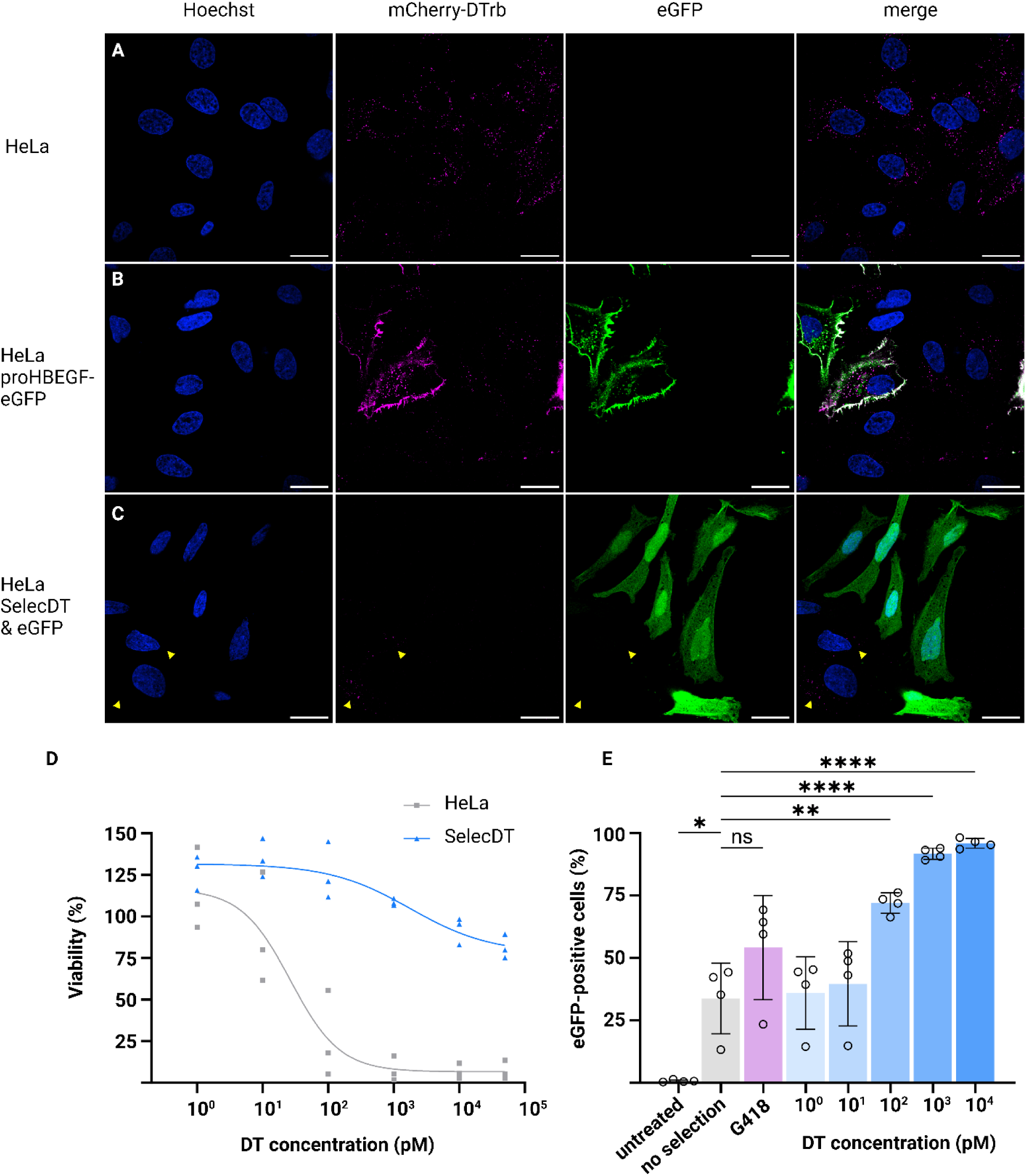
Characterization of SelecDT in HeLa cells. **A, B, C:** Confocal microscopy images of cells incubated with 1 nM mCherry-labeled proHBEGF-binding subunit (magenta). The nuclei were stained with Hoechst (blue). The scale bars represent 25 µm. All images were acquired via identical microscope and analysis settings. **A**, Naïve HeLa cells. **B**, HeLa cells overexpressing proHBEGF-eGFP (DT receptor) (green). **C**, selecDT cells coexpressing cytoplasmic eGF (green). Arrowheads indicate a punctate mCherry signal in untransfected cells. **D**, Viability of naïve and selecDT HeLa cells upon DT titration measured via MTS. The data are plotted as the means of 3 independent biological replicates, each consisting of 3 technical replicates (n=3). The fitting curve indicates the IC50 value of naïve HeLa cells at 26 pM DT. The cells protected by selecDT retained 80% viability at the highest DT concentration, preventing the corresponding IC50 from being determined. **E**, Flow cytometry analysis of HeLa cells coexpressing a marker transgene (eGFP) and both resistance factors selected with G418 (magenta), DT (blue) or mock (gray) after one day of selection followed by one day of recovery in medium without any selection. The data are plotted as the means ± SDs of 4 independent biological replicates, each consisting of 3 technical replicates (n= 4).

As a result of these findings, we investigated if SelecDT was indeed able to protect cells from DT in a viability assay. We generated a HeLa cell line expressing selecDT, neoR and a marker protein (mCherry) and incubated the cells with increasing amounts of DT or G418 (Figure 2D and Supplemental Information Fig. 4). At DT concentrations above 100 pM, an almost complete loss of viability was observed for the naïve cells, whereas the selecDT-expressing cells retained 80% viability at the highest tested concentration of 50 nM. The observed toxin tolerance exhibited by selecDT cells is similar to that of a receptor knockout cell line previously described by Sugiman-Marangos *et al*^16^. At DT concentrations of 1, 10, and 50 nM, the difference between the viability values of the resistant and naïve cells exceeded 90%. This resulted in a broad selection window between 100 pM and at least 50 nM toxin, which represents a significant improvement over the narrow selection window of traditional antibiotics such as G418 (Supplemental Information Fig. 4). Next, we investigated whether the broad selection window of the selecDT system would translate to efficient selection of transiently transfected HeLa cells. We transfected a plasmid containing the genetic information for neoR, selecDT and a green fluorescent protein (eGFP) into HeLa cells. TFAMoplex transfection was performed with appropriate DNA concentrations to obtain approximately 30% eGFP-positive cells (Figure 2E). The transfected cells were incubated for one day with increasing DT concentrations or, as a control, with 2 mg/mL G418. This was followed by one day of recovery in standard growth medium. On day three posttransfection, flow cytometry analysis of eGFP fluorescence revealed that a DT concentration of 100 pM resulted in a significant increase in the percentage of eGFP-positive cells to 75%. Selection with 1 or 10 nM DT enriched the eGFP-positive cell population to nearly 100%. When selection was performed with G418, the fraction of cells expressing eGFP increased, but not significantly, compared with that in the negative control without selection. When the experiment was performed at relatively high initial transfection rates, the required amount of DT could be reduced even further (Supplemental Information Fig. 5). These results indicate that selecDT can indeed be applied as a selection tool for human cells.

Since selecDT can be used to apply selection pressure on transiently transfected cells, we tested its ability to rapidly generate cell lines that contain genomic insertions of an expression cassette. We employed the recently discovered cp36 DNA recombinase from *Clostridium perfringens*^15^. Cp36 is able to catalyze recombination between two DNA sequences originating from the bacterial genome (attB) and phage genome (attP) with high efficacy. The human genome naturally contains a defined series of pseudo-attB sites that the promiscuous cp36 can use for efficient integration. We incorporated the corresponding cp36 attP site on a donor plasmid along with expression cassettes for neoR, selecDT, an eGFP maintenance marker and a protein of interest (mCherry). To validate the ability of this plasmid to be stably integrated into the human genome, we cotransfected HeLa cells with both the donor plasmid and a cp36 integrase expression plasmid. To assess the success of donor plasmid genomic insertion, we quantified the fraction of mCherry-positive cells 3 weeks posttransfection *via* flow cytometry (Figure 3A). A sustained mCherry signal was observed only when both the attP-containing donor plasmid and the cp36 integrase plasmid were cotransfected. We quantified the fraction of HeLa cells with persistent transgene expression at 36%, which is similar to the value reported by Durrant *et al.* in K562 cells^15^.

**Figure 3:**
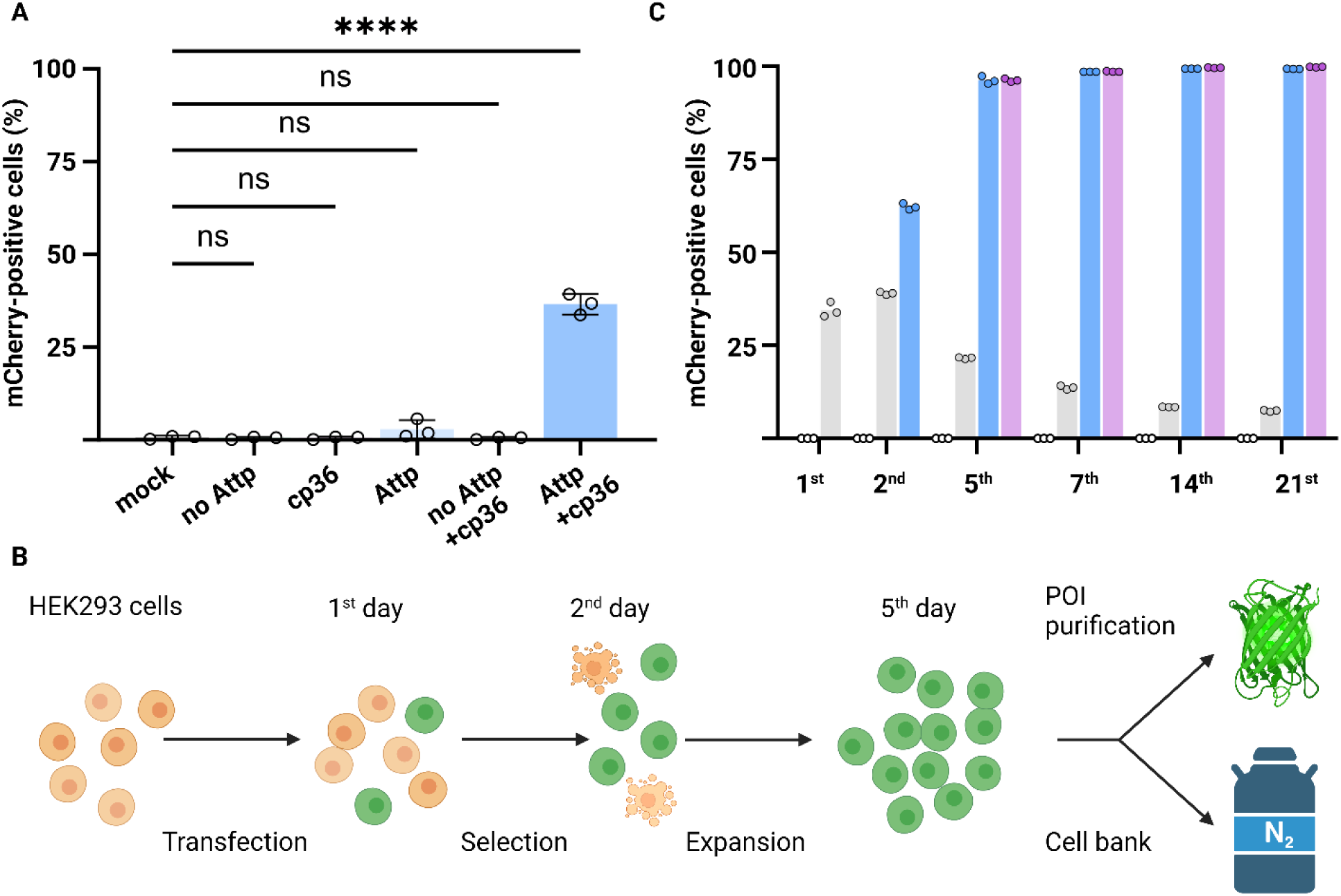
Expression of stably integrated transgenes in HEK293 and HeLa cells. **A**, Flow cytometry analysis of transgene (mCherry) expression in HeLa cells three weeks posttransfection (blue); the data are plotted as the means ± SDs of 3 independent biological replicates, each conducted with 3 technical replicates (n= 3). **B**, Schematic representation of the cell line generation protocol for recombinant protein production, with transgene-expressing cells indicated in green. **C**, Flow cytometry analysis of transgene (mCherry) expression in HEK293 cells on the first, second, fifth, seventh, 14^th^ and 21 ^st^ day posttransfection. DT selection was performed on day two. For each day, the mock control (white circles), the transfected but not selected cells (gray), cells selected with DT (blue), and cells first selected with DT and subsequently cultivated in medium containing G418 (magenta) are plotted. All the data are presented as the fraction of the mCherry positive cell population of the 3 technical replicates (n= 3).

These results confirm that the cp36 recombinase is a simple tool for stable integration of large DNA segments into the unmodified human genome. Finally, we aimed to create a straightforward protocol to generate human producer cell lines. For this purpose, we combined the integration and selection tools described above with PEI transfection of the widely available HEK293 FreeStyle™ suspension cells. We evaluated mCherry expression in HEK293 cells at various stages posttransfection (Figure 3B). Initial transfection resulted in the expression of the mCherry marker in 30% of the cells (Figure 3C). The cells were subsequently split, and half were incubated with 1 nM DT for 24 h, which resulted in a moderate increase in the mCherry-positive fraction to approximately 60%. The selected cells were again divided equally, and cultivation of the first half was continued in FreeStyle™ growth medium, whereas the second half was cultivated in maintenance medium (FreeStyle™ growth medium supplemented with 200 µg/mL G418). In general, in cell culture settings, the expression of introduced transgenes is maintained via the addition of low amounts of a selection antibiotic. Low selection pressure is necessary to prevent epigenetic silencing of exogenous DNA; in addition, it allows the continuous removal of cells that do not productively integrate the donor plasmid into their genome and therefore lose the transgene. In theory, DT could also serve this purpose; however, to minimize the application of the highly toxic DT we decided to use G418 to maintain the selection pressure instead. The neoR gene offers an inexpensive, simple and long-established maintenance marker, and G418 can more easily be separated from the much larger recombinantly expressed protein of interest compared to DT. The 3-day expansion period allowed surviving selecDT-expressing cells to proliferate, leading to an increase in the transgene-expressing fraction to nearly 100% (Figure 3C). Even without G418, after 21 days, no cells exhibited loss of transgene expression, indicating that all the selected cells had successfully integrated the plasmid into their genome. Therefore, we concluded that DT selection was sufficient to generate the stable cell line. To ensure the long-term expression of the protein of interest and prevent epigenetic silencing, however, it is advisable to include G418 after selection.

## Discussion

Genetic fusion of the receptor-binding subunit of DT to a human TM domain allows the expression of a desired transgene to be linked to almost immediate DT resistance. DT is commercially available and well studied, which makes this tool easily accessible. This system, selecDT, allowed us to shorten the required selection time for transgene-expressing cells by more than one week. For example, HeLa cells, which are resilient to commonly used antibiotics, were successfully selected overnight.

Antibiotics are typically small molecules that indiscriminately enter cells. Commonly used antibiotics such as puromycin, hygromycin B and neomycin are inactivated in the cytoplasm by the corresponding resistance-conferring proteins through enzymatic phosphorylation or acetylation^10,17–23^. Incomplete antibiotic inactivation, however, has the potential to induce cellular stress and impact protein synthesis^24–26^. Unlike antibiotics, DT is a membrane-impermeable macromolecule that requires active, receptor-mediated cellular uptake^27^. Toxin uptake is mediated exclusively by the proHBEGF receptor, which is widely expressed on human cells. This has two important advantages. First, selecDT blocks the only cellular entry pathway of DT, limiting the possibility of intracellular side effects. Second, the ubiquitous expression of proHBEGF makes selecDT highly translatable to other cell lines with minimal requirements for optimization. The broad applicability of this method is further enhanced by the orthogonality of selecDT to current antibiotics. This complementarity allows selecDT to be integrated into existing protocols that require multiple selection moieties. One possible disadvantage of selecDT is that the cells need to express an artificial protein on their plasma membrane. Both the DT fragment and the TM domain may lead to unwanted changes on the cell surface, potentially interfering with the study of plasma membrane-related biological processes. Generally, the overexpression of any protein can generate cellular artifacts, which should be considered in the design of cell experiments. Our selection system should, however, greatly simplify the generation of protein-producing cell lines.

In addition to the integrase-based genome insertion demonstrated in this work, more targeted insertion strategies, such as CRISPR-based genome modifications, which typically exhibit lower integration rates, may also benefit from the improved selection capacity of selecDT. In addition to the simple generation of stable cell lines, the high selection pressure of selecDT may enable the development of novel strategies for directed protein evolution. Efforts to exploit the potency of DT and similar toxins to treat diseases, such as cancer, have been ongoing for decades^28^. To create these treatments, DT was genetically engineered to replace the native receptor-binding subunit by another receptor-binding moiety, for example, targeting the IL2 receptor on tumor cells^29–31^. In this example, a modified version of selecDT containing an IL2 receptor ligand would enable selection against this specific recombinant fusion toxin. SelecDT variants that target orthogonal receptors may therefore enable multiple simultaneous selections with high efficacies. The herein described combination of selecDT with efficient transfection and integration methods simplifies transgenic mammalian cell culture, bringing it closer to bacterial systems in terms of accessibility.

## Methods

### 1. Cloning

Plasmid backbones were obtained from either Thermo Fisher Scientific (Waltham, MA, USA) (pcDNA3.4) or Addgene (Watertown, MA, USA) (pIRES-EGFP-puro, CMV-mEGFP-CREB, pAF92 and pU6-(BbsI)_CBh-Cas9-T2A-mCherry). pIRES-EGFP-puro was a gift from Michael McVoy (Addgene plasmid 45567). CMV-mEGFP-CREB was a gift from Ryohei Yasuda (Addgene plasmid 137001). pAF92 was a gift from Andrew S. Flies (Addgene plasmid # 135929). pU6-(BbsI)_CBh-Cas9-T2A-mCherry was a gift from Ralf Kuehn (Addgene plasmid # 64324). pET His6 TEV LIC cloning vector (1B) was a gift from Scott Gradia (Addgene plasmid # 29653). Recombinant DNA encoding for the transgenes was synthesized by Twist Bioscience HQ (South San Francisco, CA) (Supplemental Information Table 1). The remaining DNA sequences were obtained via polymerase chain reaction (PCR) using Phusion polymerase (Thermo Scientific, Waltham, MA, USA) and the primer sequences listed in Supplemental Information Table 2. The sequence of the resulting PCR fragments are listed in Supplemental Information Table 3. Plasmids were generated with restriction enzyme-based cloning techniques. The insert and backbone DNA were digested with the appropriate restriction enzymes (Fast Digest, Thermo Fisher Scientific (Waltham, MA, USA)) as indicated in Supplemental Information Table 4 and purified using a PCR purification Kit (QIAGEN, Hilden, Germany). Subsequent ligation with T4 DNA Ligase from NEB (Ipswich, MA) was followed by transformation into chemically competent *Escherichia coli* DH5alpha (Promega AG, Dubendorf, Switzerland). To purify the resulting plasmids, bacteria were grown overnight and the plasmids isolated from the resulting overnight cultures using Qiagen QIAprep Spin Miniprep Kit (Hilden, Germany) followed by sequencing (Microsynth AG, Balgach, Switzerland). The full plasmid sequences are disclosed in Supplemental Information Table 5.

**Table 1:**
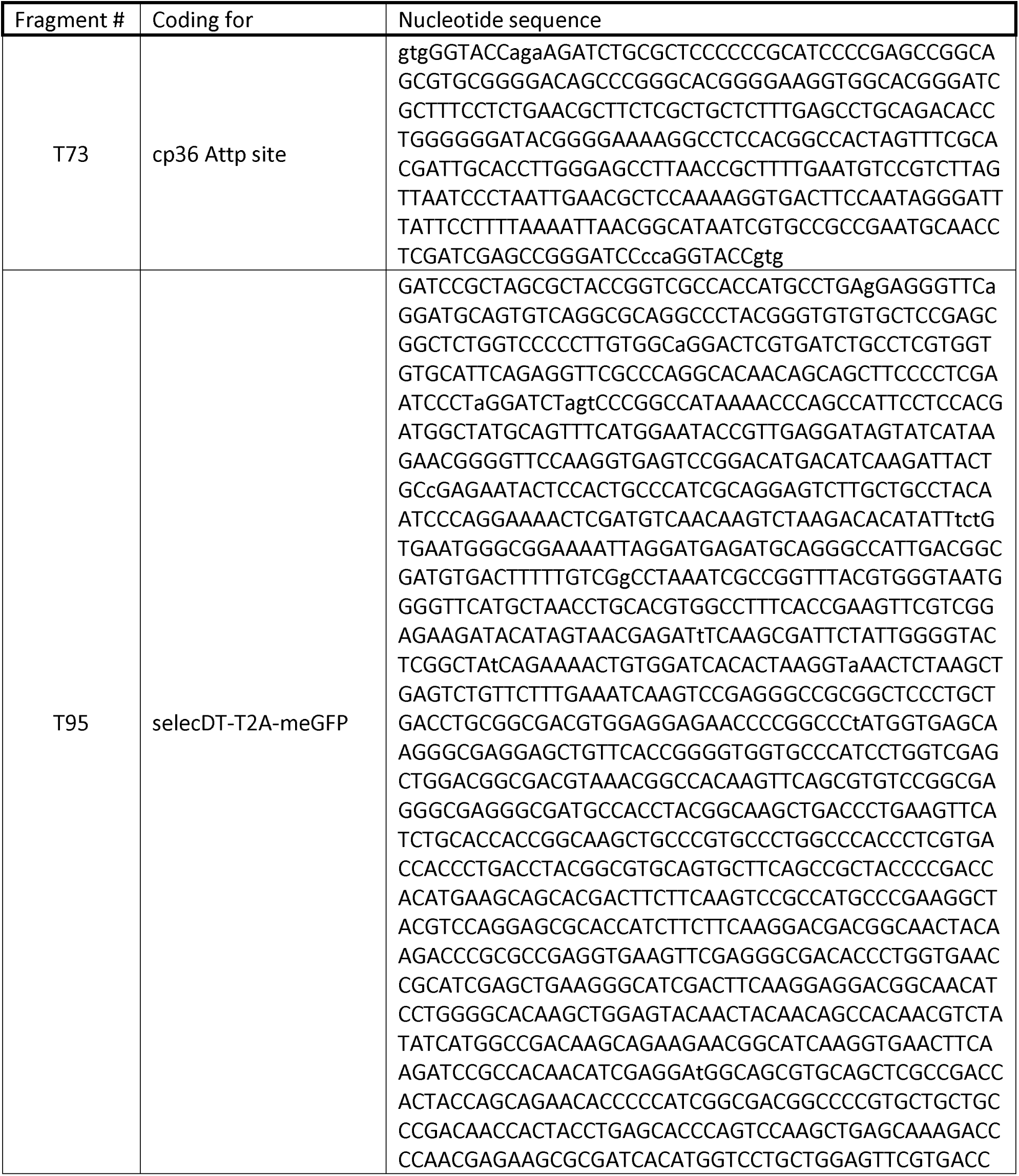
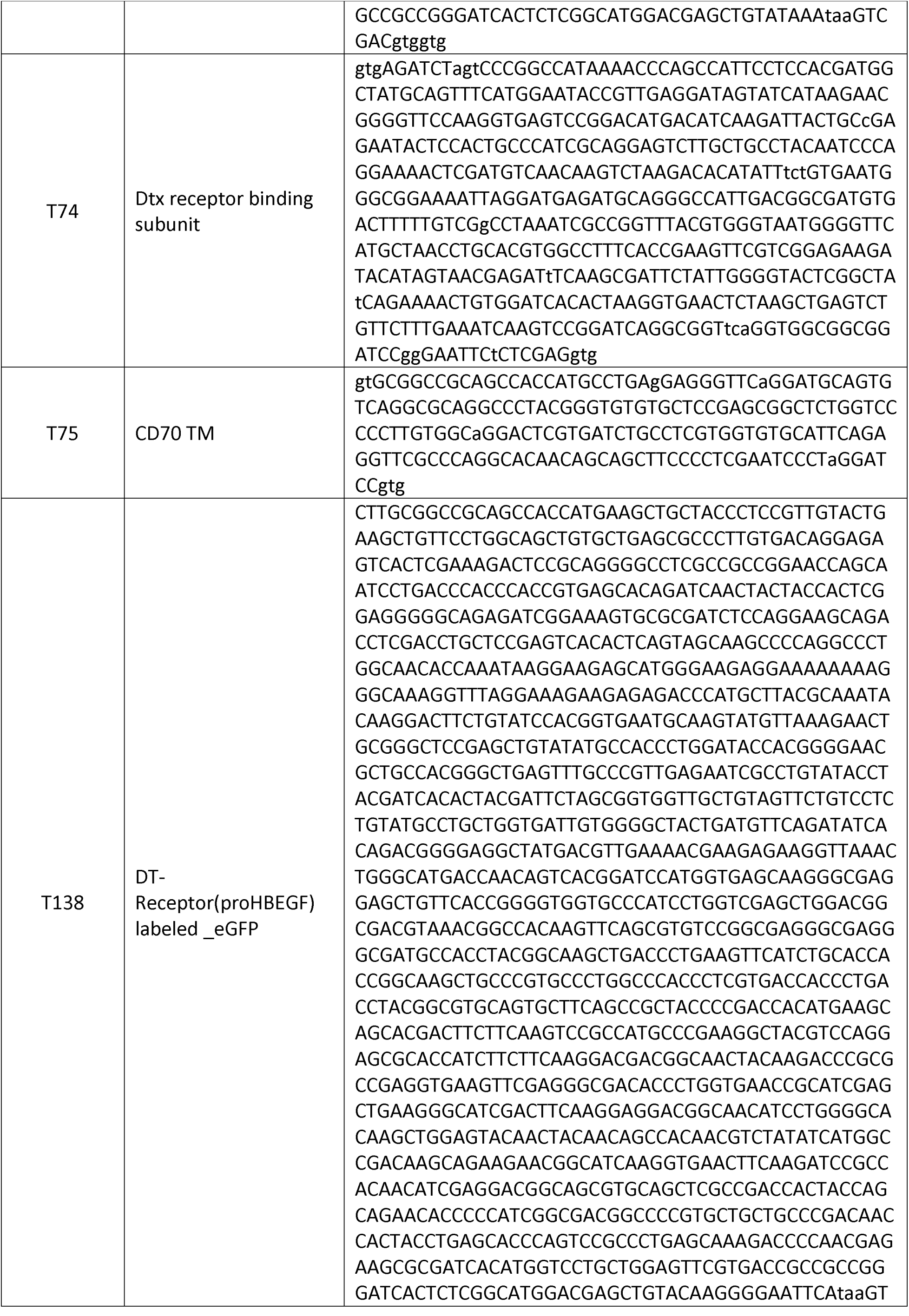
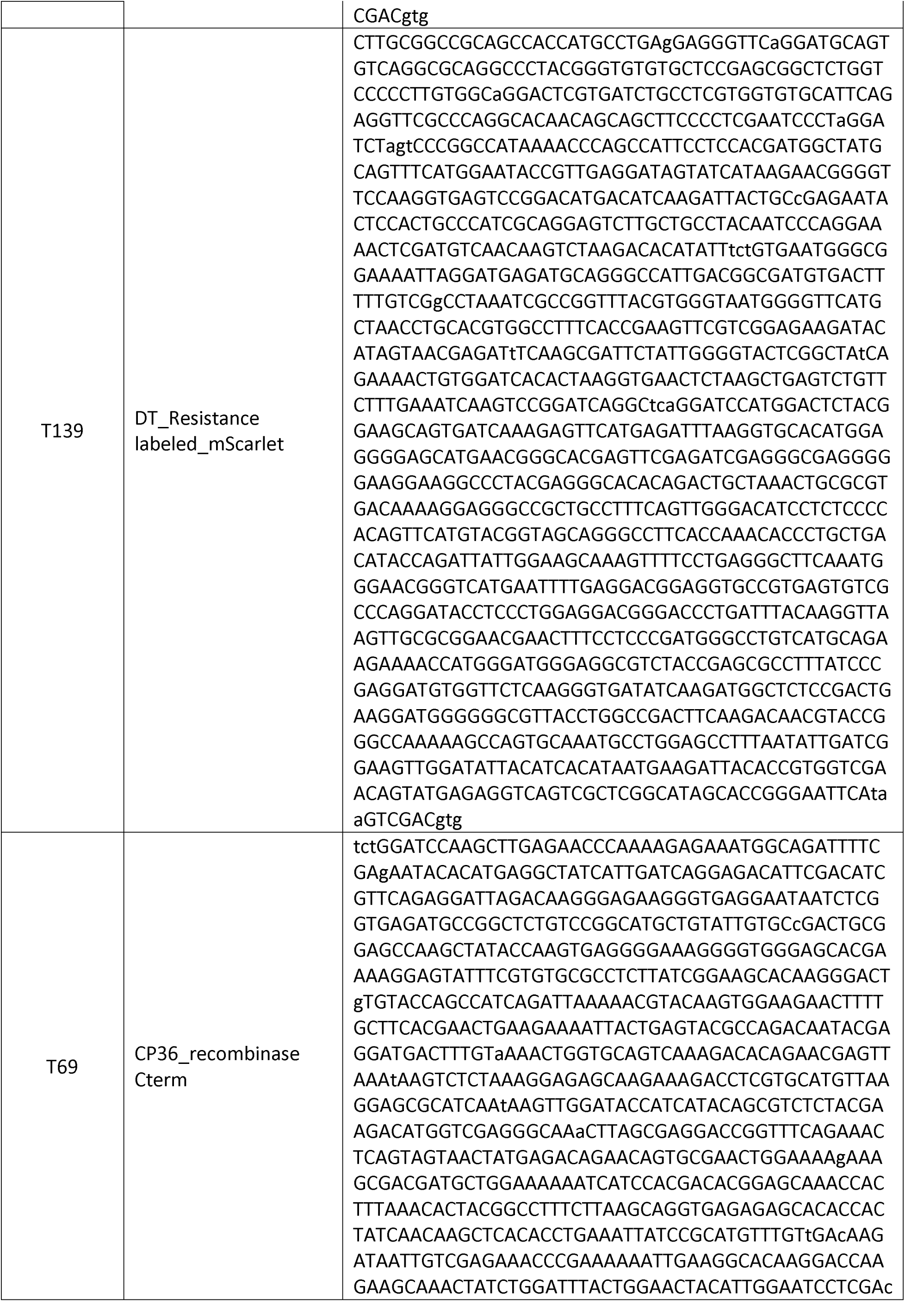
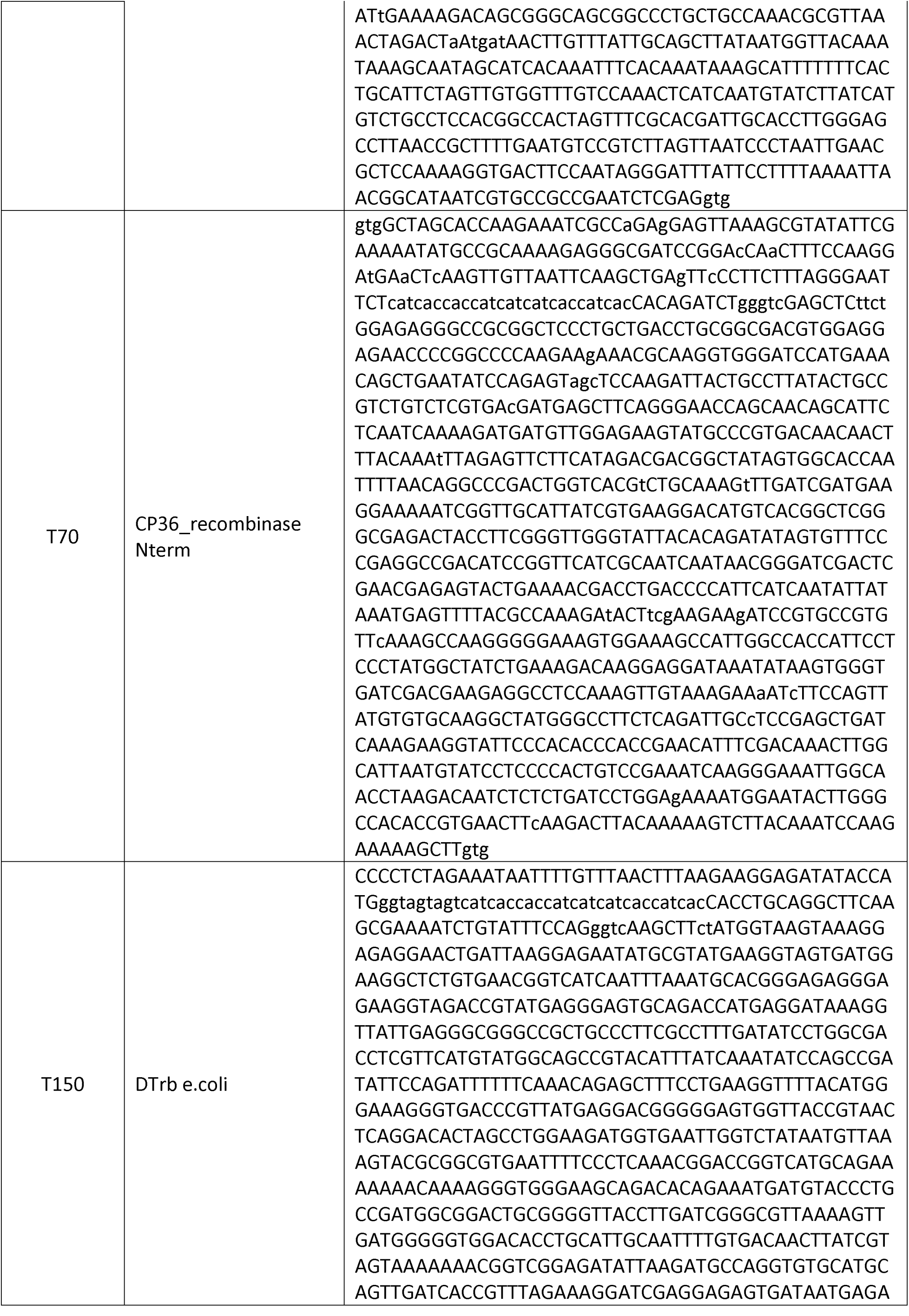
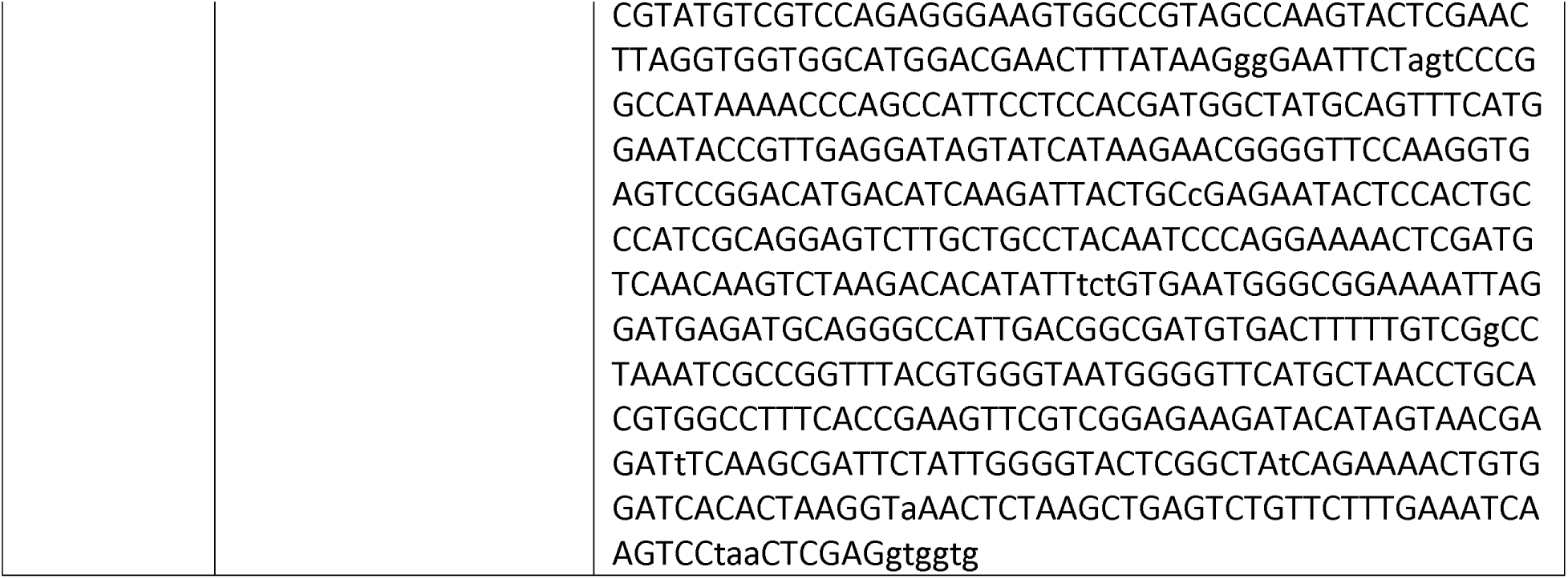
Recombinant DNA fragments:

**Table 2:**
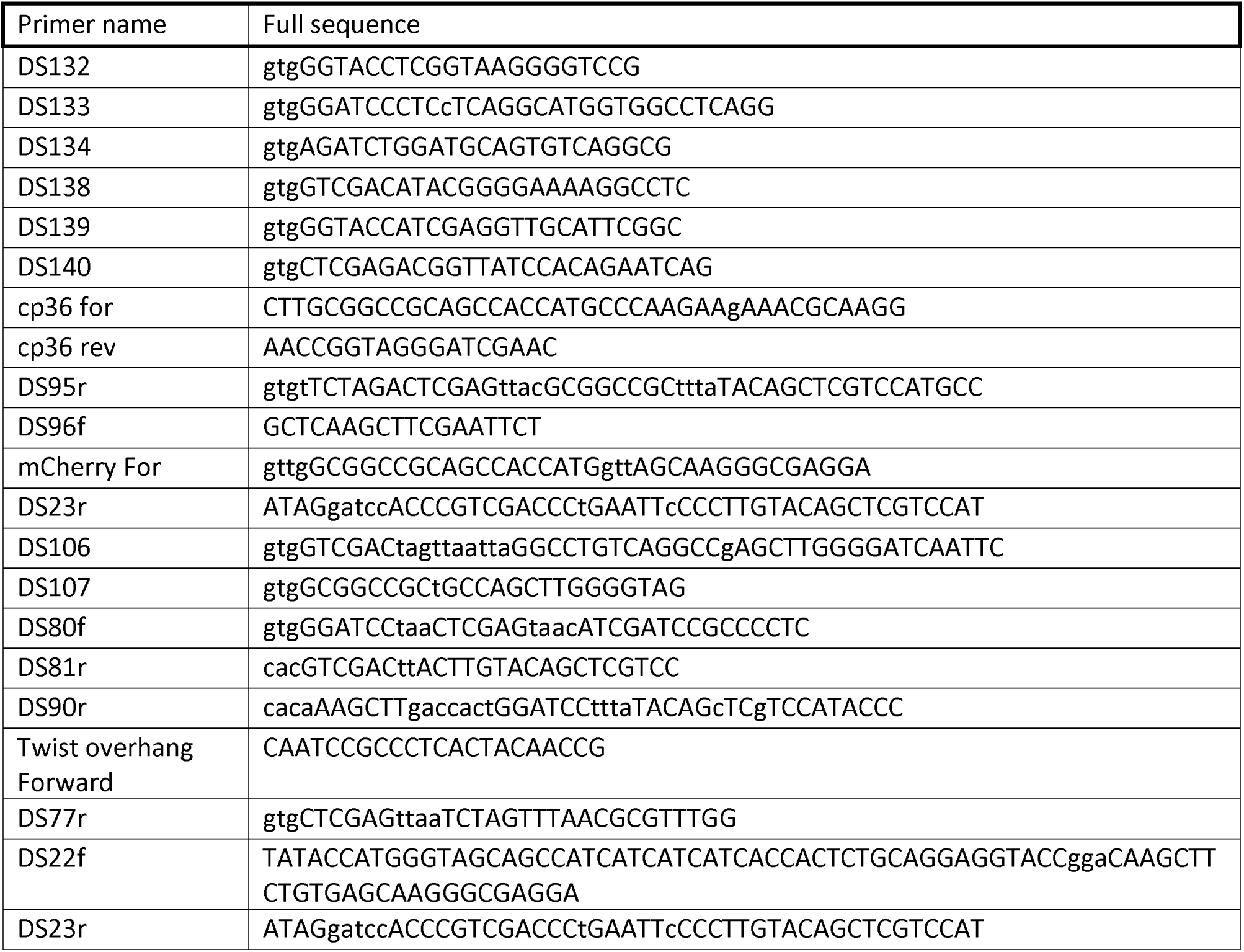
Primers used for PCRs:

**Table 3:**
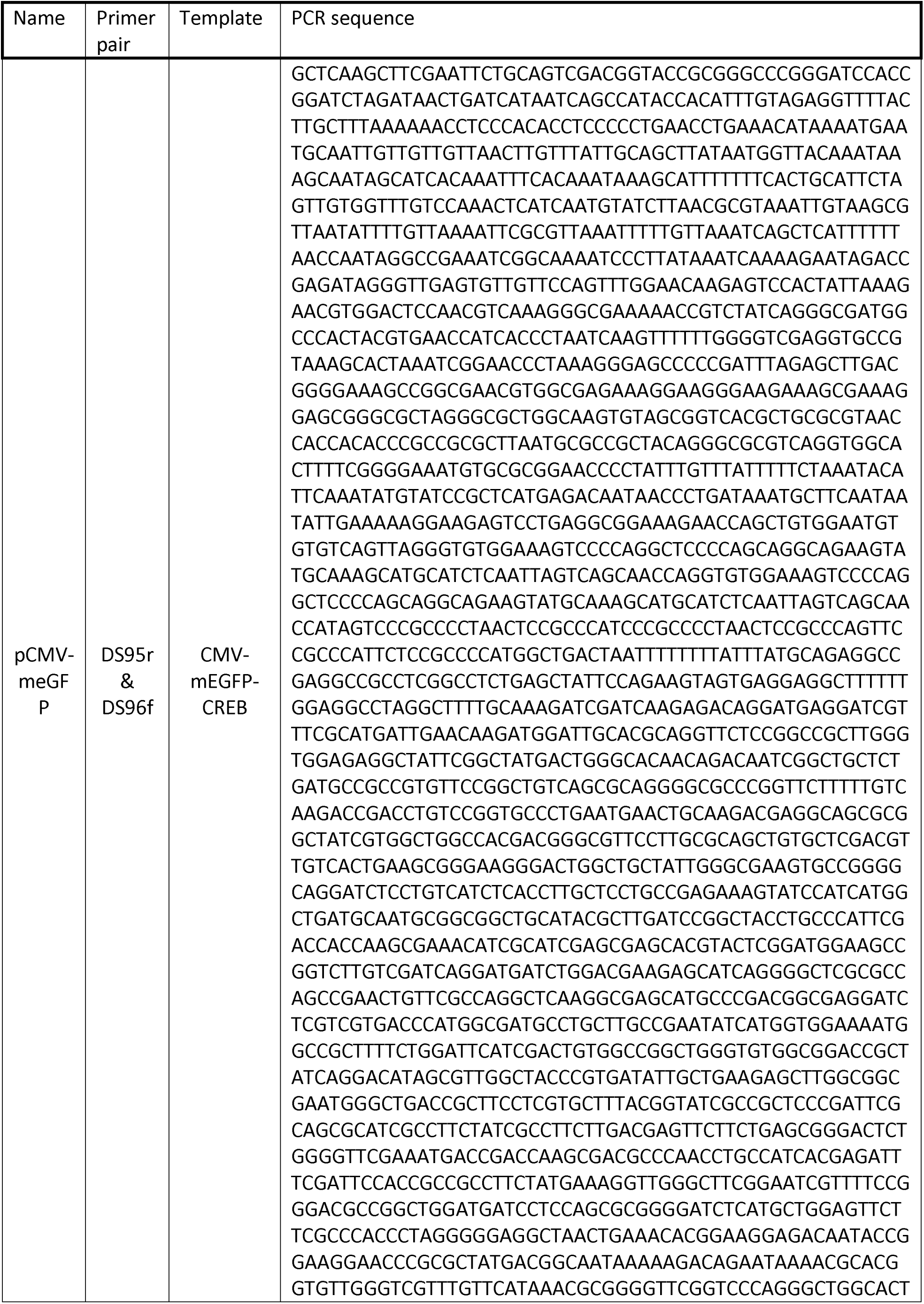
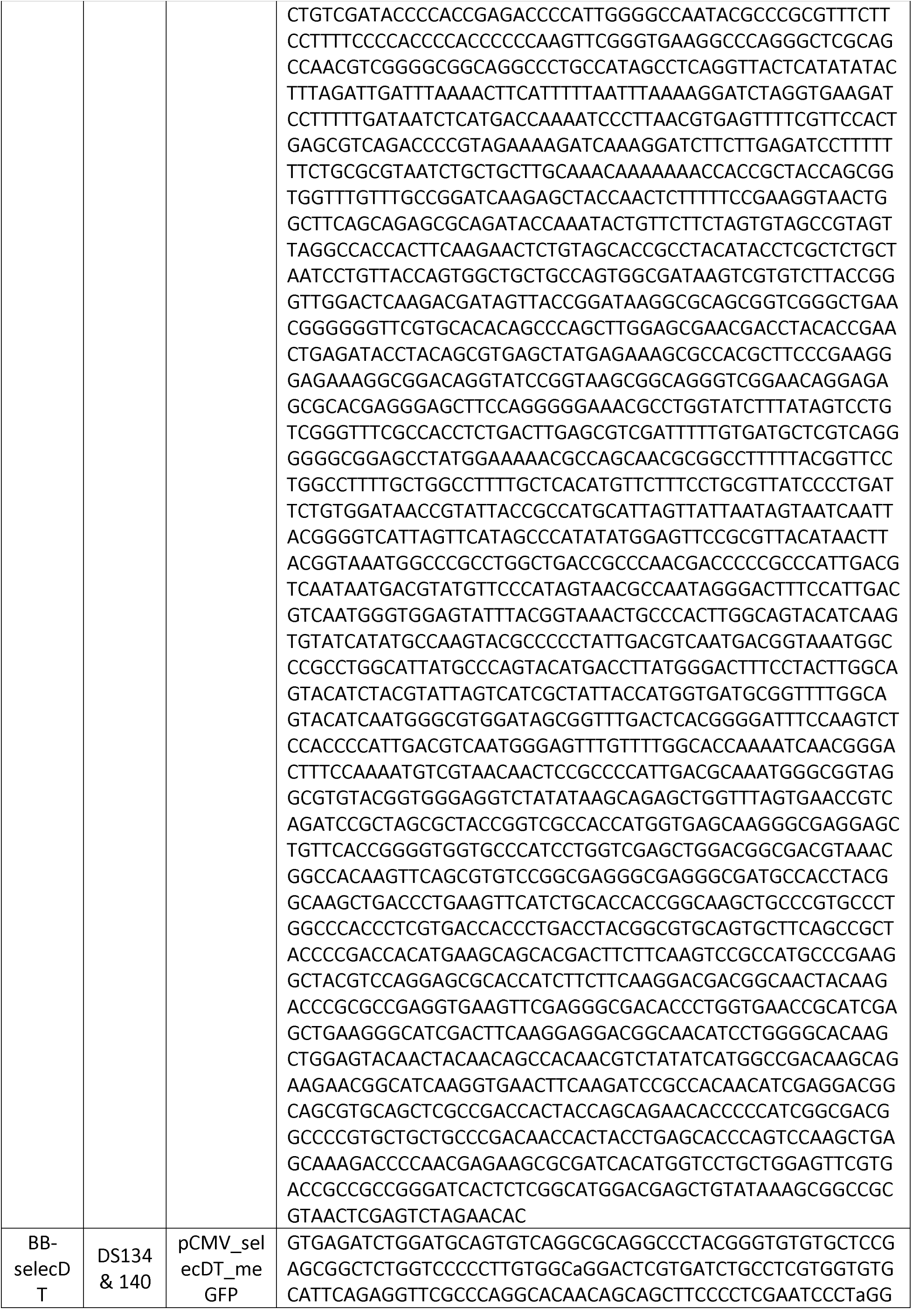
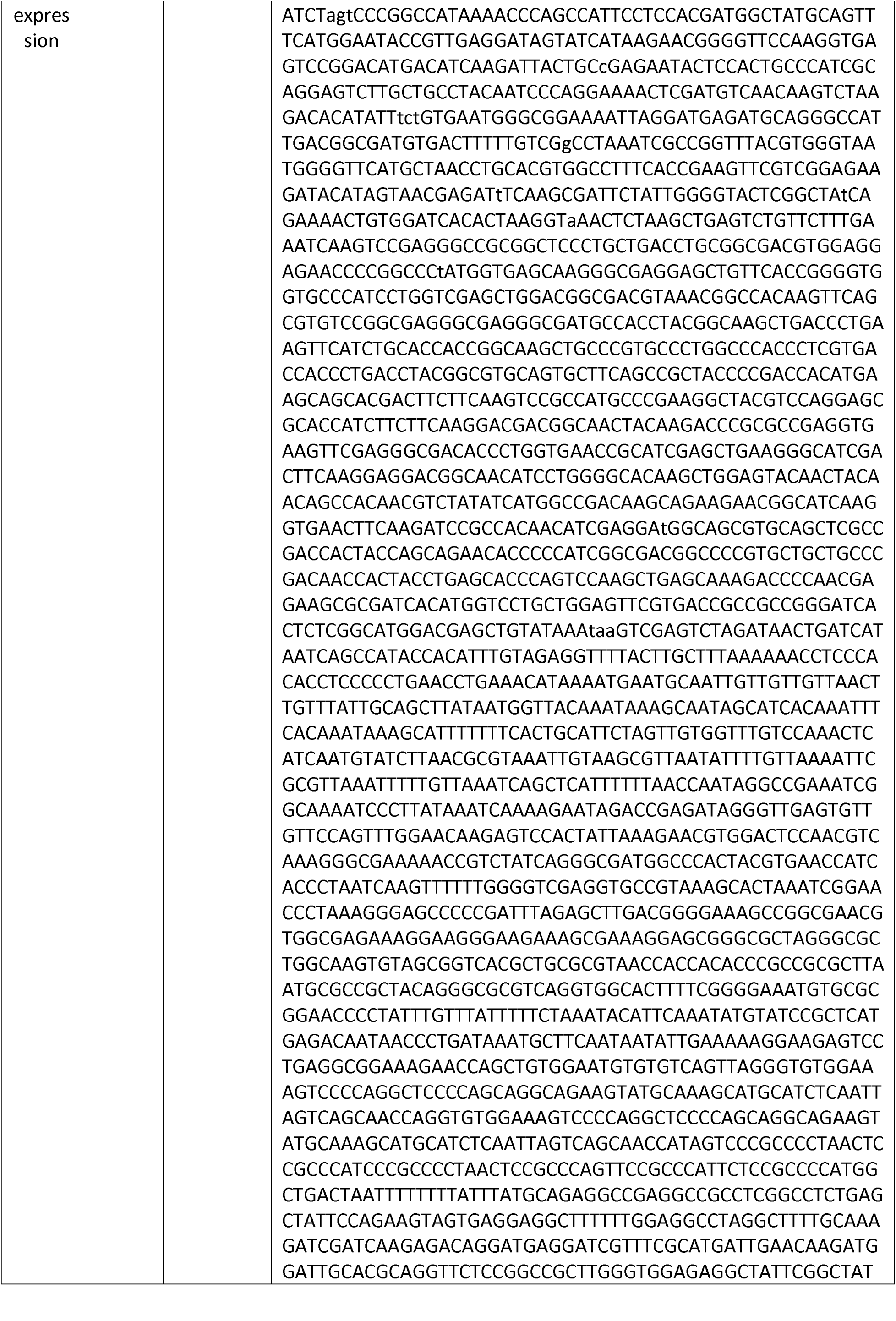
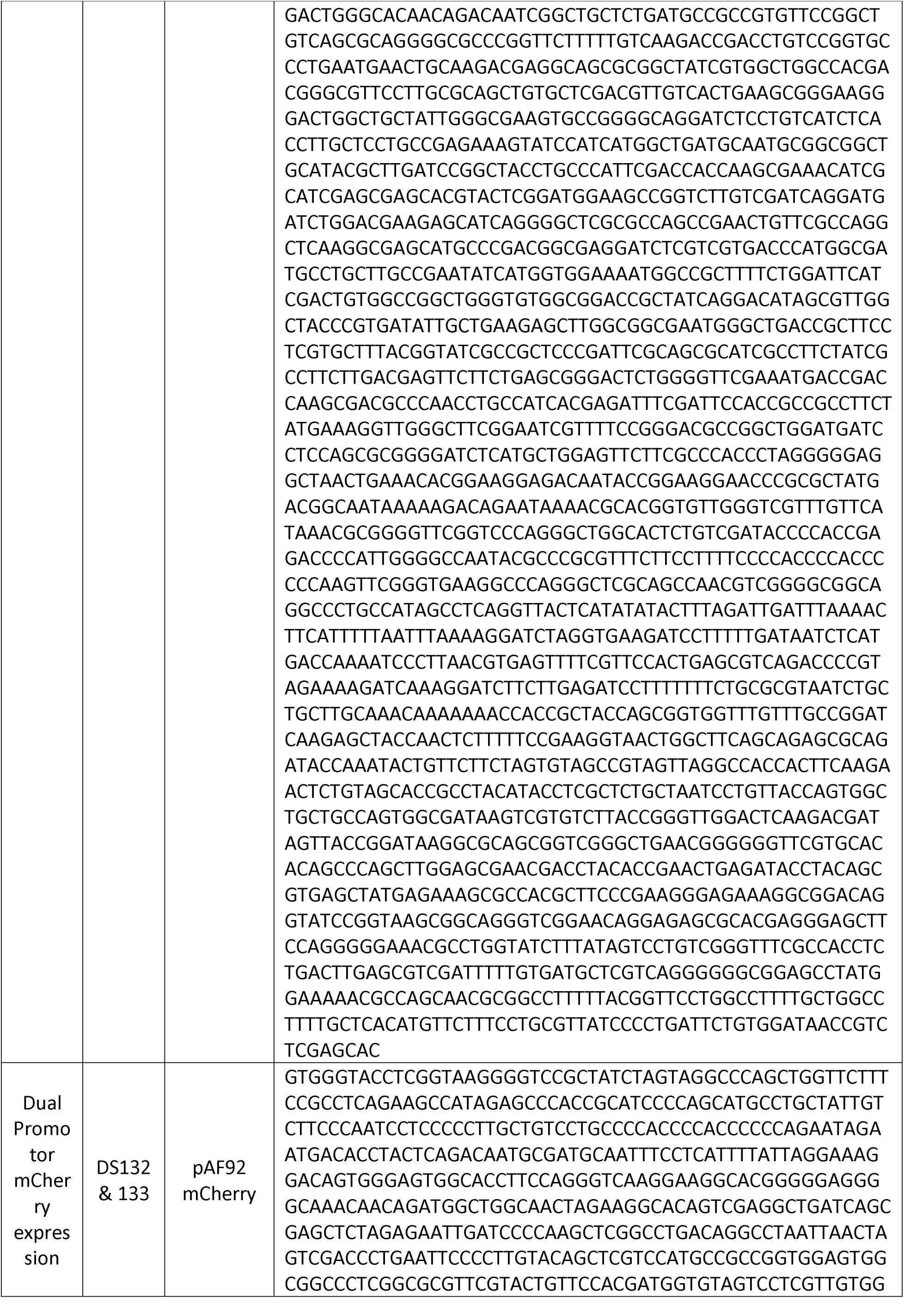
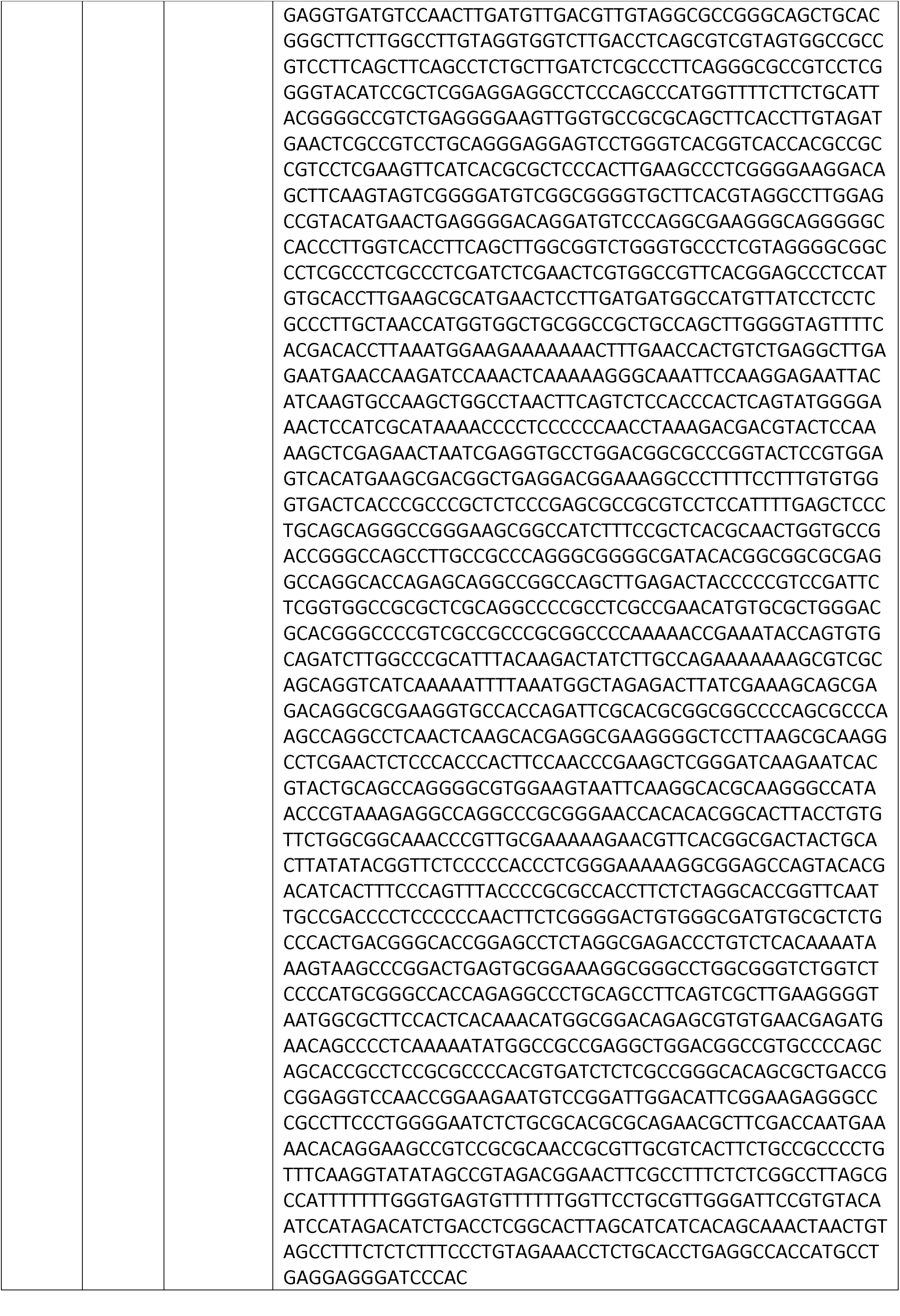
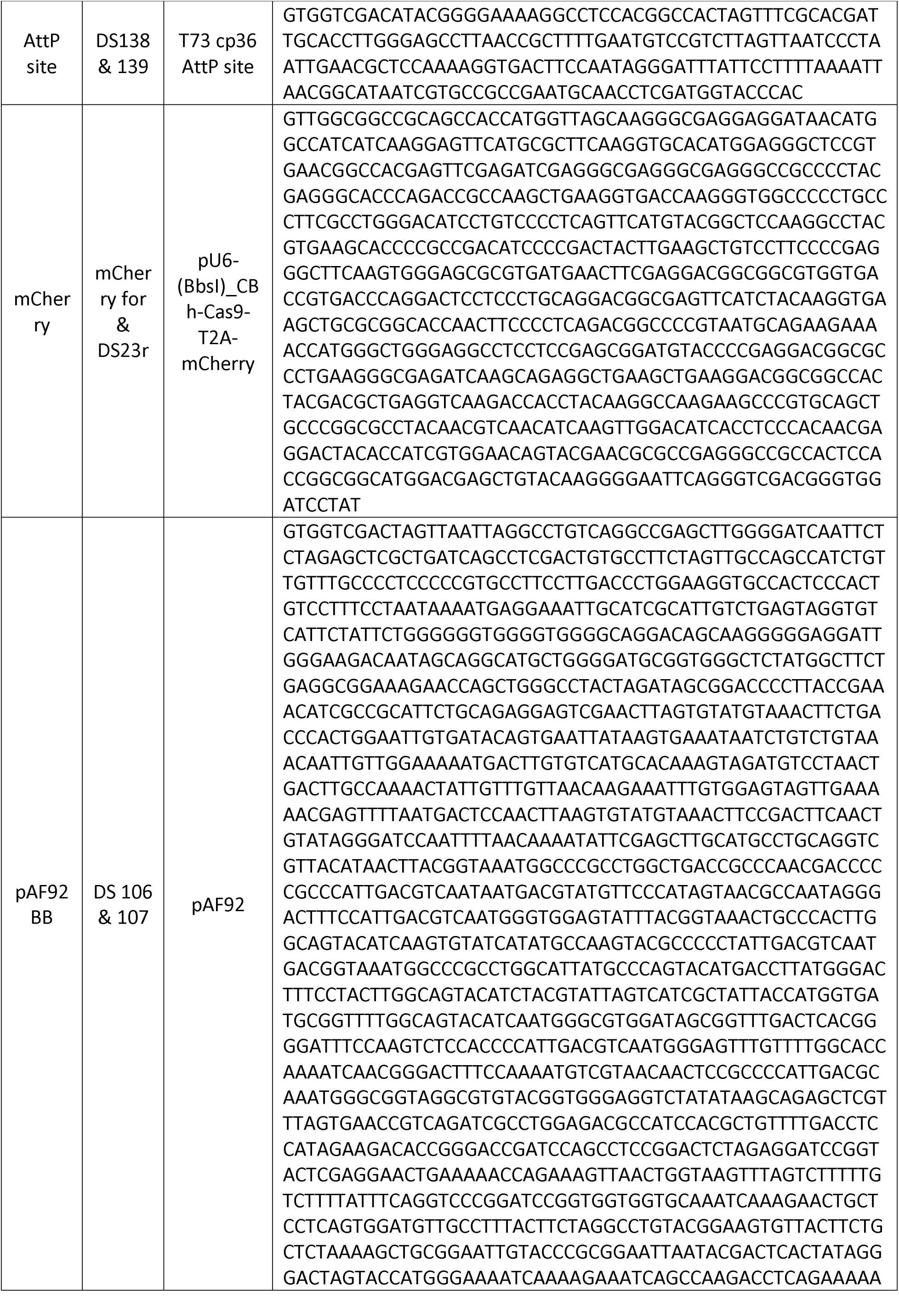
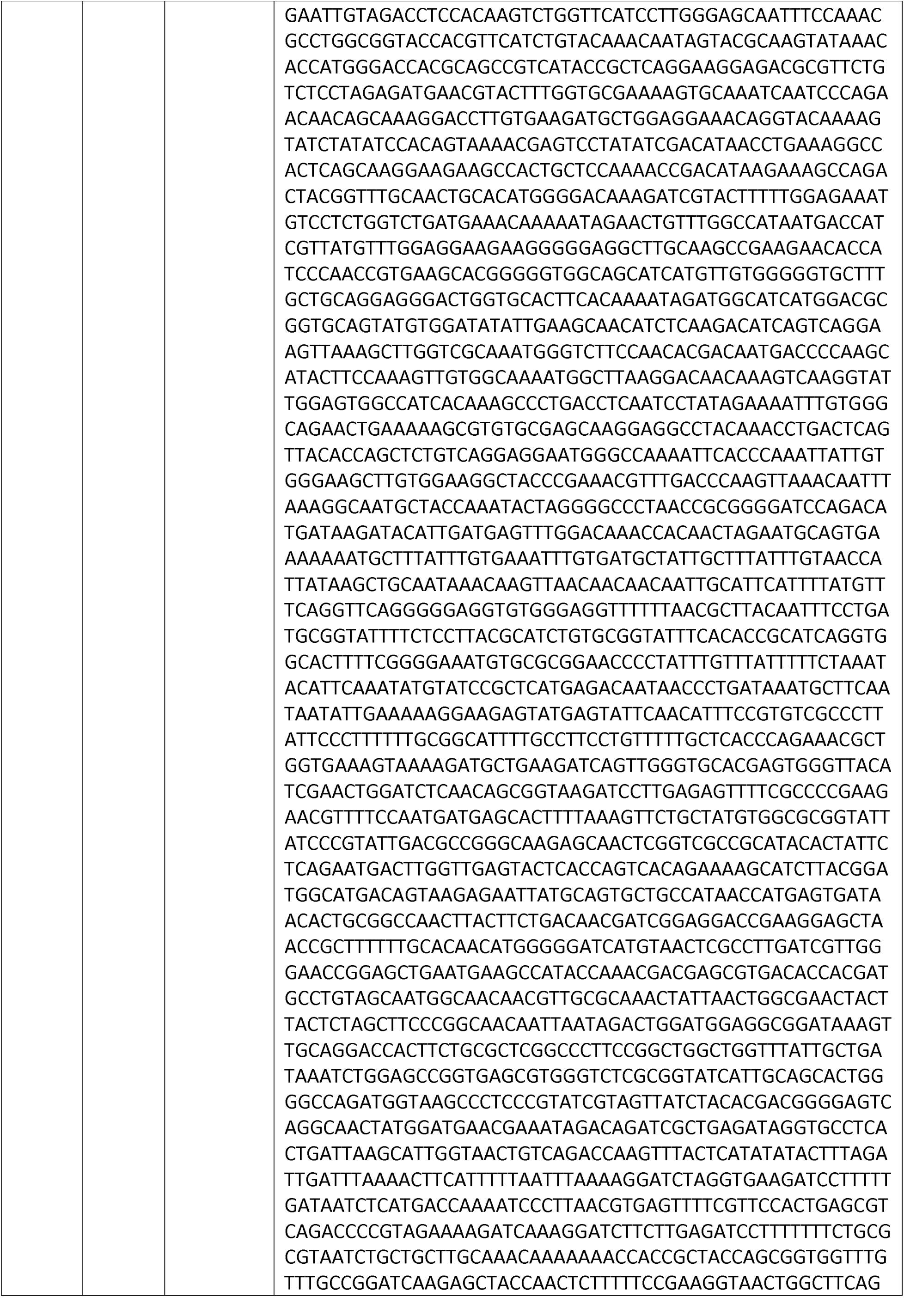
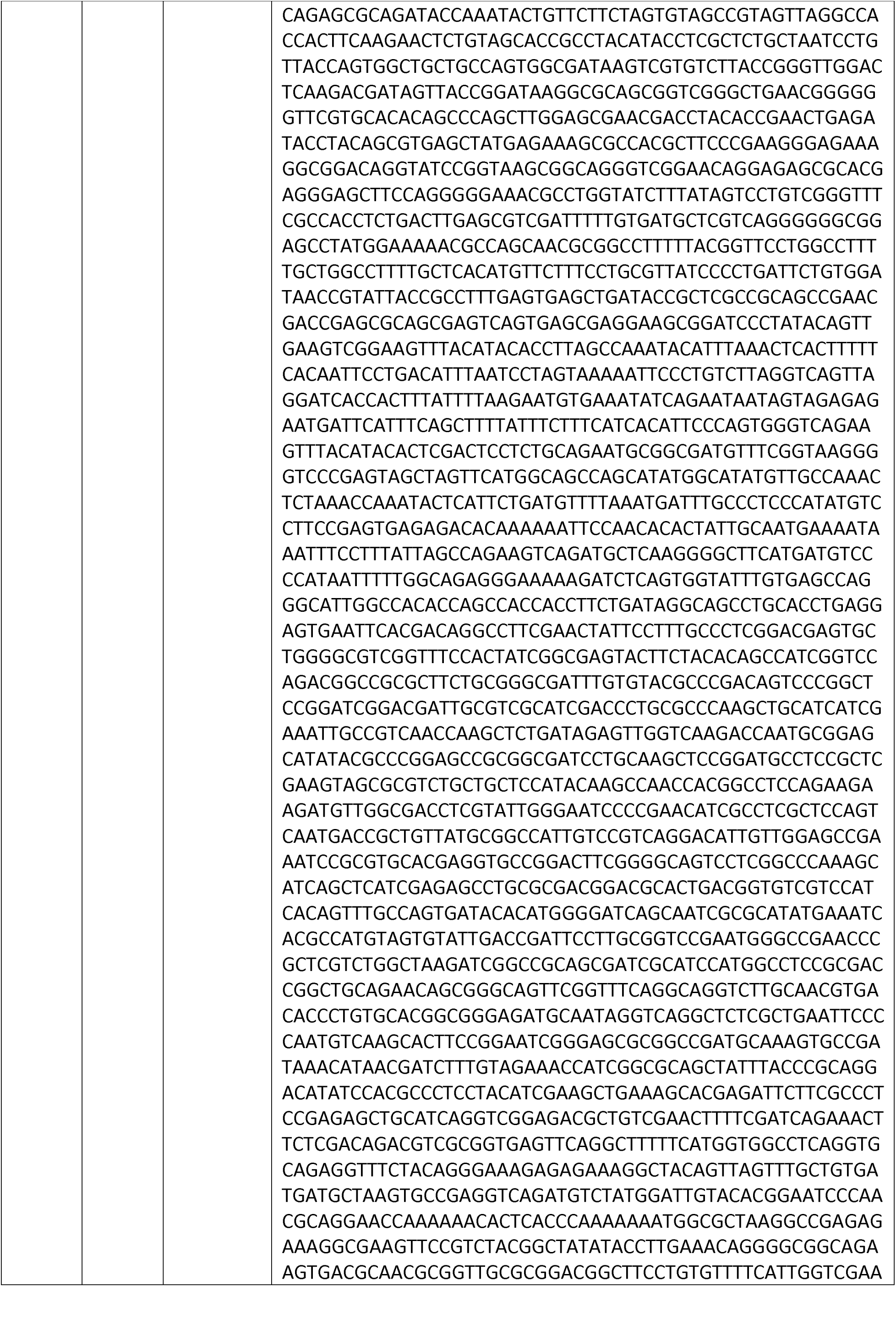
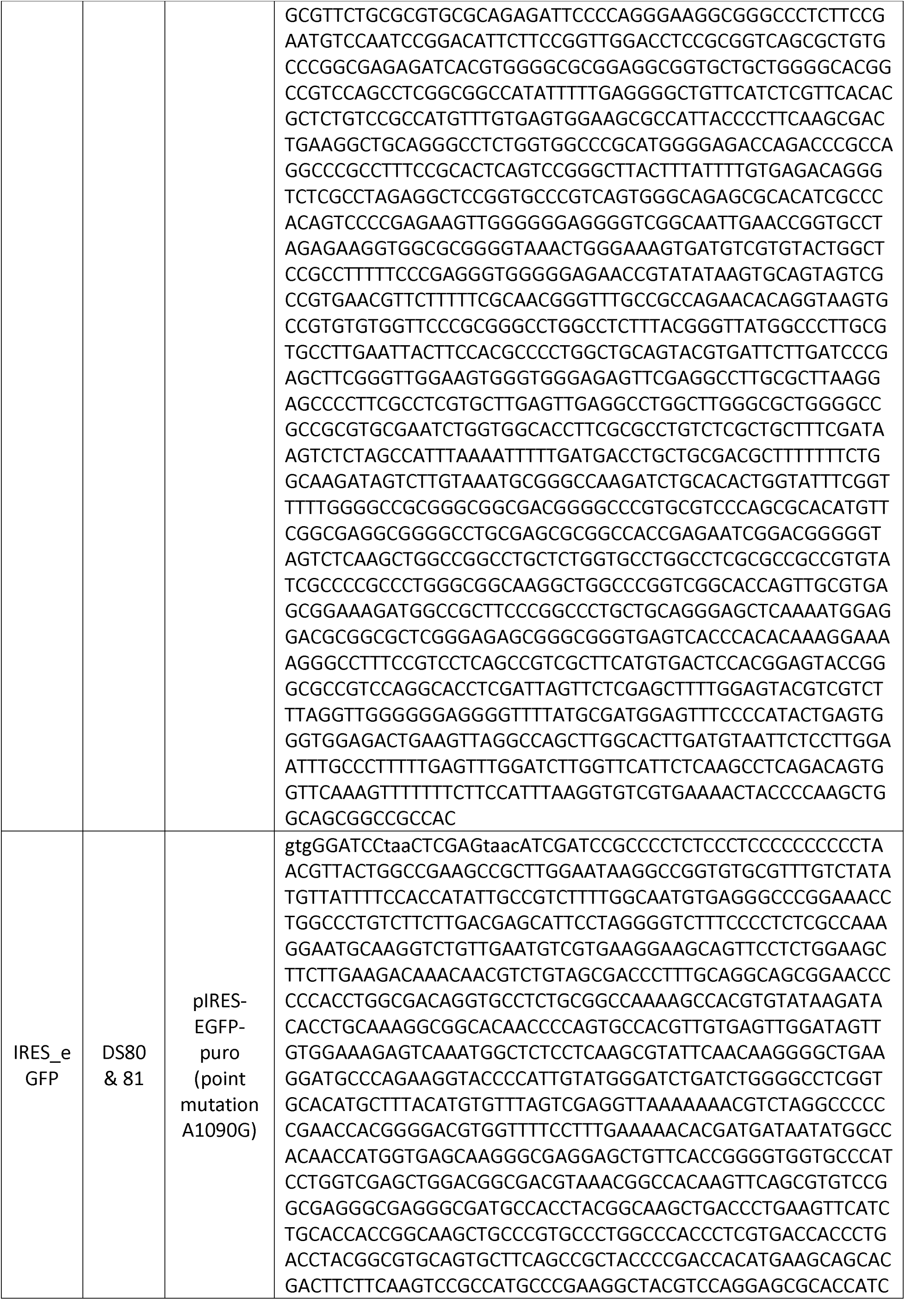
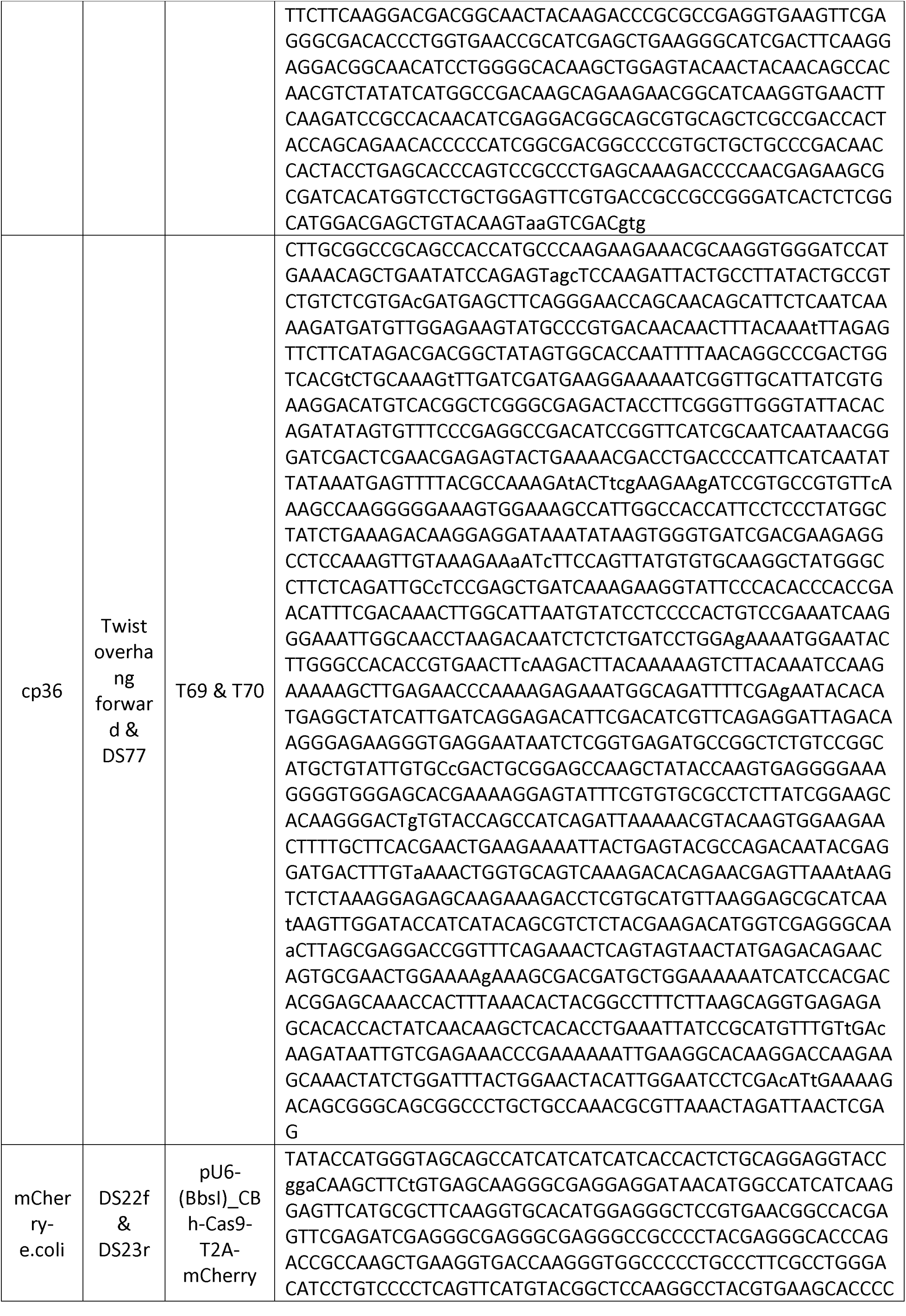
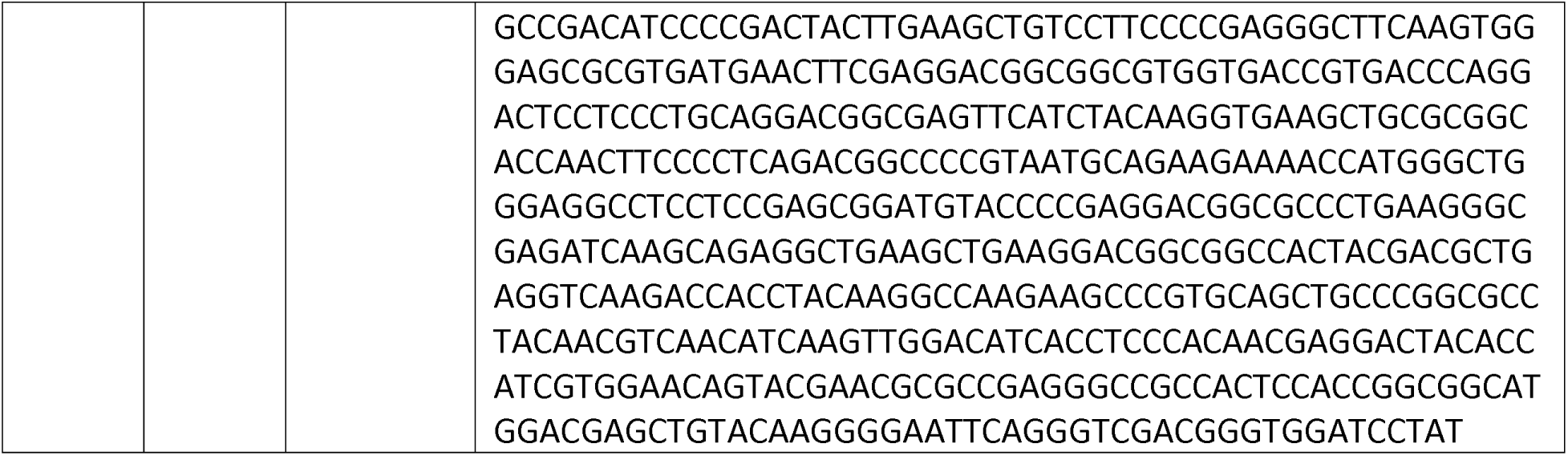
PCR fragment Sequences:

**Table 4:**
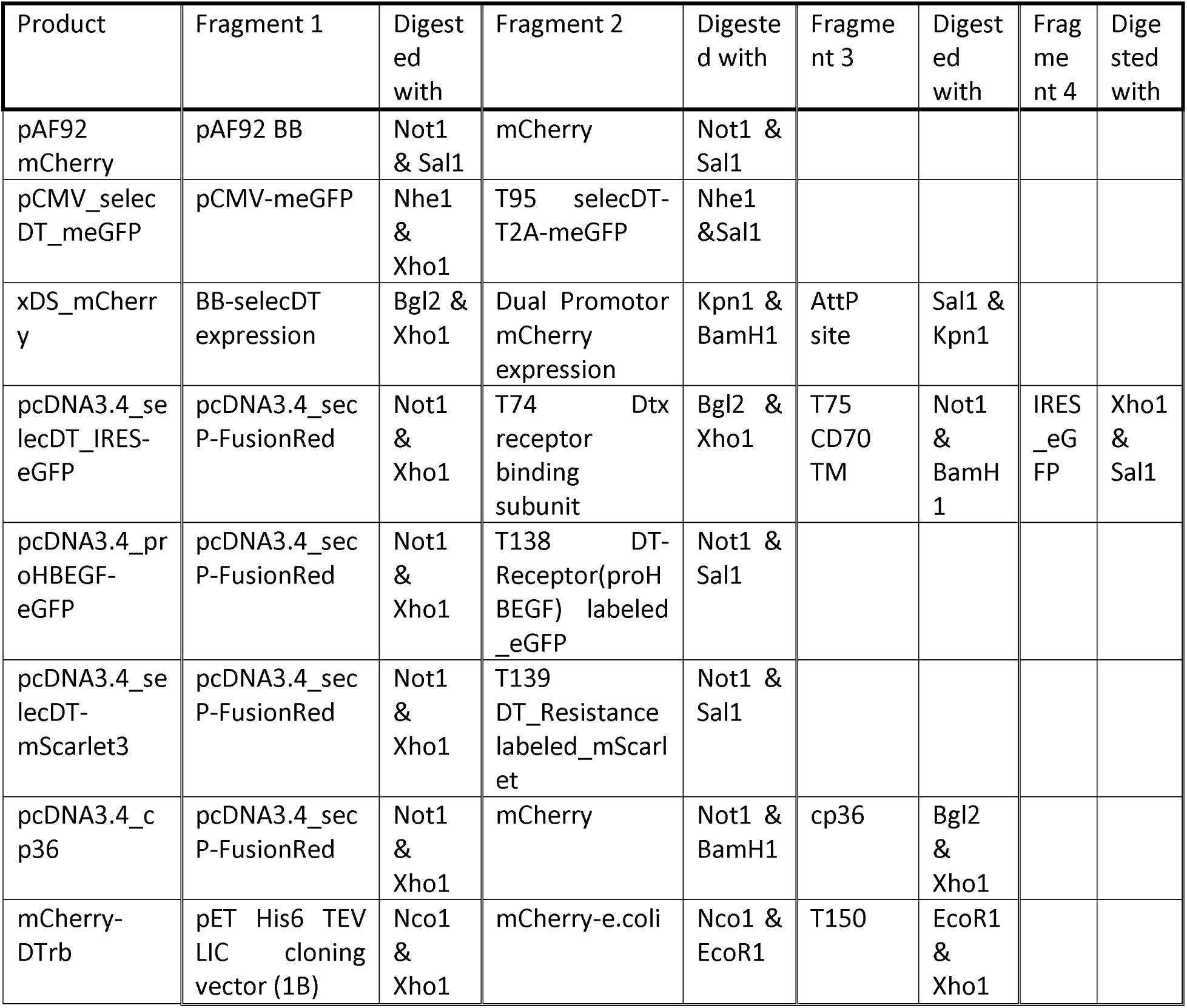
Cloning strategy to obtain final plasmids:

**Table 5:**
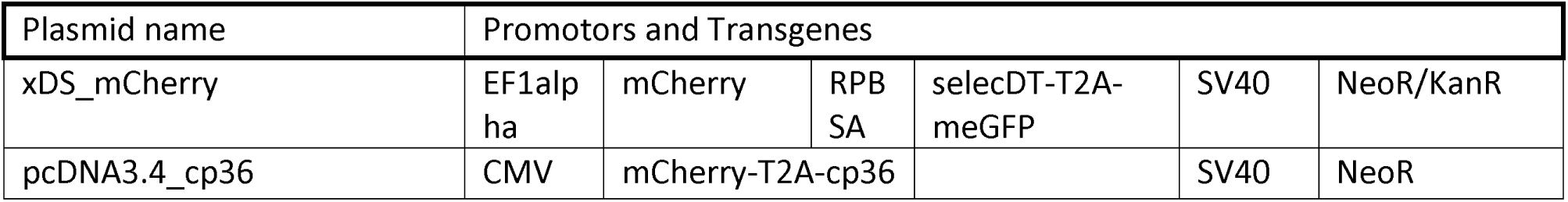
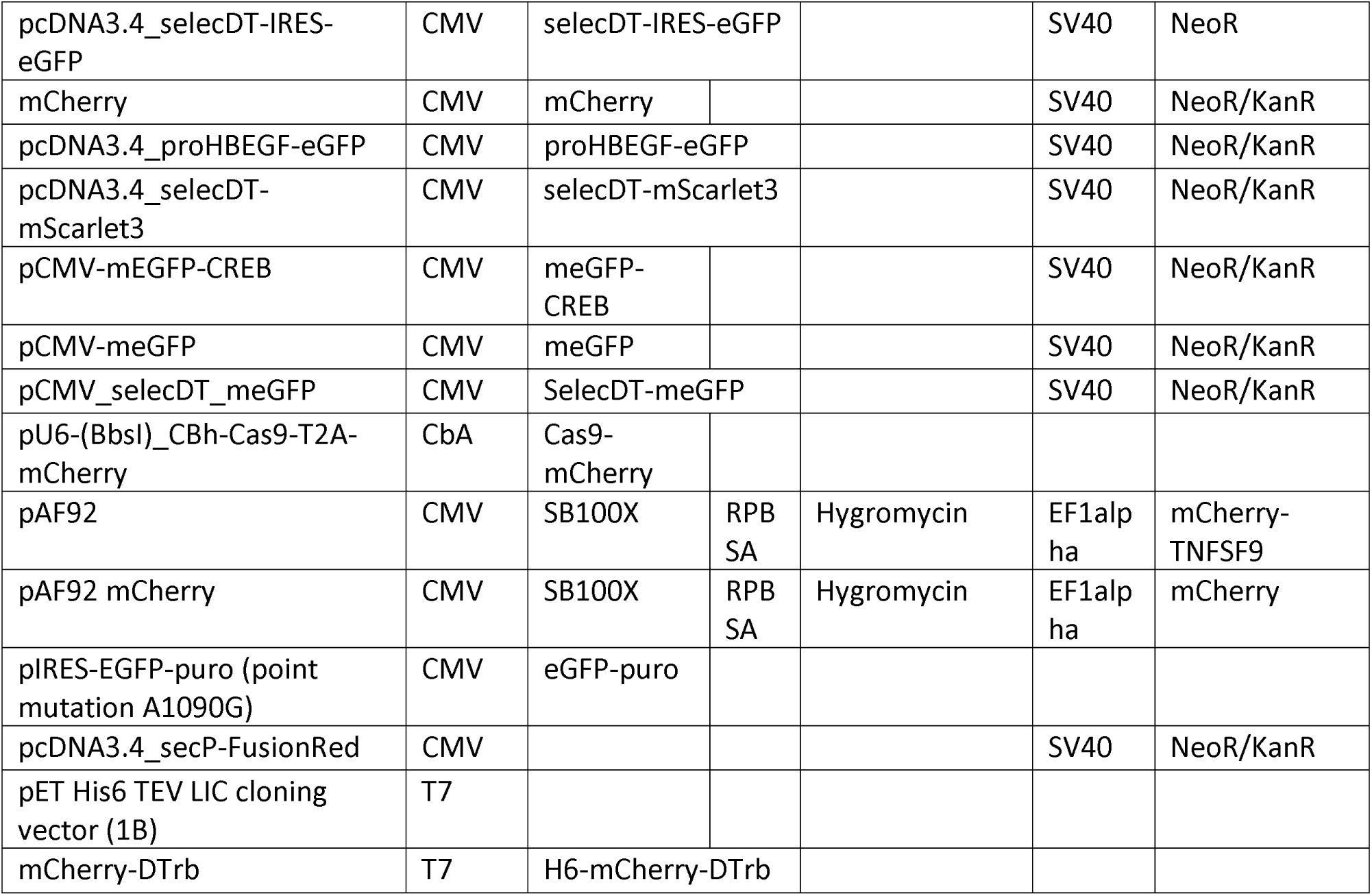
Plasmids used in this study:

### 2. Cell culture

HeLa (ATCC CCL-2) cells were obtained from ATCC (Manassas, VA, USA) and cultivated in Dulbecco’s Modified Eagle Medium (DMEM), high glucose, GlutaMAX supplemented with 10% (v/v) fetal bovine serum (FBS) and 1% (v/v) 100 x Penicillin-Streptomycin (DMEM++) all from Thermo Fisher Scientific (Waltham, MA, USA) at 37°C and 5% CO_2_. The cells were used for experiments between passage 5-30.

Human embryonic kidney (HEK) FreeStyle™ 293-F cells (Thermo Fisher Scientific) were cultured in 30 mL FreeStyle™ medium (Thermo Fisher Scientific) in a 125 mL disposable flask (Jet Bio-Filtration Co., Ltd, Guangzhou, China). The cells were maintained at 37°C and 8% CO_2_ in a HERAcell 240i incubator (Thermo Fisher Scientific) on a CO_2_ resistant cell culture shaker (Thermo Fisher Scientific) at 135 rpm. The cells were maintained at a density between 1.5 x 10^5^ and 3 x 10^6^ cells per mL.

The absence of mycoplasma contamination was regularly checked with the MycoAlert PLUS Mycoplasma Detection Kit (Lonza, Visp, Switzerland).

### 3. Toxin preparation

Unnicked *Corynebacterium diphtheriae* DT was obtained from Calbiochem (San Diego, CA, USA). Activation of DT requires nicking of the amino acid sequence with a protease, while the two resulting fragments remain connected via a disulfide bridge. The required nicking of DT was performed by 1 h incubation of DT at 37°C with 1:1 (w/w) trypsin-EDTA (0.25% w/v) from Thermo Fisher Scientific. The nicked DT was diluted to the appropriate concentration in cell medium and immediately used.

### 4. Cell viability assay (MTS)

Cytotoxicity was assessed *via* the CellTiter 96® Aqueous One Solution Cell Proliferation Assay (Promega Corporation, Madison, WI, USA). Cells were seeded in 96 well plates (Techno Plastic Products AG, Trasadingen, Switzerland) at 2500 cells per well one day prior to toxin exposure. The cells were washed with phosphate buffered saline (PBS) (Gibco™ PBS, pH 7.4, Thermo Fisher Scientific, 37 °C) and 100 µL of maintenance medium containing the indicated DT concentration or 2 mg/mL G418 were added. After 48-h incubation the 3-(4,5-dimethylthiazol-2-yl)-5-(3-carboxymethoxyphenyl)-2-(4-sulfophenyl)-2H-tetrazolium (MTS) assay was performed according to the manufacturer’s protocol. The cells were washed once with 37 °C warm PBS, subsequently 100 µL of medium containing 20% (v/v) MTS reagent was added. After 2 h incubation at 37°C the absorbance at 490 nm was measured for each well in a Spark multimode microplate reader (Tecan Trading AG, Mannedorf, Switzerland). A sigmoidal, 4-parameter logistic curve where X is the concentration of either DT or G418 was fitted to the data in GraphPad Prism version 10 (Graphpad, Boston, MA, USA), and the best fit IC50 values were extracted.

### 5. Transfection

TFAMoplex transfections were performed in 24-well cell culture plates (Techno Plastic Products AG, Trasadingen, Switzerland) with 2,5 x 10^4^ HeLa cells per well as described by Burger et. al^14^. First, the two TFAM fusion proteins with either phospholipase C from *Listeria monocytogenes* (PLC-TFAM) or the human vaccinia related kinase 1 (ccTFAM-VRK1) were mixed into FBS to achieve a final concentration of 0.8 μM of each protein in a final volume of 11 μL. Subsequently, the plasmid was added to obtain a final DNA concentration of 10 ng/μL. The mixture was gently mixed and incubated for 30 min at room temperature. For the integration assays of cp36 (Figure 2A), 10 μL of this mixture were added to the 500 μL DMEM++ in each well. For the selection assay of transiently transfected cells (Figure 1D) the mixture was diluted in FBS to a final volume of 220 μL to achieve a final DNA concentration of 0.5 ng/μL. From this mixture, 10 μL were given to each sample well of a 24-well plate containing 500 μL DMEM++, resulting in 5 ng DNA per well. Before starting the selection protocols, the cells were transfected overnight at 37°C and 5% CO_2_.

HEK cells were transfected with branched PEI of 25 kDa average molecular weight (408727, CAS:9002-98-6) (Sigma Aldrich, St. Louis, MO, USA) at a cell density of 1 x 10^6^ cells/mL with 1 μg of DNA per 1 x 10^6^ cells. To transfect 3 x 10^7^ cells in a volume of 30 mL, 30 μg of plasmid DNA was mixed with FreeStyle™ medium (Thermo Fisher Scientific) to a final volume of 1 mL. Then, 60 μL of PEI (1 μg/μL in PBS) was diluted to a final volume of 1 mL in the same medium. The DNA solution was added to the PEI solution resulting in a final DNA to PEI ratio of 1:2 (w/w). The mixture was vortexed for 20 s, followed by 15 min of incubation at room temperature. Eventually, 2 mL transfection mix was slowly added to the cells while gently swirling the flask. After transfecting for 5 h the cells were spun down in a Sorvall ST 16R centrifuge (Thermo Fisher Scientific) at 50 x *g* for 4 min, the supernatant was discarded and the cells washed once with PBS. The cells were spun down once more and resuspended in fresh FreeStyle™ medium.

### 6. Flow cytometry

To quantify transfection efficiency, cells were analyzed via flow cytometry for expression of a fluorescent marker protein (eGFP or mCherry). HeLa cells in a 24-well plate were washed three times with 500 μL 37°C warm PBS and detached with 200 μL of trypsin-EDTA (0.05% w/v) (Thermo Fisher Scientific). The detached cells were transferred into a 96-well U-bottom plate (Greiner Bio-One, Kremsmünster, Austria) and supplemented with 20 μL FBS per well to inactivate the trypsin. Analogously, the HEK cells were transferred immediately from their culture flask to the U-Bottom plate. The plate was spun down at 300 xg for 1 min, the supernatant was removed and the cell pellets were resuspended in 125 μL ice cold PBS containing 1% (w/v) BSA and 1 mM EDTA. Flow cytometry analysis was subsequently performed in a CytoFLEX Flow Cytometer (Beckman Coulter Life Sciences, Nyon, Switzerland), eGFP fluorescence was measured with the 488/6 nm excitation laser combined with the 525/40 emission filter, while mCherry was measured at 561/10 excitation and 610/20 emission. For each well 10,000 cells were analyzed, while each experiment was performed in three technical replicates. The obtained cell populations were analyzed using the FlowJo software (Tree Star Inc., Ashland, OR, USA). After gating for single cells, the transfection efficacy was determined by setting the gate for the negative control (untransfected cells) to 1% GFP-positive cells and applying this gate to all samples.

### 7. Microscopy

HeLa cells were seeded in µ-Slide 8 Well Glass Bottom (Ibidi, Gräfelfing, Germany) microscopy slides at a density of 25,000 cells per well. The next day, the cells were transfected with 25 ng plasmid DNA per well using the TFAMoplex system as described above. One day later the cells were incubated with mCherry-DT receptor binding probe at 10 μM for 2 h. After washing 3 times with PBS the cells were fixed with 4% paraformaldehyde at pH 7.4 for 20 min. After washing the fixed cells 3 times with PBS they were stained with Hoechst 33342 (2.5 µg/mL) (Sigma Aldrich, St. Louis, MO, USA) for 20 min and subsequently washed again twice with PBS. The cells were stored in PBS for not more than 3 days at 4°C until they were imaged via confocal microscopy (Nikon Eclipse Ti2 (inverse) with a Yokogawa Confocal Scanner Unit CSU-W1-T2 SoRa, and a sCMOS Hamamtasu Orca Fusion BT camera (Nikon, Tokyo, Japan)). The cells were imaged in three different channels with all lasers set to 20% intensity, the Hoechst channel was excited at 405 nm and the fluorescent emission was collected through a 447 nm filter with a bandwidth of 60 nm. Correspondingly, eGFP was excited with a 488 nm laser and detected using a 525 nm filter with a bandwidth of 50 nm, and mCherry was excited with the 561 nm laser and emission was detected through a 610 nm filter.

### 8. Expression and purification of recombinant mCherry-DT-receptor binding subunit (mCherry- DTrb)

The plasmid encoding for the His-tagged mCherry-DTrb protein was transformed into SHuffle® T7 Express lysY Competent E. coli (NEB, Ipswich, MA). The next day, a single colony was picked and grown at 37 °C and 210 rpm in an orbital shaker (Infors AG, Bottmingen, Switzerland) in 8 mL of LB overnight (Lab Logistics Group, Meckenheim, Germany). The overnight culture (7 mL) was used to inoculate 700 mL LB medium containing 50 mg/L kanamycin (Sigma Aldrich, St. Louis, MO, USA) and 0.2% (w/v) glucose (Sigma Aldrich, St. Louis, MO, USA), the bacteria were left to grow at 37°C and 210 rpm until they reached an optical density of 0.6 at 600 nm absorbance. The bacteria were cooled to 24°C before protein expression was induced by addition of Isopropyl-β-D-thiogalactopyranosid (IPTG) (AppliChem GmbH, Darmstadt, Germany) to a final concentration of 0.4 mM. After overnight expression at 24°C and 210 rpm in an orbital shaker, the cells were pelleted by centrifugation (5000 x *g*, 10 min at 4 °C, Sorvall LYNX 6000 centrifuge, Thermo Fisher Scientific). The pellet was resuspended in lysis buffer (1 M KCl, 0,5 x PBS pH 7.4, 1 x Halt™ protease inhibitor cocktail (Thermo Fisher Scientific), 1 mg/mL lysozyme (AppliChem GmbH, Darmstadt, Germany)) and lysed using a Fisherbrand™ Q705 probe sonicator (Thermo Fisher Scientific) on ice with a pulse time of 5 s followed by a 10 s pause for a total sonication time of 2 min. PEI was added to the lysate to a final concentration of 0.1% (w/v) followed by centrifugation at 30,000 x g for 45 min at 4 °C to remove cell debris and DNA. To a 14 ml Protino column (Macherey-Nagel GmbH, Düren, Germany) 1 mL of Ni-NTA Agarose resin (Qiagen, Hilden, Germany) was added and equilibrated with wash buffer (1 M KCl, 0,5x PBS pH 7.4, 25 mM imidazole (AppliChem GmbH, Darmstadt, Germany)). To this column the filtered supernatant supplemented with 20 mM imidazole was added. The column was subsequently washed with 20 CV of wash buffer and finally eluted with 4 mL elution buffer (1 M KCl, 0.5x PBS pH 7.4, 250 mM imidazole). The buffer was exchanged to storage buffer (0.5x PBS pH 7.4, 20% (v/v) glycerol) by three 10x steps in an Amicon Ultra 15 Centrifugal Filter (Merck, Darmstadt, Germany). The final protein was concentrated to a final volume of 1 mL and the protein concentration was estimated by spectrophotometry at 280 nm (NanoPhotometer Pearl, Implen GmbH, Munich, Germany). The sample was aliquoted and snap frozen in liquid nitrogen. Protein size and purity was estimated by sodium dodecyl sulfate polyacrylamide gel electrophoresis (SDS-PAGE) performed with 5 µg protein sample diluted with 3 µL Laemmli buffer (4% (w/v) SDS, 20% (v/v) glycerol, 10% (w/v) 2-mercaptoethanol, 0.004% (w/v) bromophenol blue, 125 mM Tris-HCl) (Sigma Aldrich, St. Louis, MO, USA). Protein samples were heated for 3 min at 95°C and loaded on a 12-well Mini-PROTEAN TGX Stain-Free Gel (Bio-Rad Laboratories, Inc, Hercules, CA, USA). SDS-PAGE followed by densiometric analysis using the software ImageJ^32^.

### 9. Statistical analysis

Statistical analysis of the obtained flow cytometry and viability data was performed using the GraphPad Prism software version 10 (Graphpad, Boston, MA, USA). Data are presented as the mean ± standard deviation. The data are derived from at least three independent experiments, if not stated otherwise each performed as technical triplicates. ANOVA followed by Tukey’s multiple comparisons method was used for significance testing.

## Acknowledgements

This project has received funding from the European Research Council (ERC) under the European Union’s Horizon 2020 research and innovation program (grant agreement No 884505).

The authors gratefully acknowledge ScopeM (ETH Zurich) for their support and assistance in this work.

## Author Contributions

D.S.: Conceived the idea for SelecDT, designed and performed all experiments, primary author of the manuscript.

S.H.: Helped performing some experiments and gave intellectual input for the experimental design J.-C.L.: Manuscript revision and general supervision.

M.B.: Contributed to experimental design, data evaluation, manuscript writing and revisions.

The authors do not declare any competing interests.

## Supplemental Information

Cellular localization of recombinantly tagged version of the receptor and resistance involved in selecDT:

**Figure 1:**
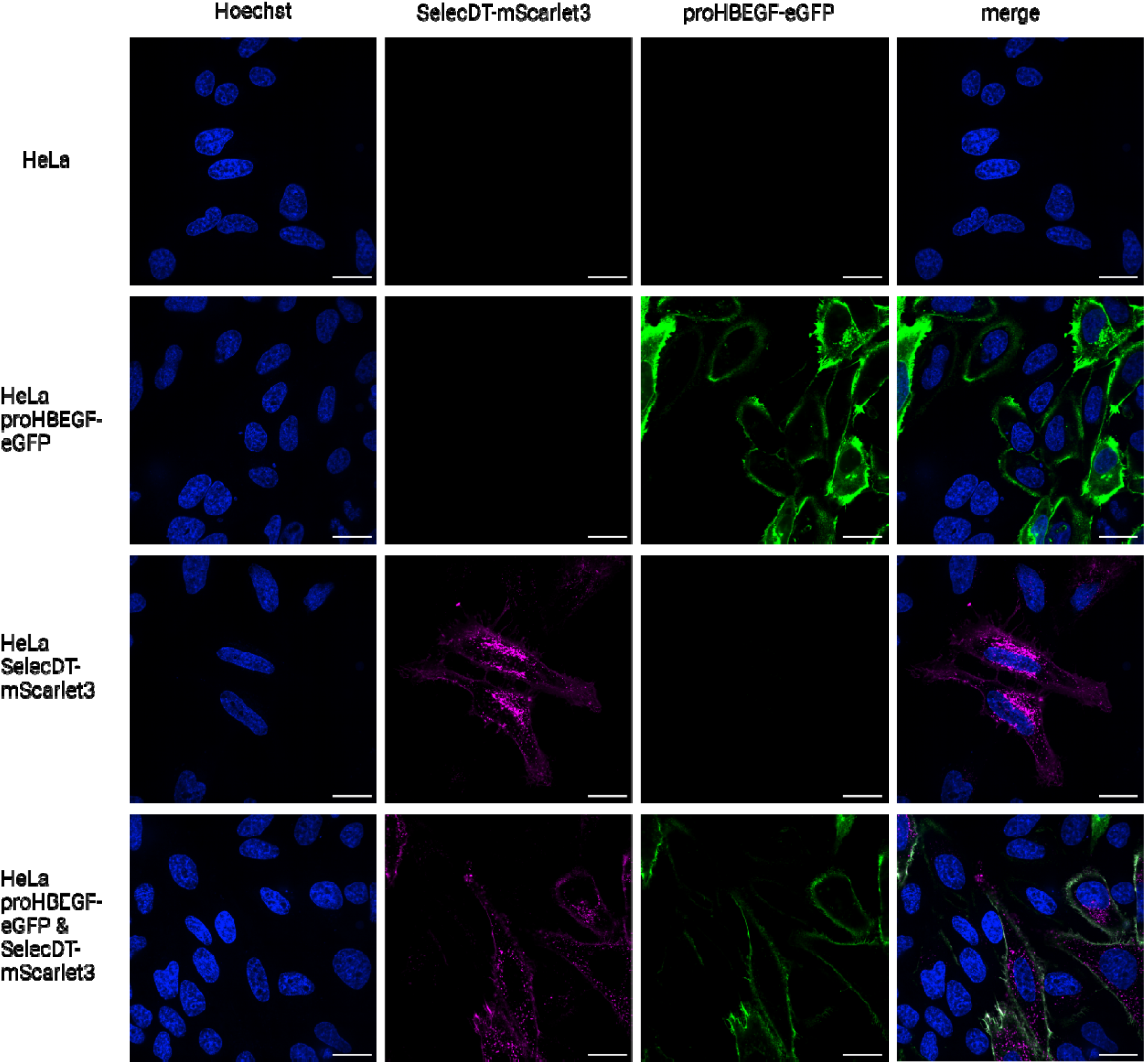
confocal microscopy images of a single slice in the middle of each cell. Untreated HeLa cells are shown i the top panel (slice 16 of 45), the second panel depicts cells overexpressing proHBEGF-eGFP (DT-receptor) (green) (slice 10 of 34). In the third panel, selecDT was C-terminally labeled with an mScarlet3 fluorophore (magenta) (slice 15 of 45), while the bottom panel shows coexpression of both proHBEGF-eGFP and selecDT-mScarlet3 (slice 14 o f 41). The indicated scale bars represent 25 µm. All images were acquired using identical microscope and analysis settings.

Cells transfected on the same day as cells in figure 2A, B, C, treated and imaged identically except for the omission of mCherry-DTrb.

**Figure 2:**
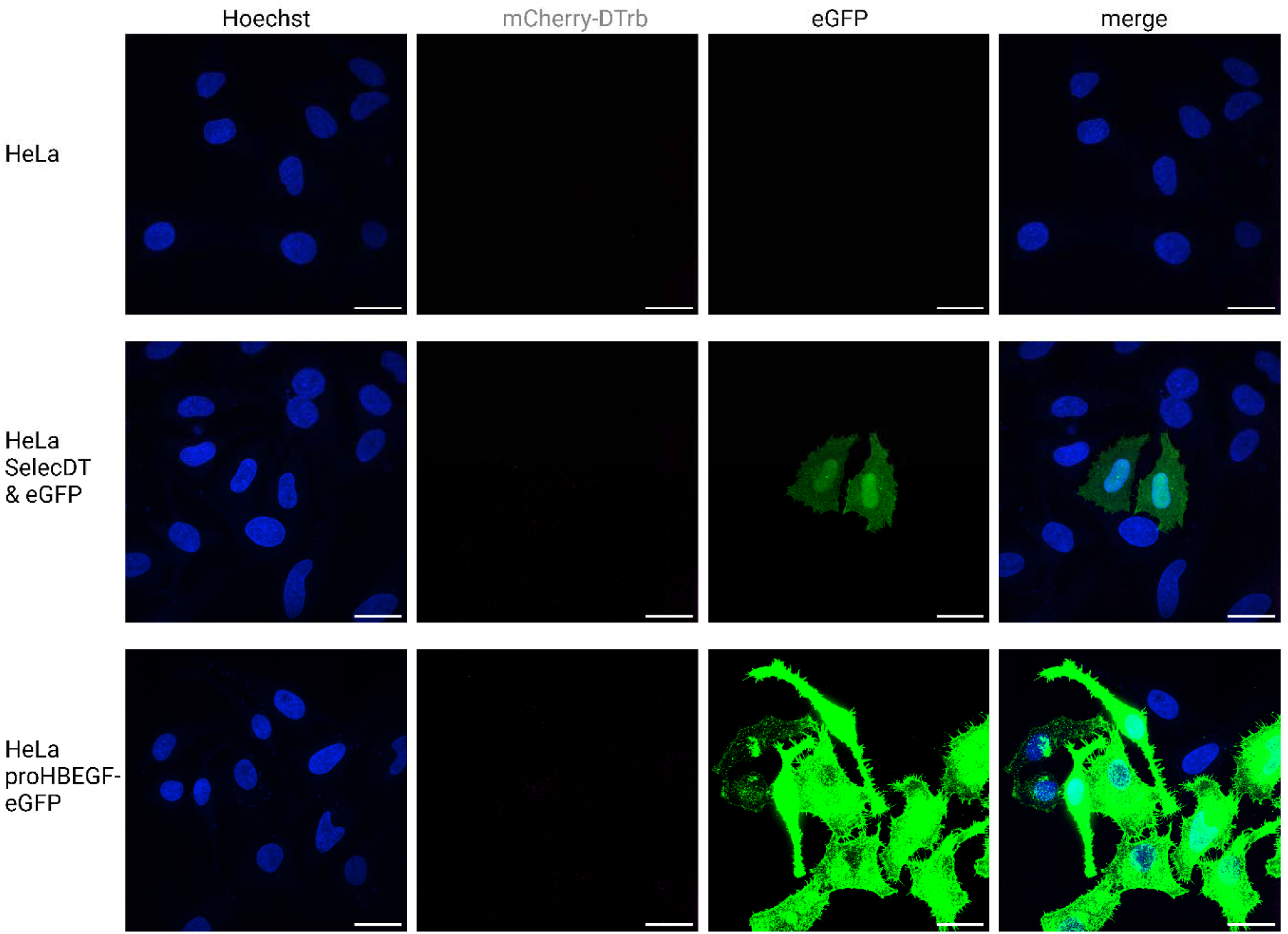
confocal microscopy images of control cells for Figure 2A,B,C without addition of mCherry labeled DTr (magenta) the cells nuclei were stained with Hoechst (blue). The images were acquired in 200 nm-spaced slices throughout the height of the cells, subsequently the resulting slice stacks were projected to depict the maximal value for each pixel (Z-projection). Naïve HeLa cells are shown in the top panel, the middle panel depicts selecD cells coexpressing soluble eGFP (green), while the bottom panel shows cells overexpressing proHBEGF-eGFP (DT-receptor) (green). The indicated scale bars represent 25 µm. All images were acquired using identical microscope and analysis settings, identical to the settings used for Figure 2 & Supplemental Information Fig. 3.

The images for figure 2A, B, C were acquired in 200 nm-spaced slices throughout the height of the cells, subsequently the resulting slice stacks were projected to depict the maximal value for each pixel (Z-projection).

**Figure 3:**
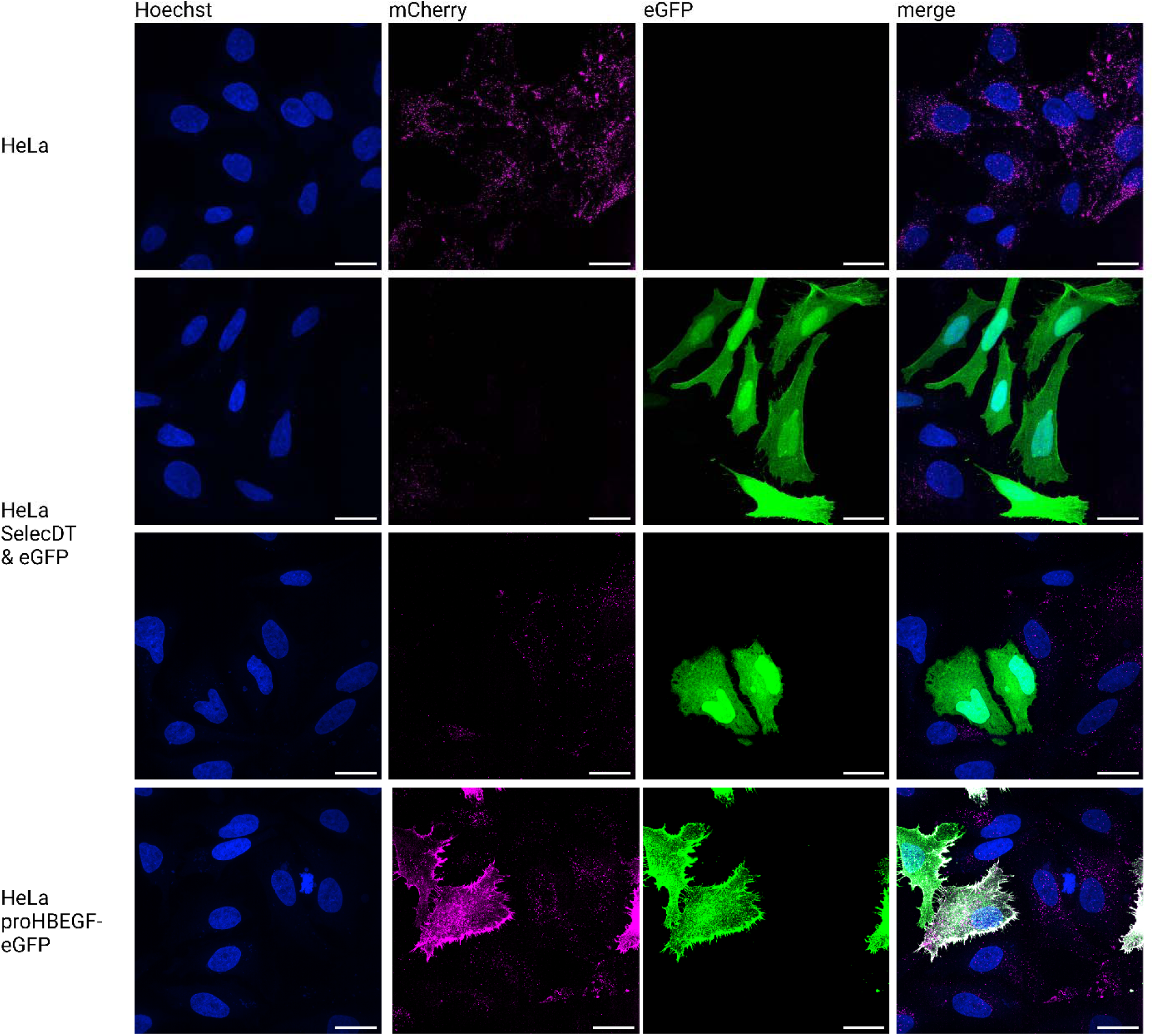
Z-projection of the confocal microscopy images of the cells depicted in Figure 2A,B,C after incubation with 10 nM mCherry labeled DTrb (magenta) the cells nuclei were stained with Hoechst (blue). The images were acquired in 200 nm-spaced slices throughout the height of the cells, subsequently the resulting slice stacks were projected to depict the maximal value for each pixel (Z-projection). Naïve HeLa cells are shown in the top panel, the middle panels depicts selecDT cells coexpressing soluble eGFP (green), while the bottom panel shows cells overexpressing proHBEGF-eGFP (DT-receptor) (green). The indicated scale bars represent 25 µm. All images were acquired using identical microscope and analysis settings, identical to the settings used for Figure 2 & Supplemental Information Fig. 2.

Hela cells and neoR expressing HeLa cells subjected to increasing amounts of G418 for one day:

**Figure 4:**
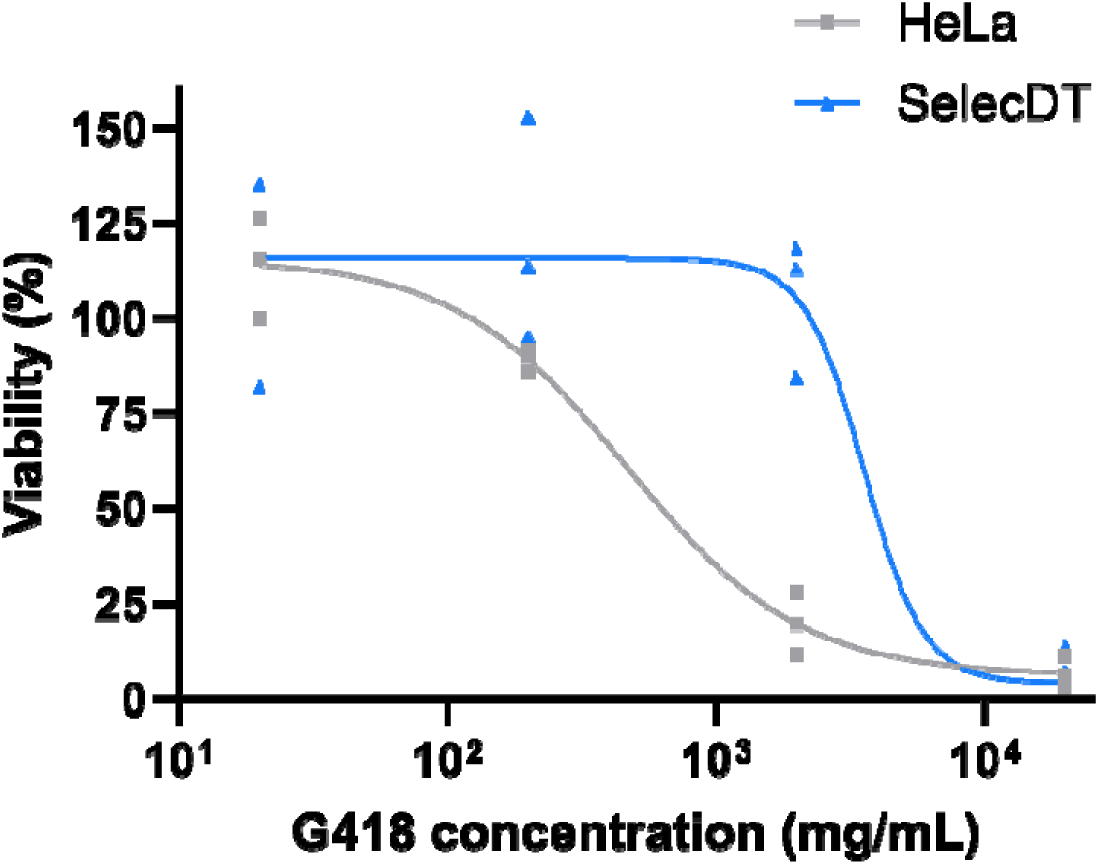
Viability of naïve and neoR HeLa cells upon G418 titration measured via MTS. The data plotted is the mean of the three independent biological replicates each conducted in 3 technical replicates (n=3) The fitting curve indicates an IC50 value of naïve HeLa cells at 466 μg/mL G418. Cells protected by neoR survived higher doses of G418 and the fitted curve indicated an IC50 value of 3559 μg/mL.

HeLa cells transiently transfected to a higher initial transfection rate selected for one day with DT:

**Figure 5:**
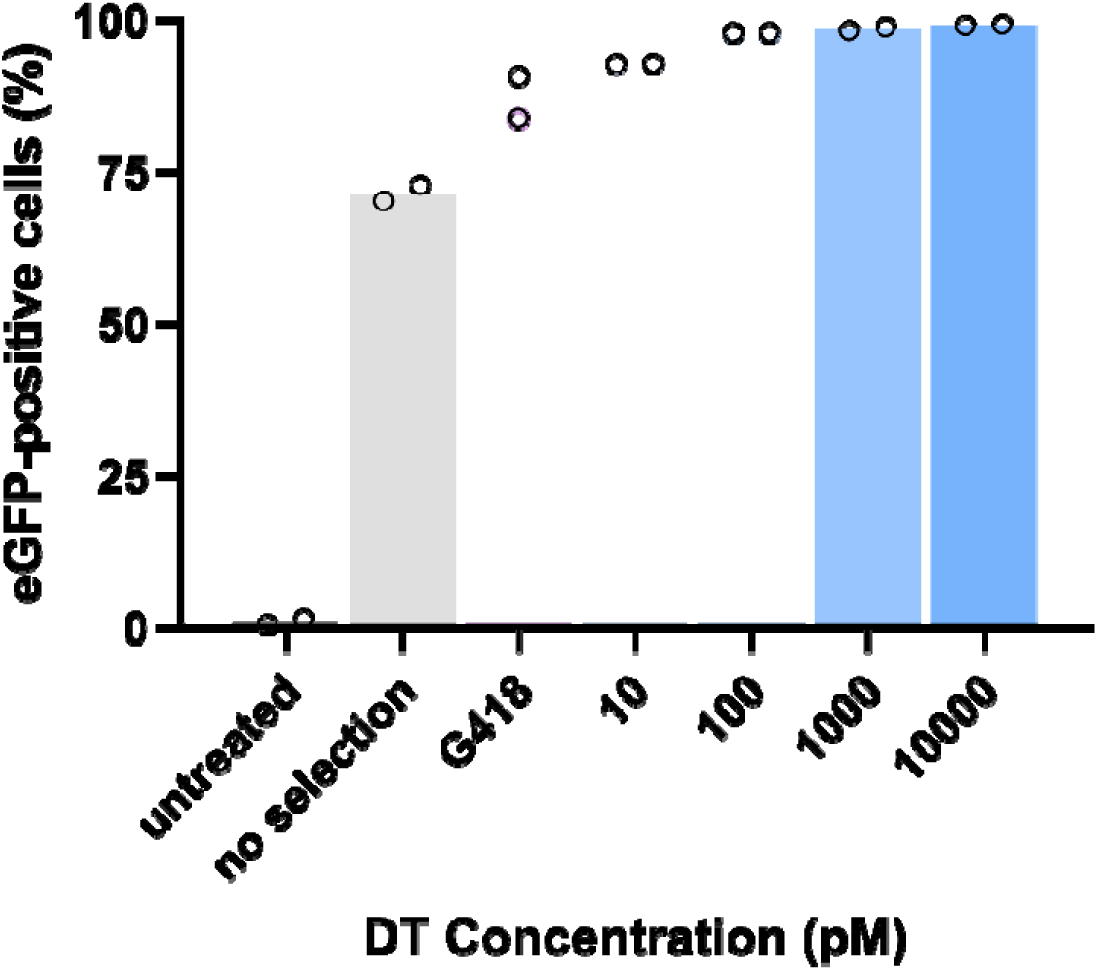
Flow cytometry analysis of HeLa cells coexpressing a marker transgene (eGFP) and both resistance genes SelecDT and neoR, selected with G418 (magenta), DT (blue) or mock (gray) after one day of selection. Cells were transfected with 20 ng DNA per well to achieve higher initial transfection efficacy. Y-axis: GFP-positive cells (%); X-axis: selection conditions; the data are plotted as mean of two independent biological replicates (n= 2) each mad e of of a technical triplicate.

Nucleotide sequences:

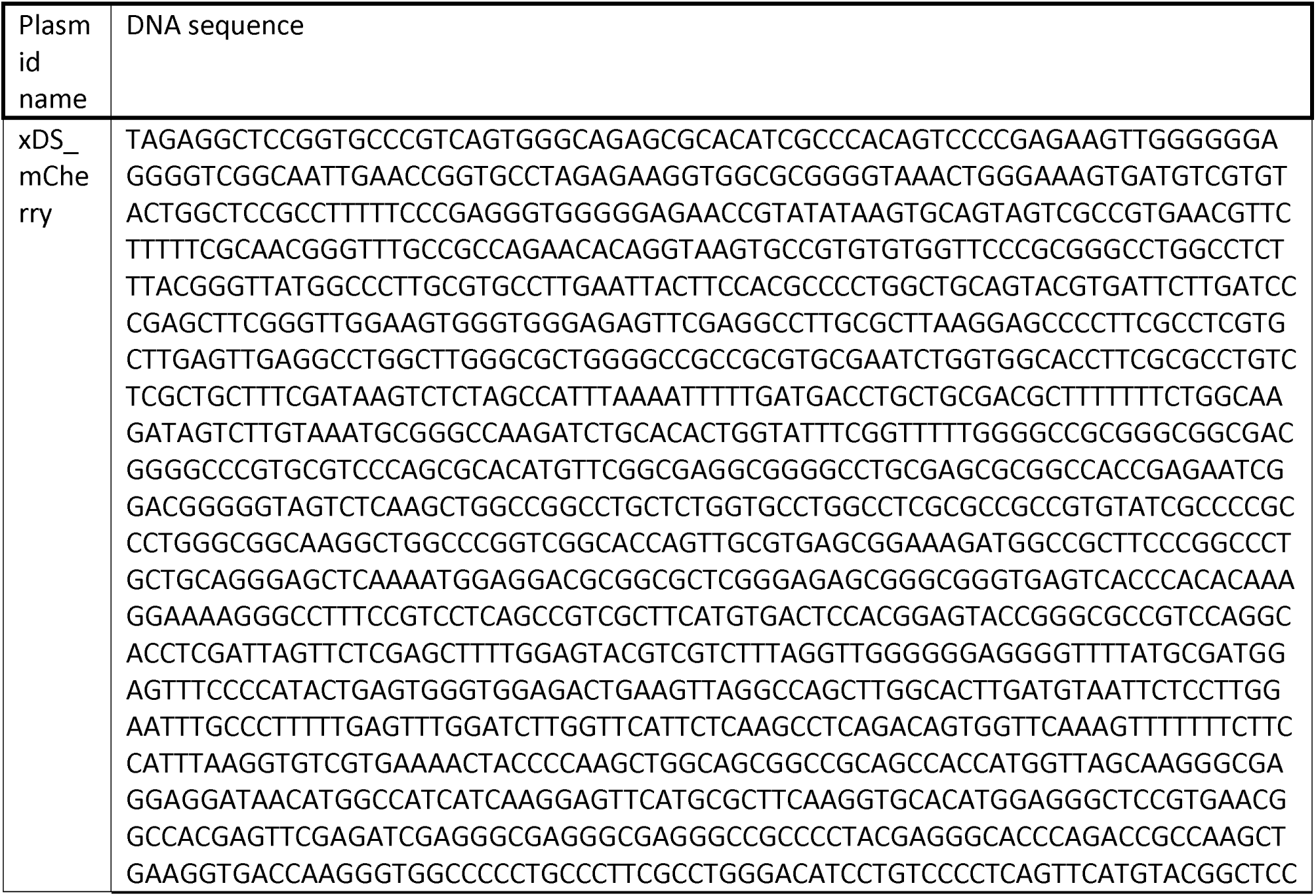
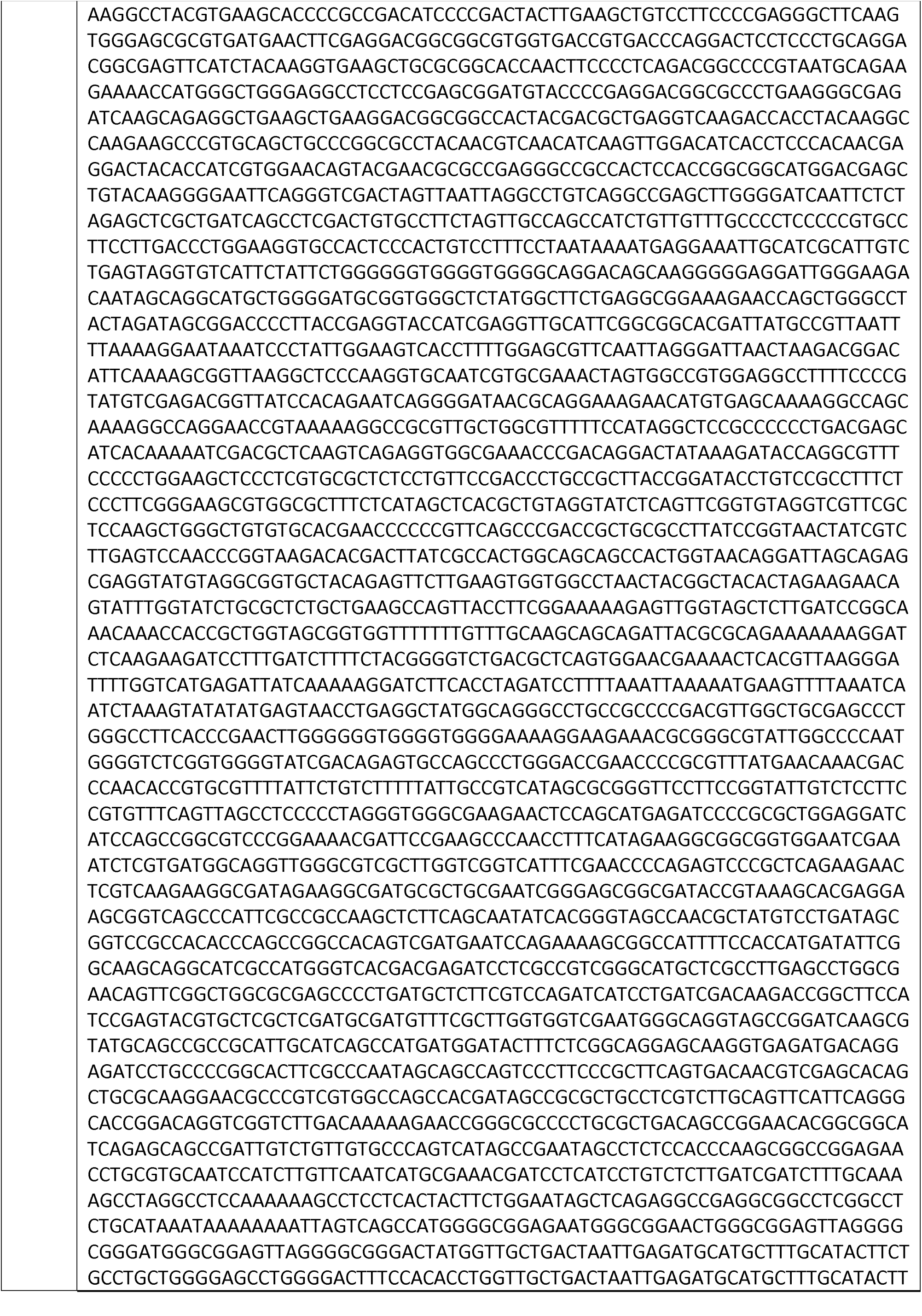
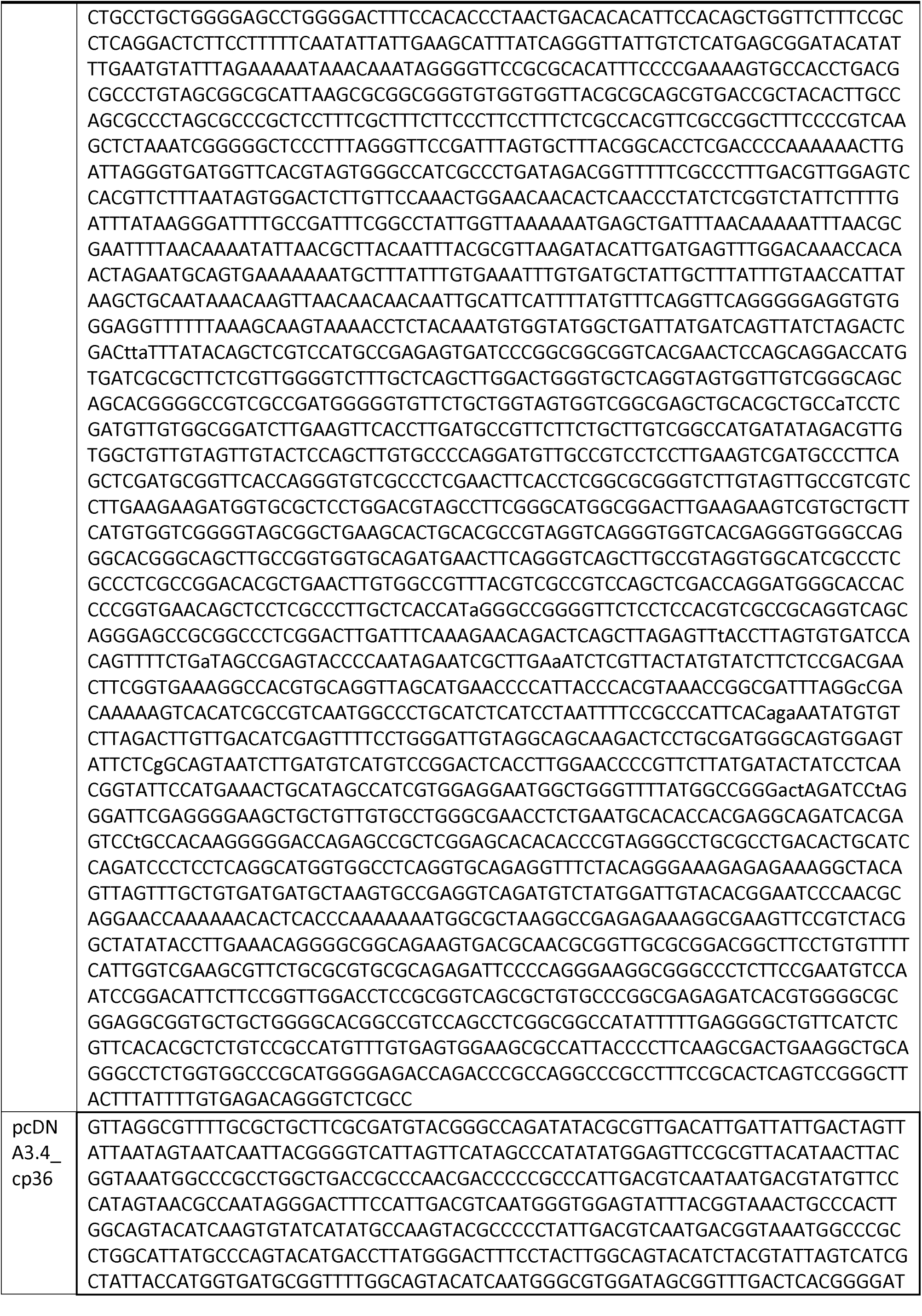
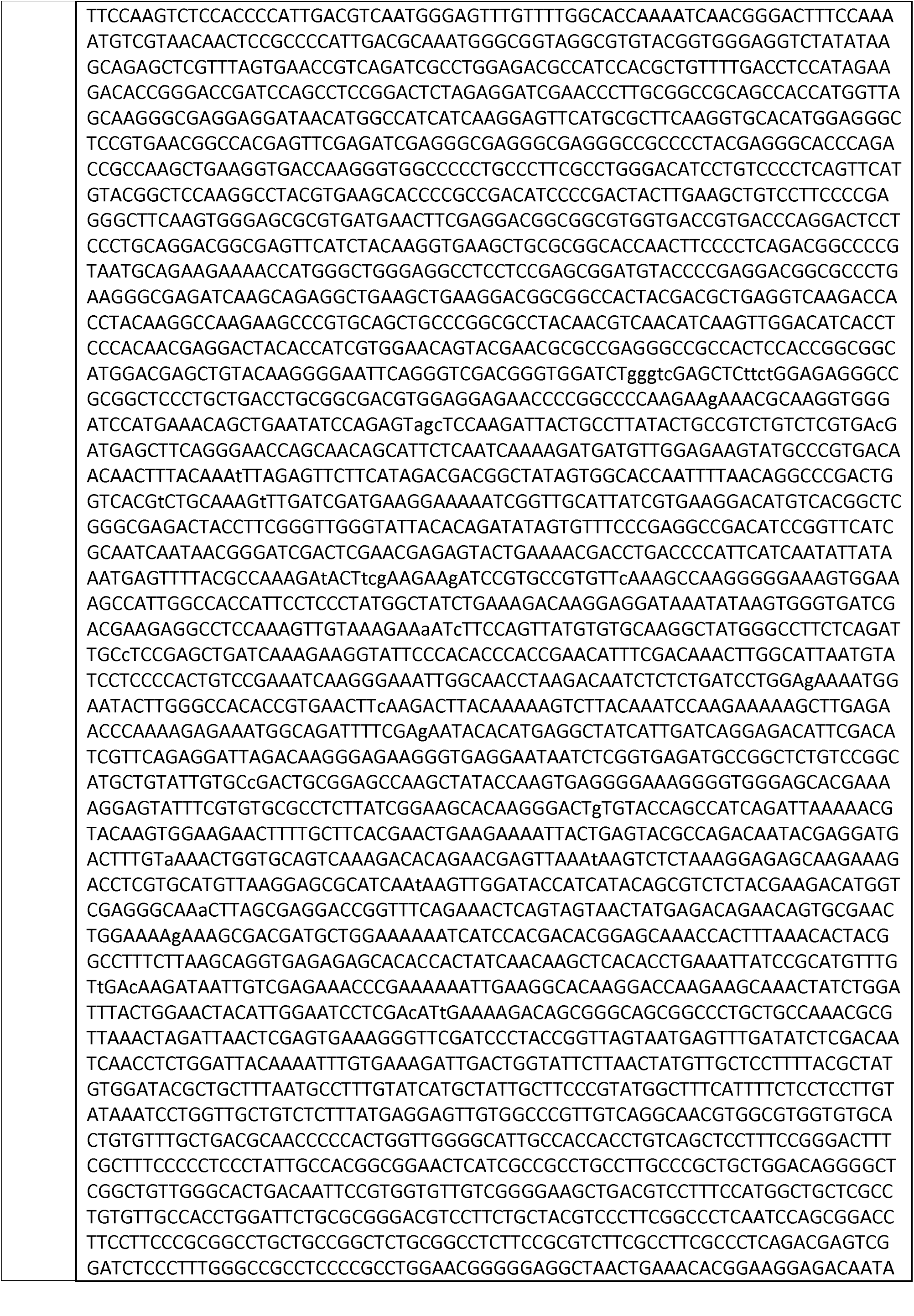
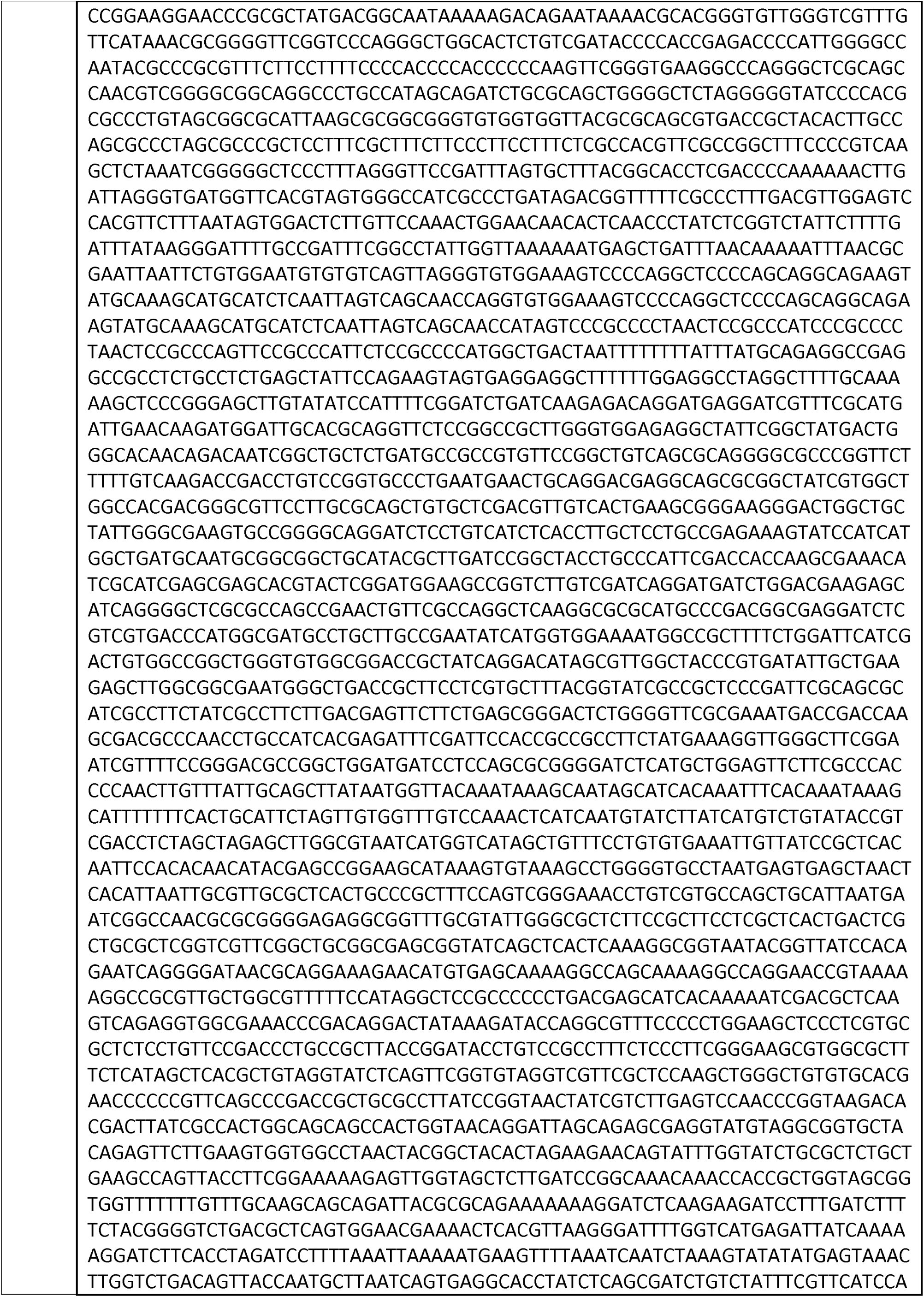
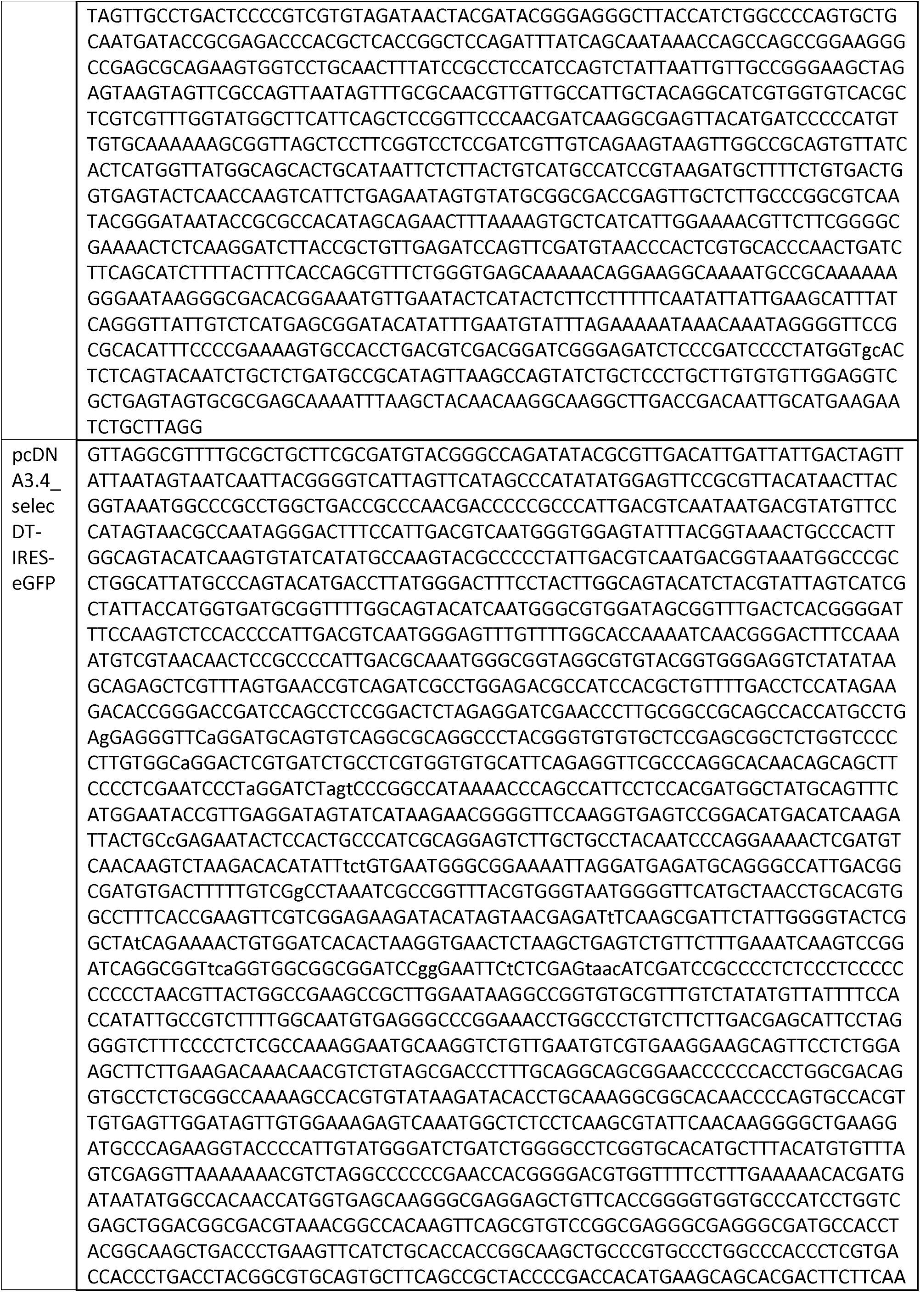
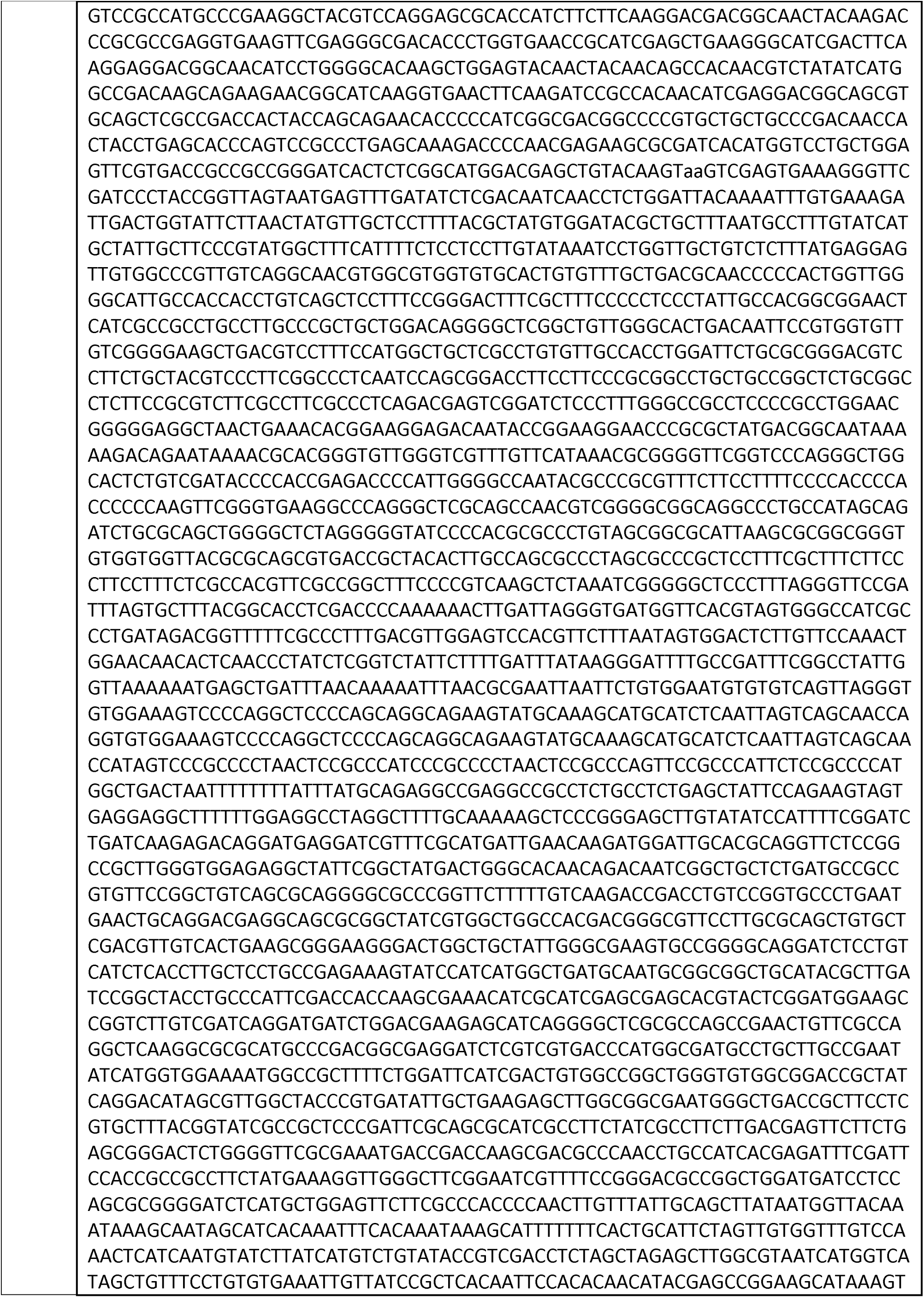
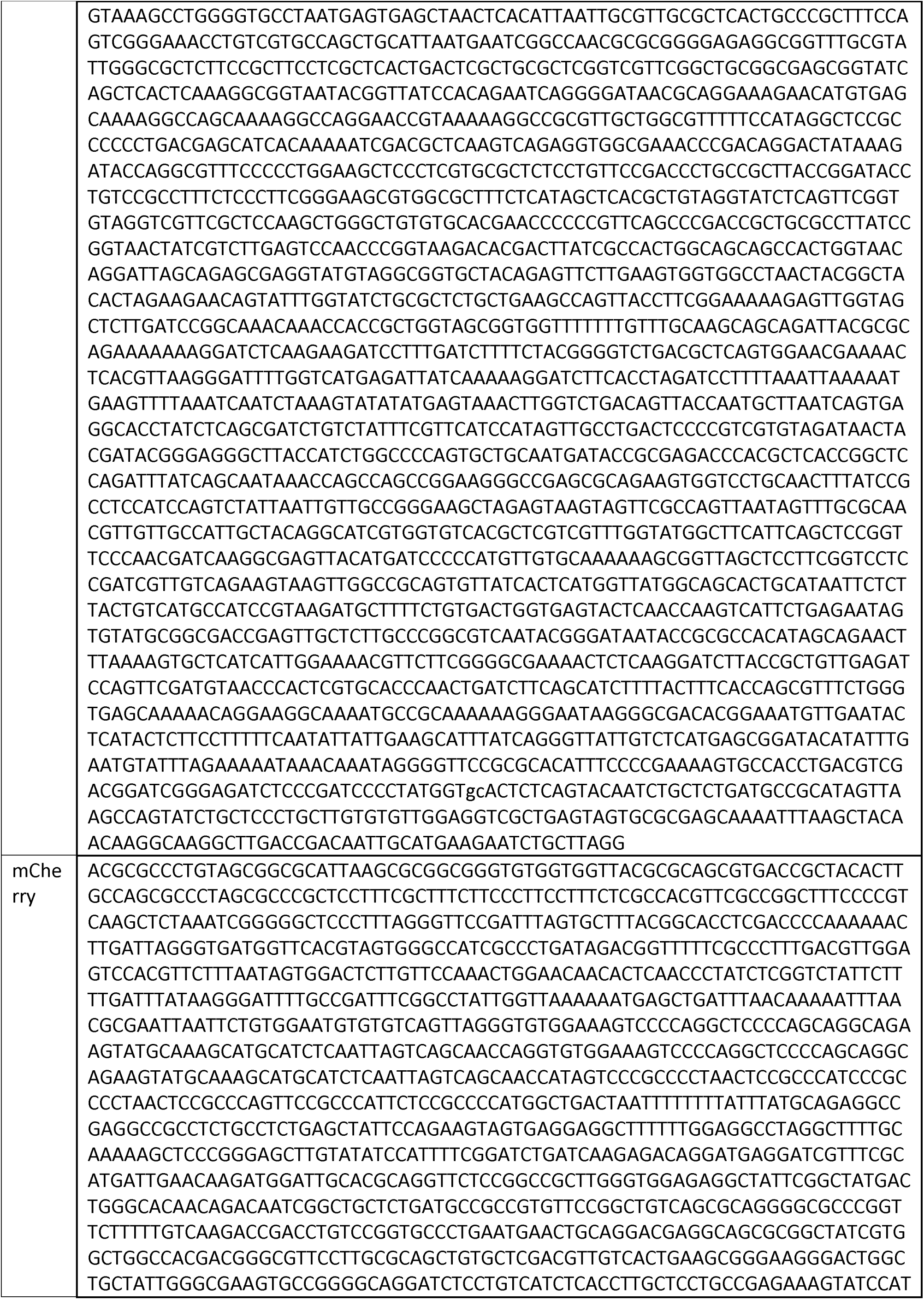
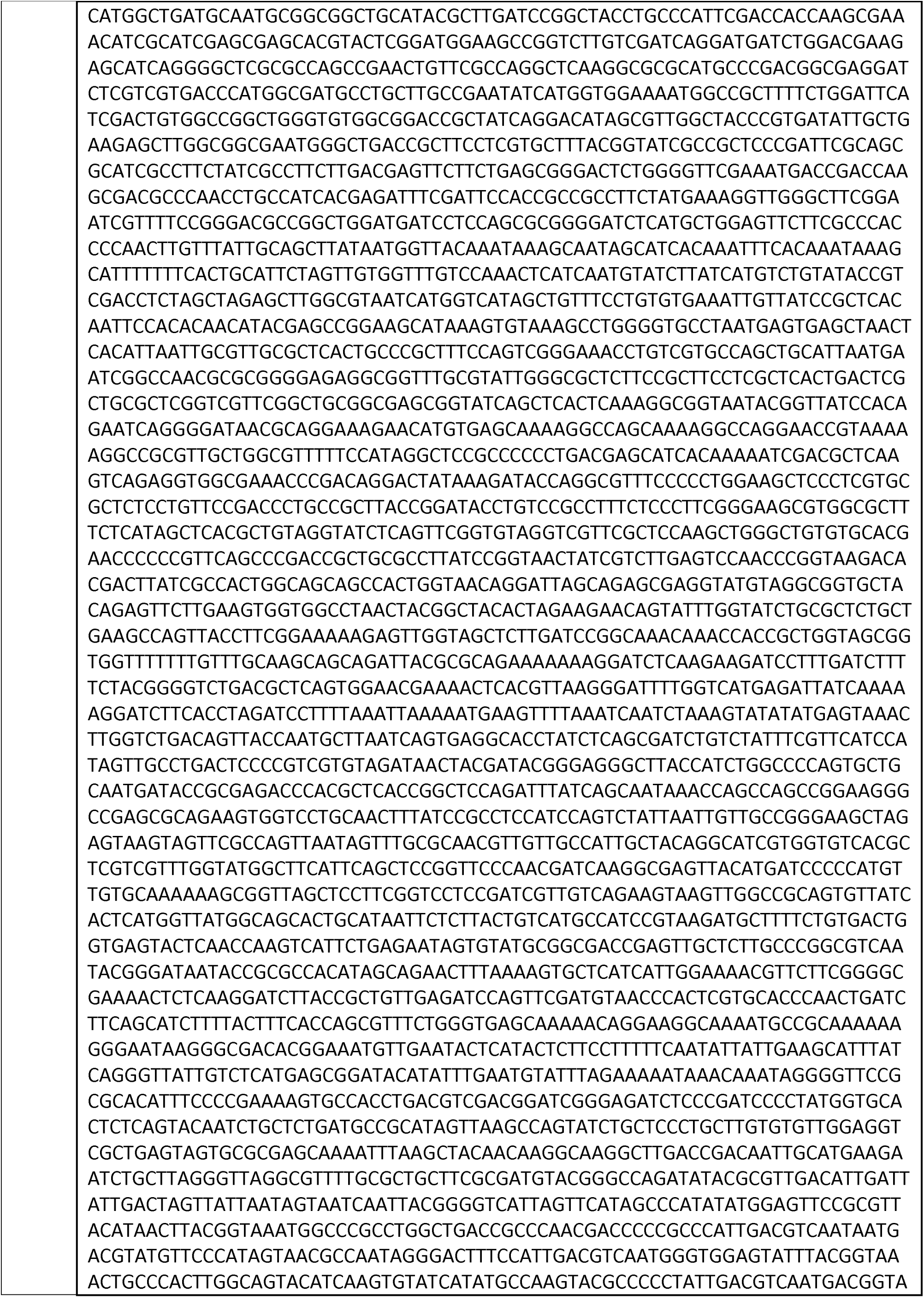
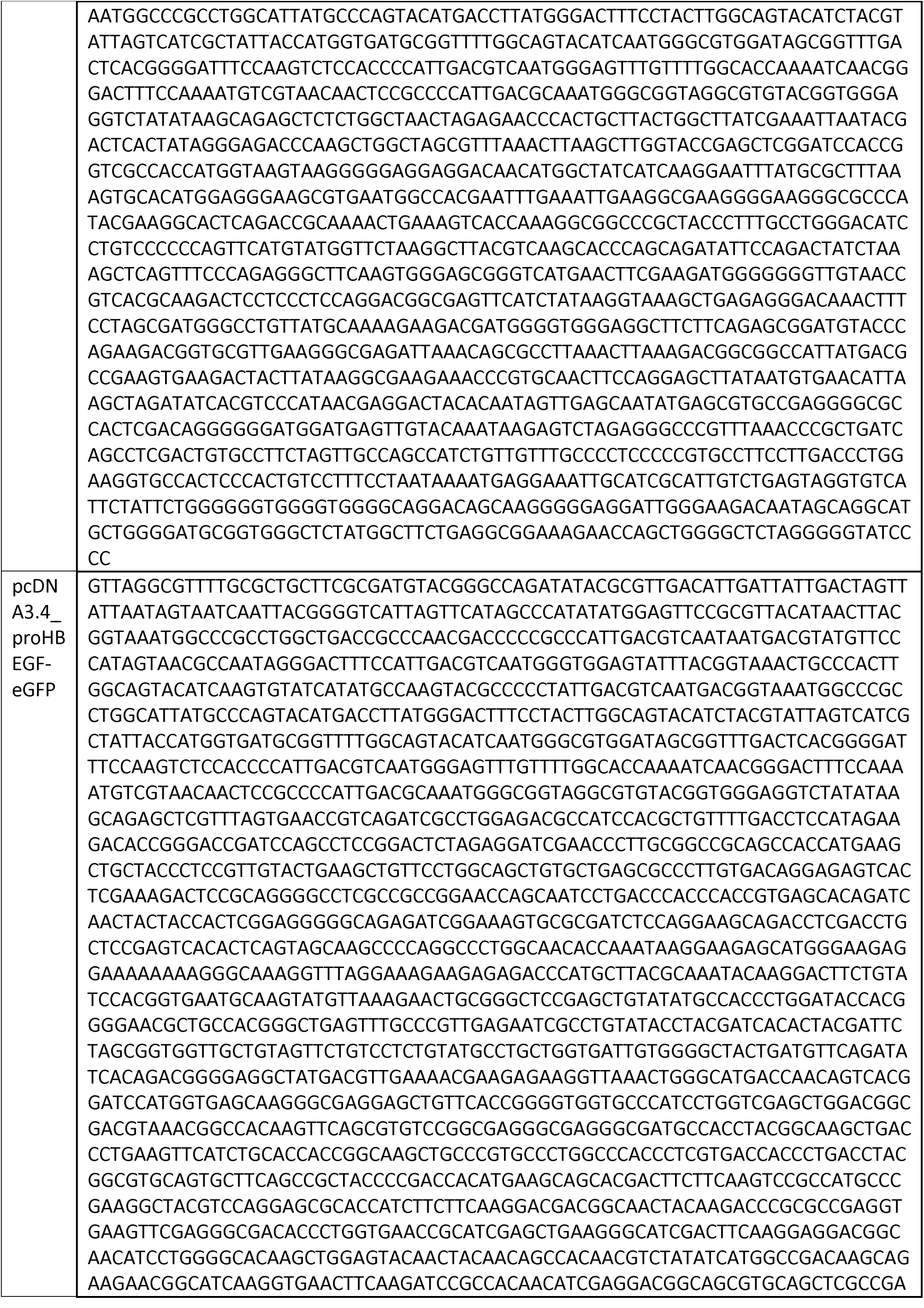
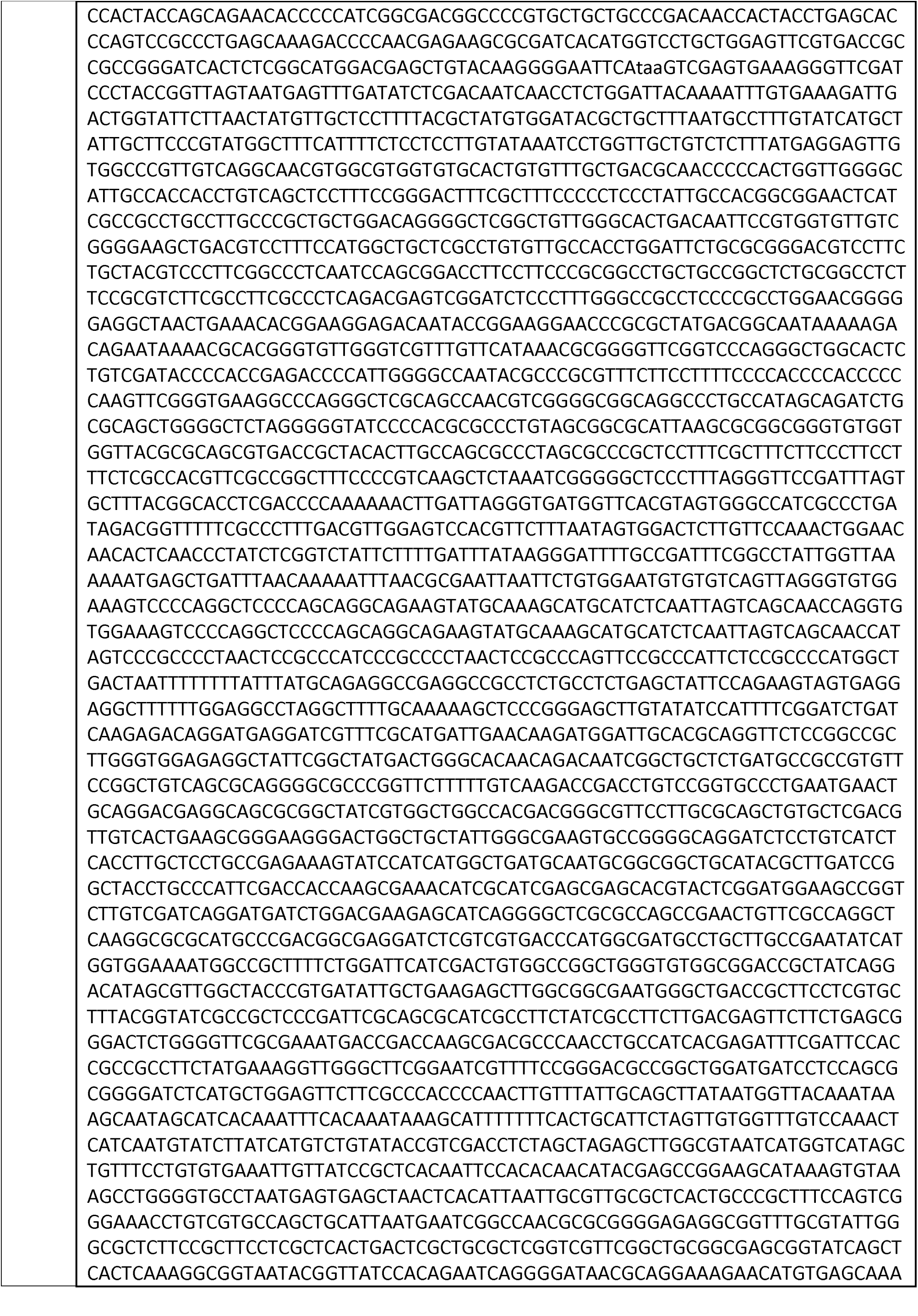
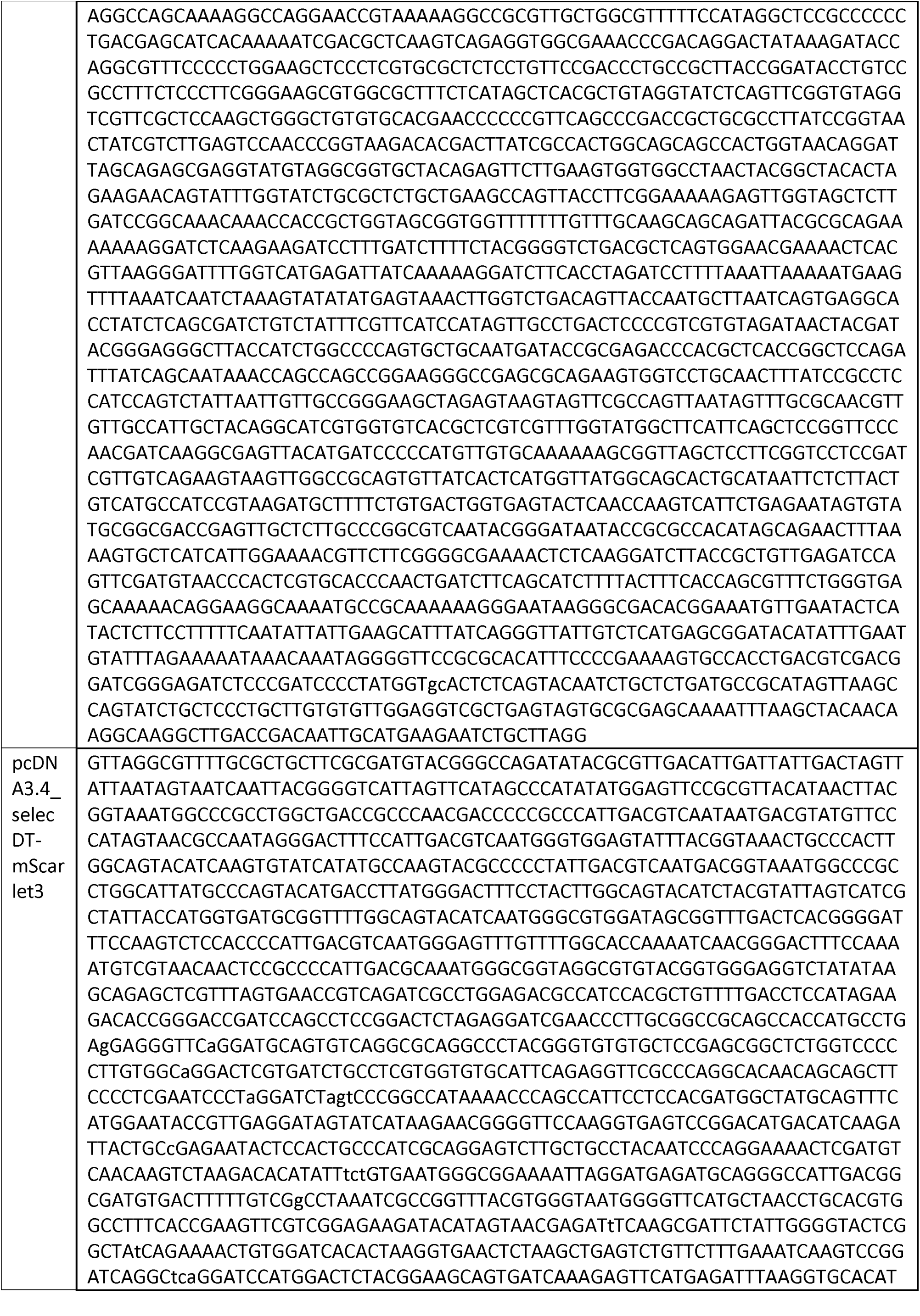
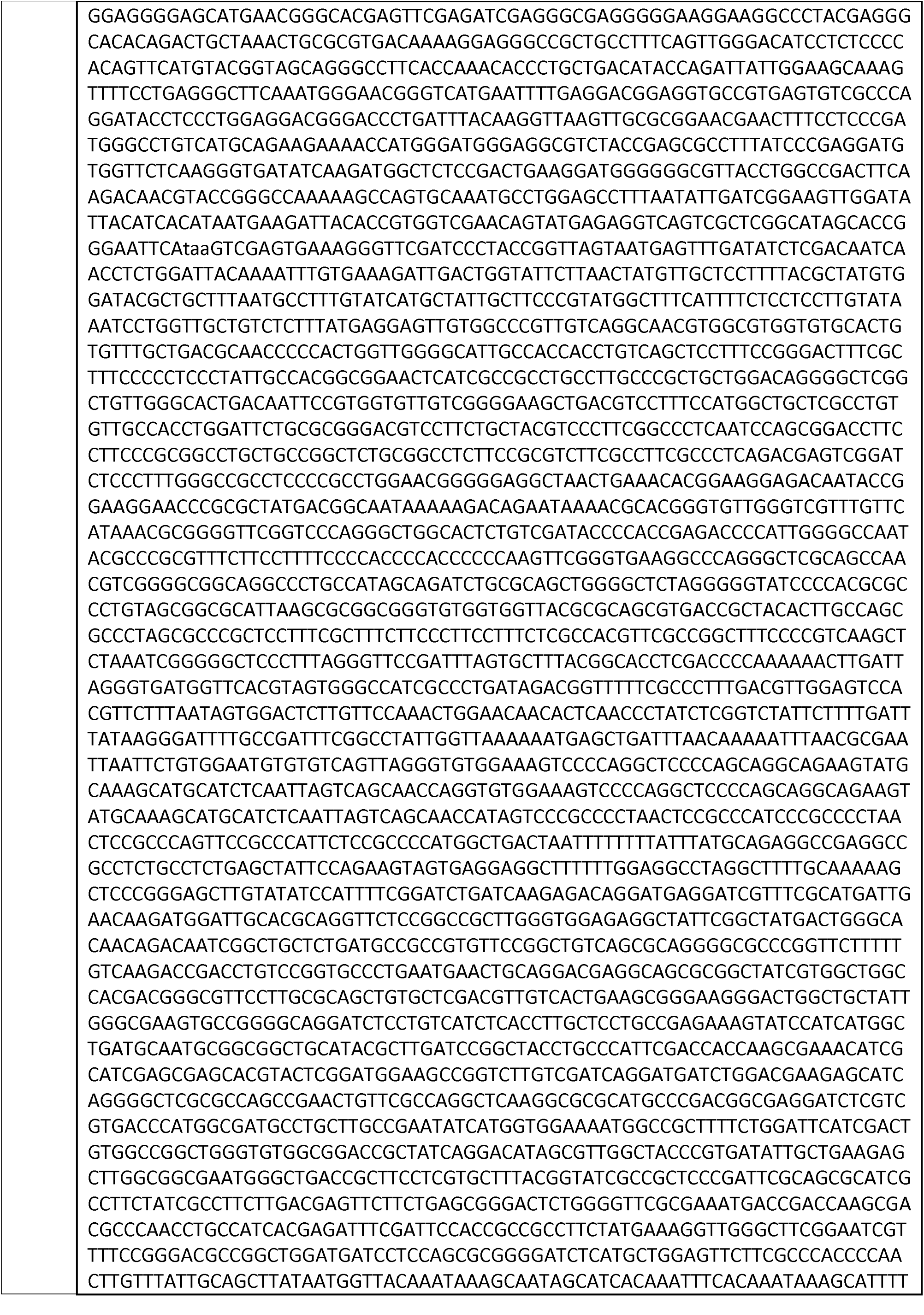
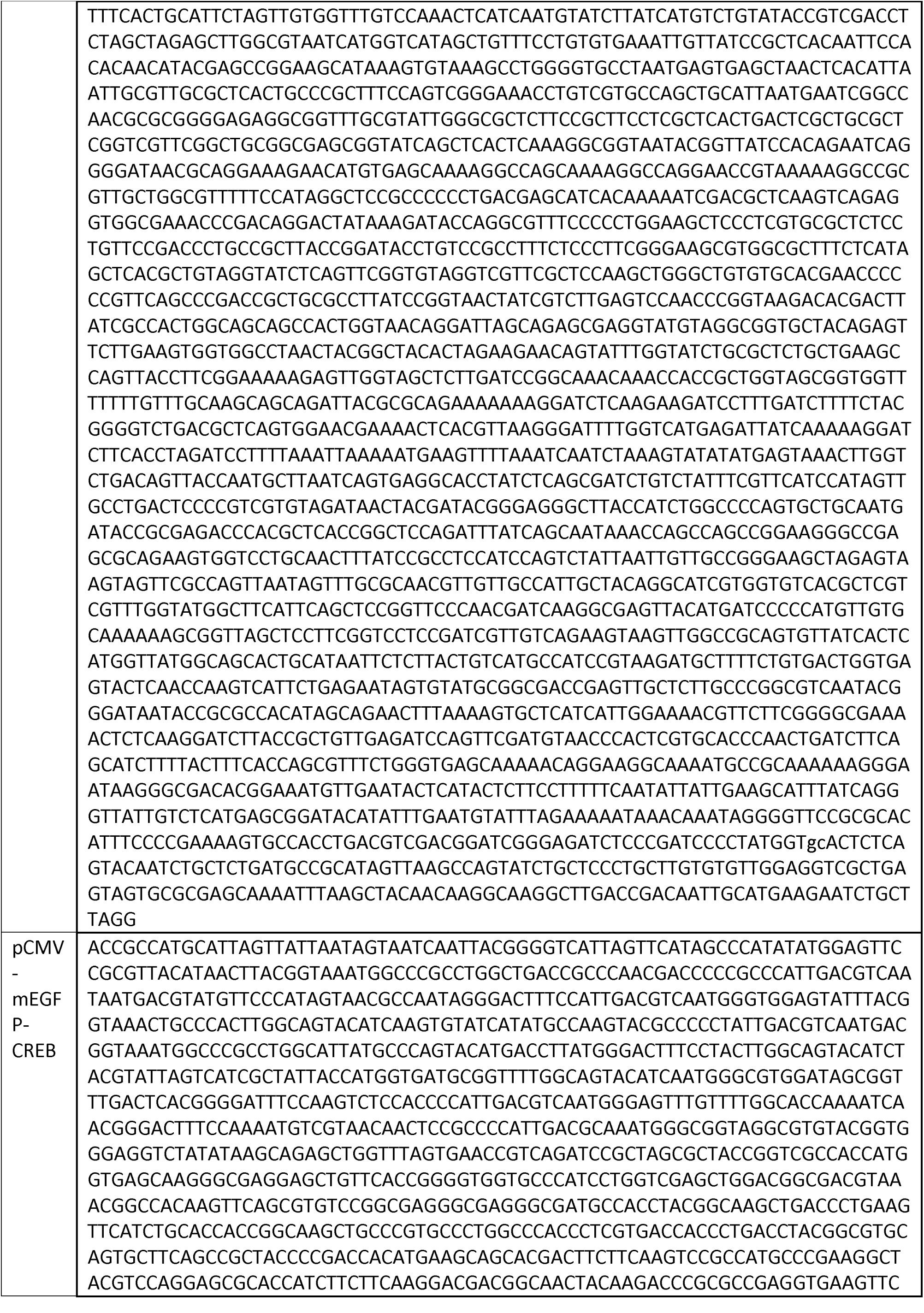
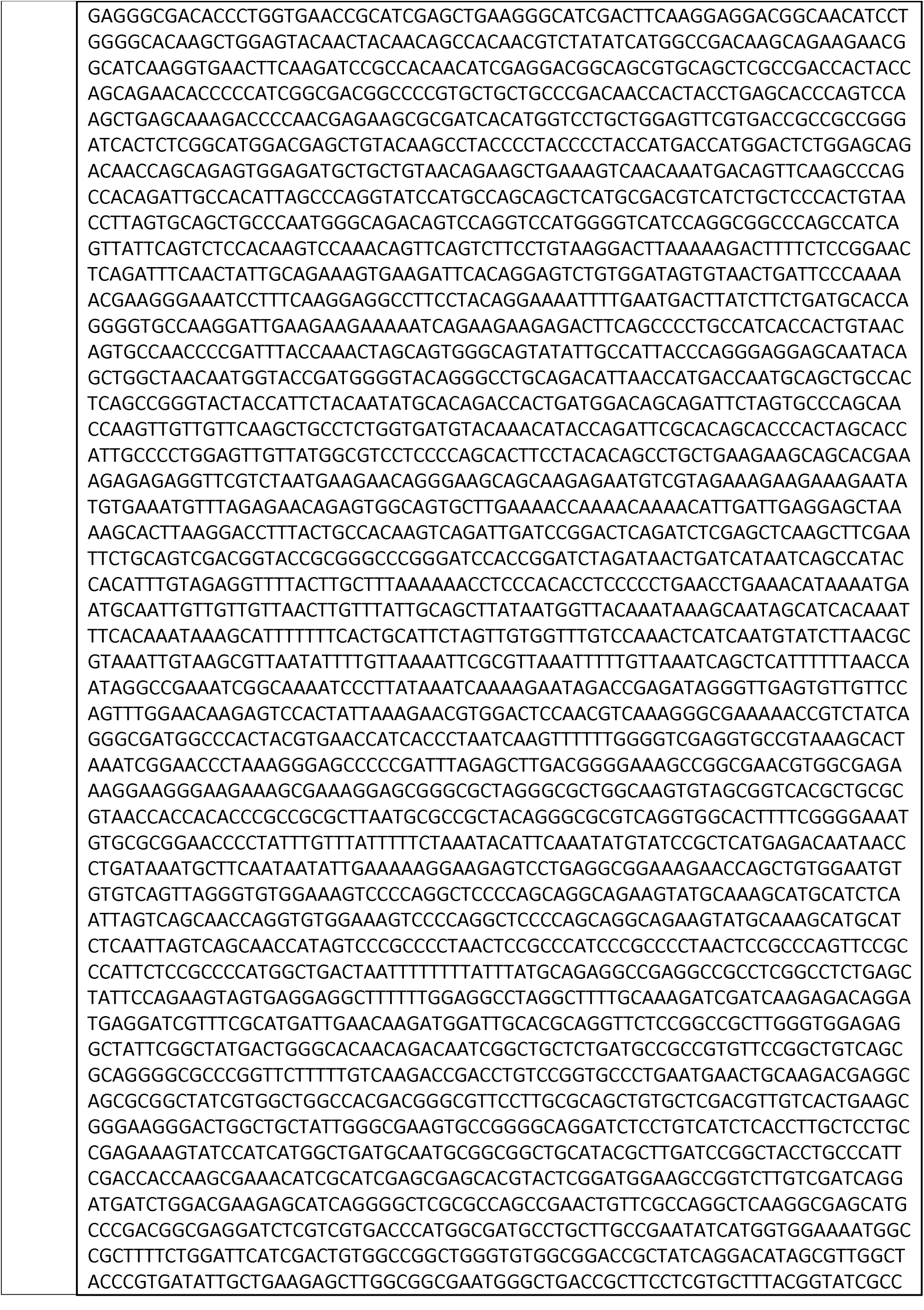
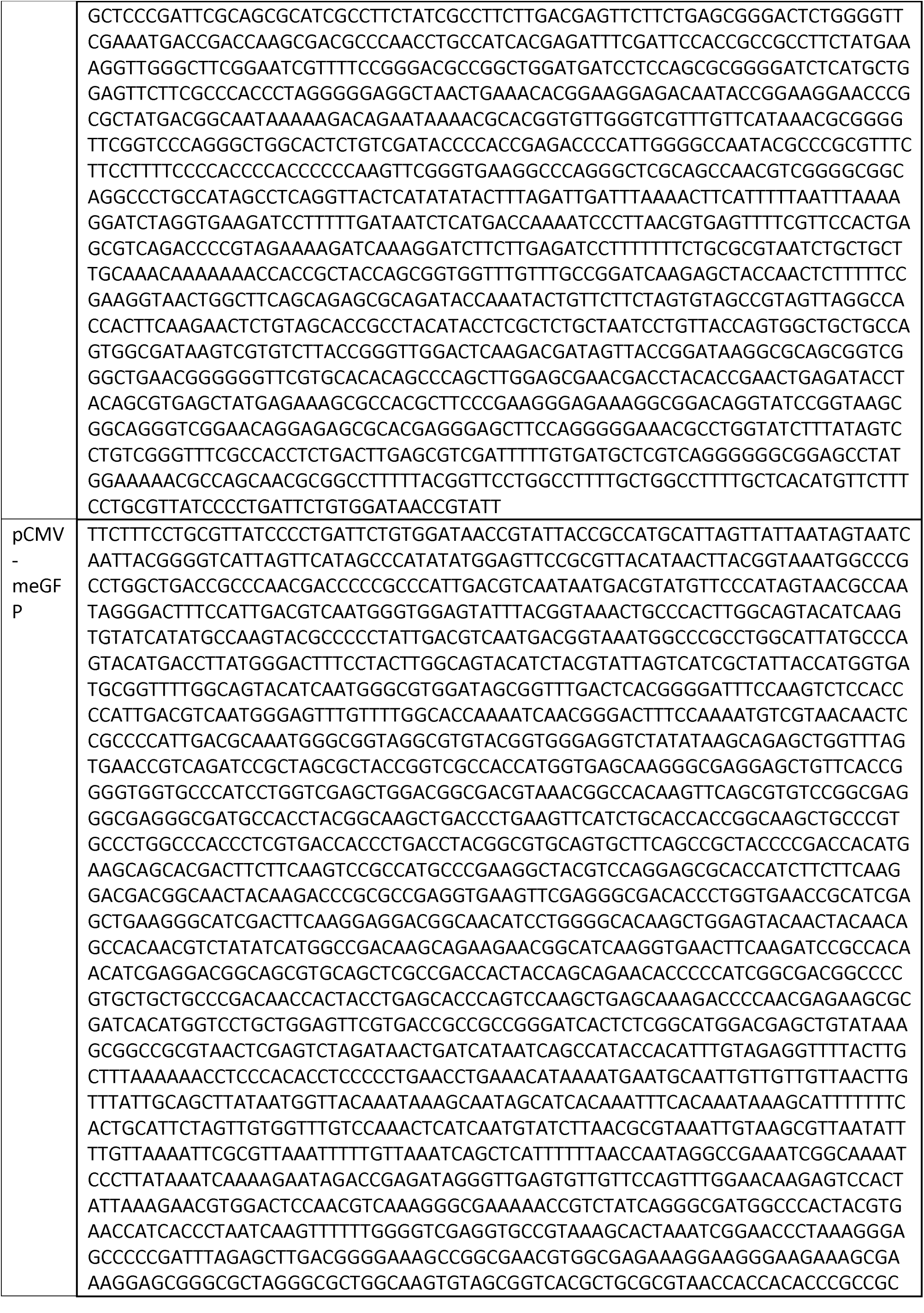
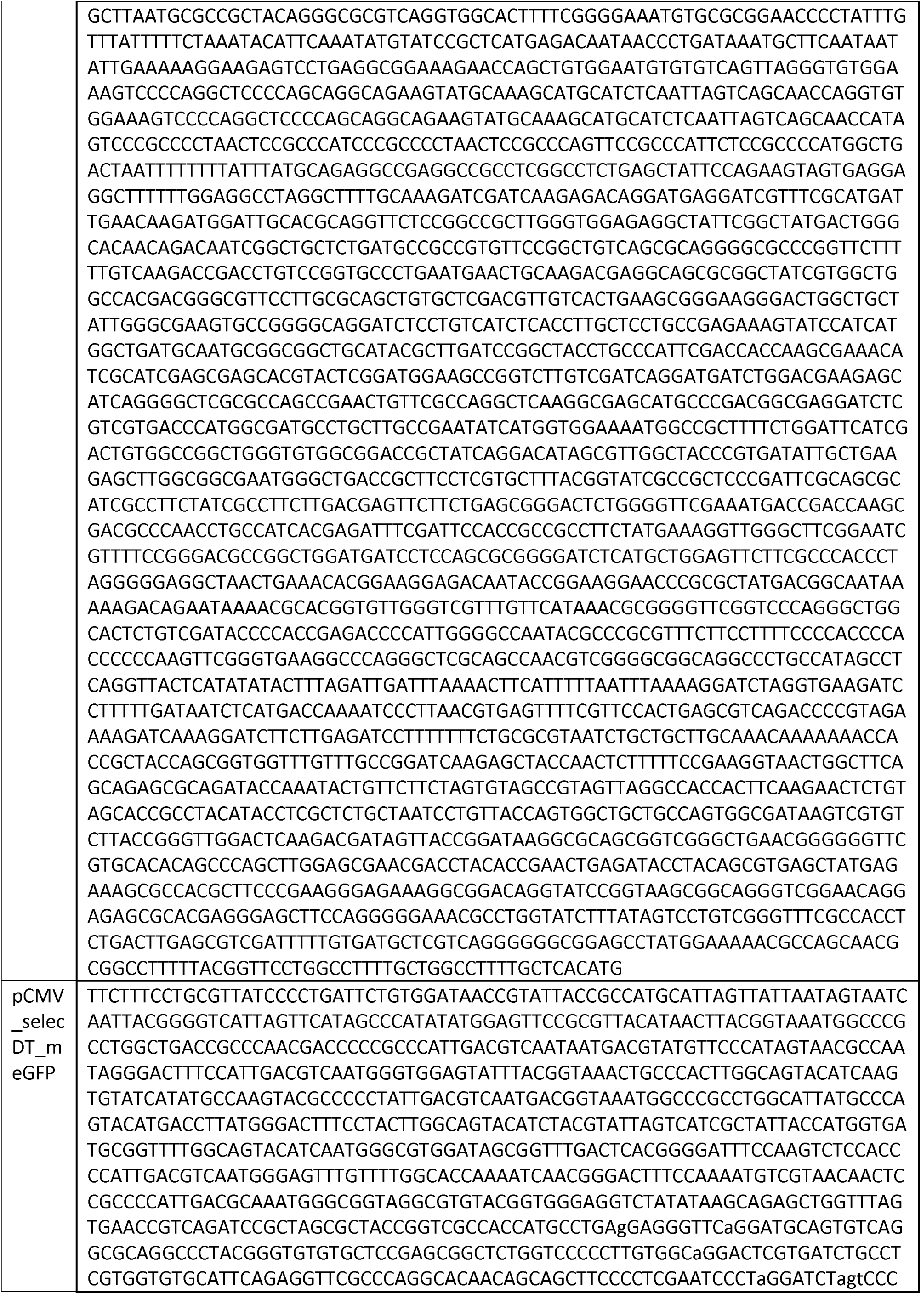
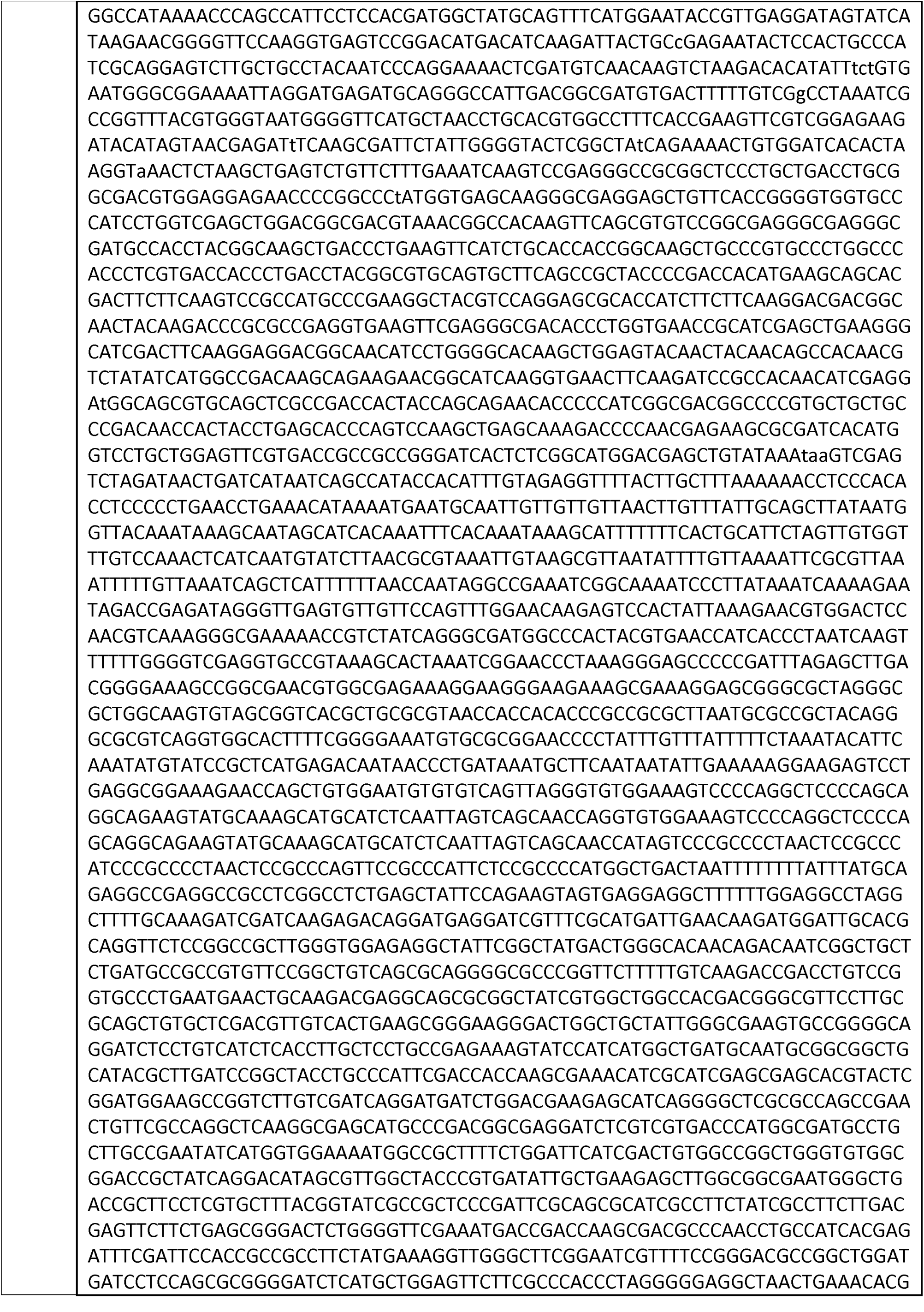
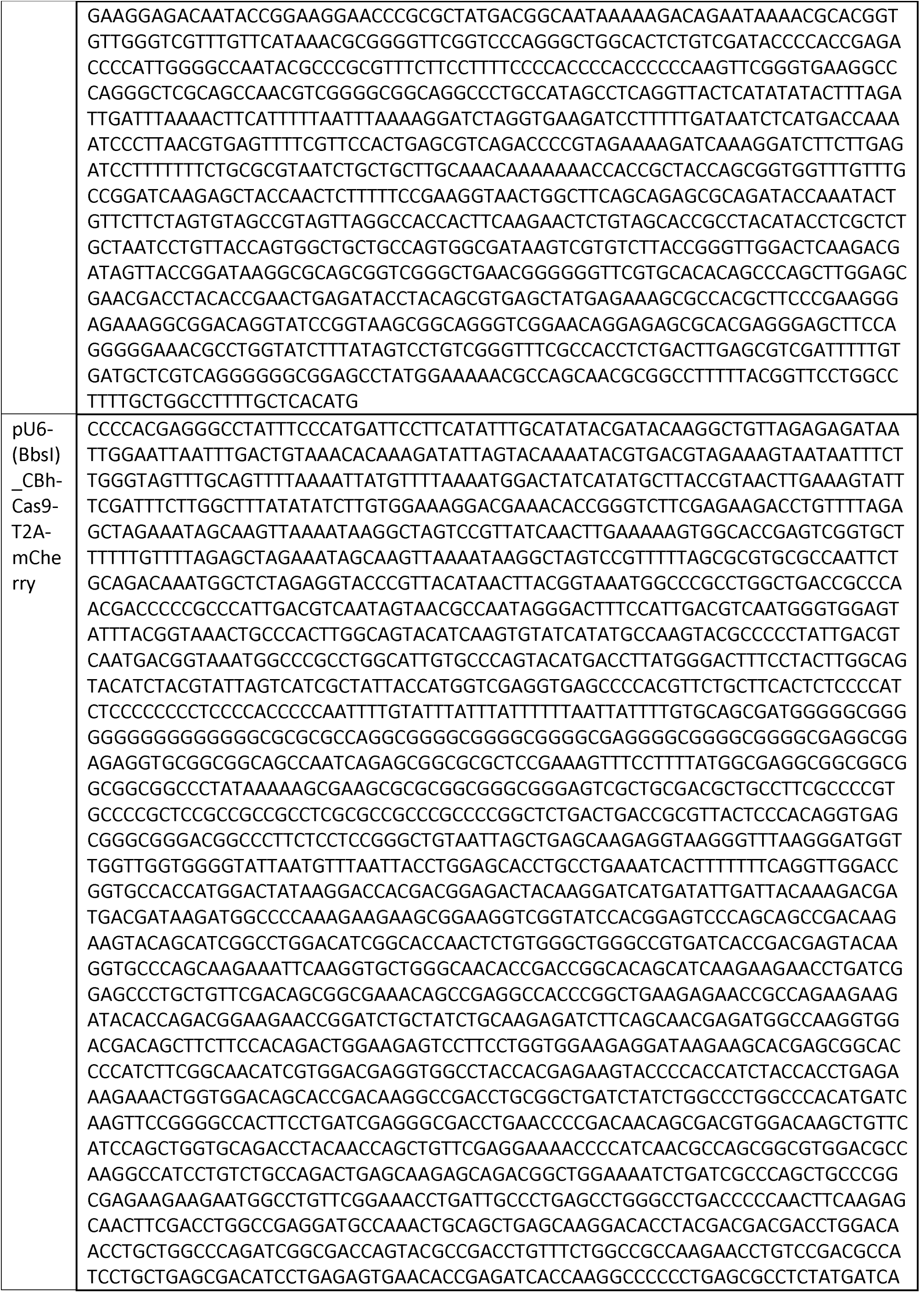
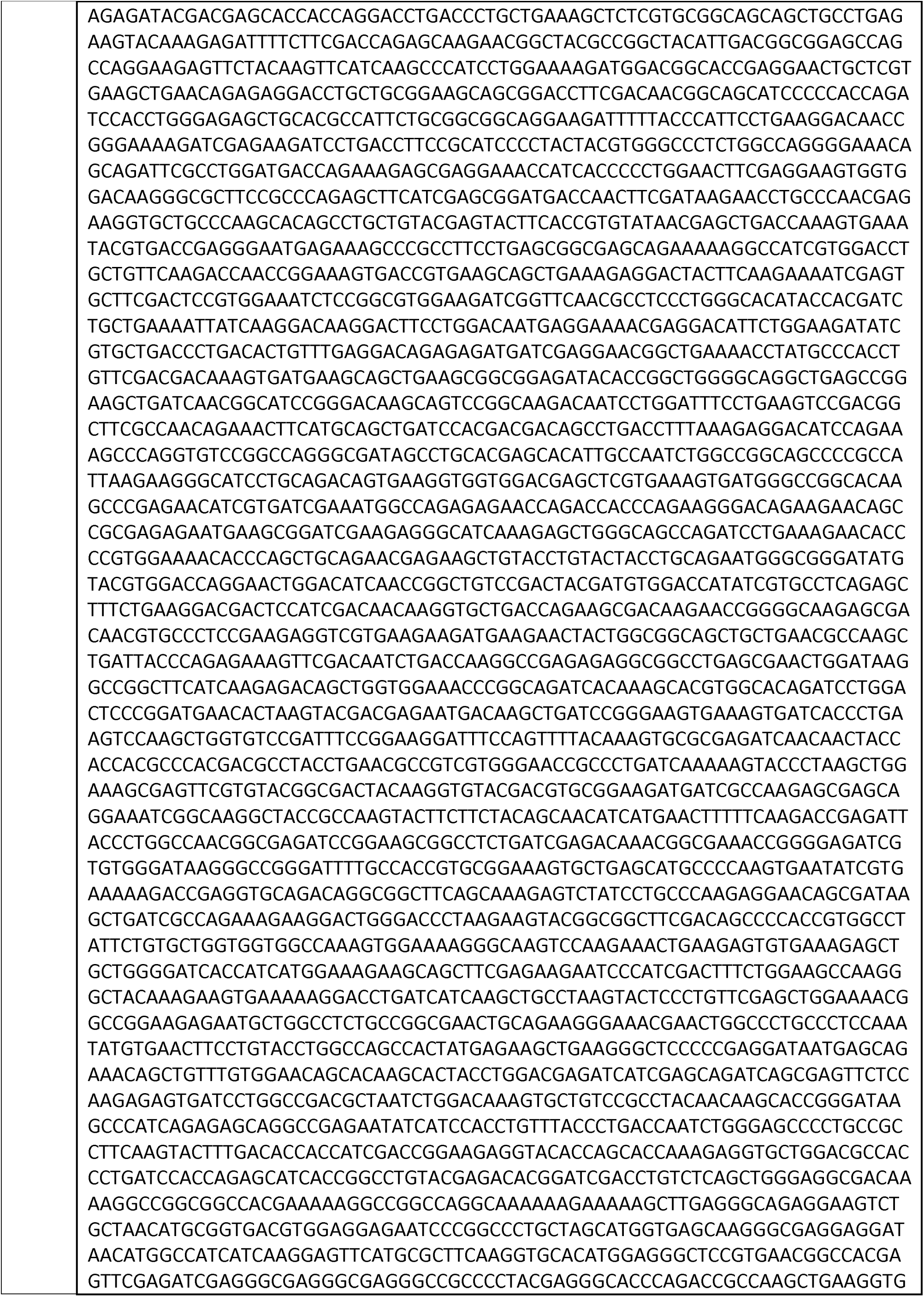
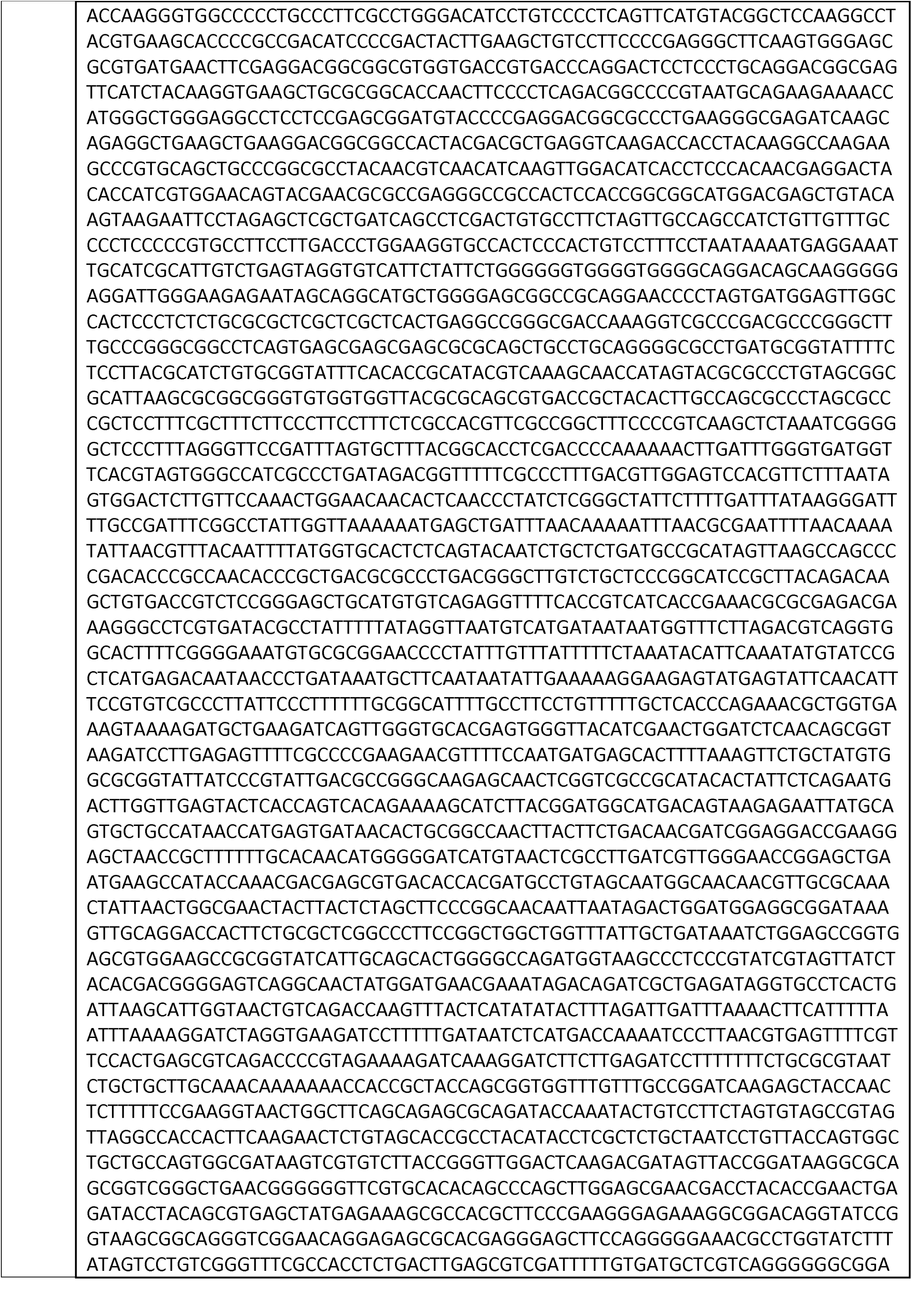
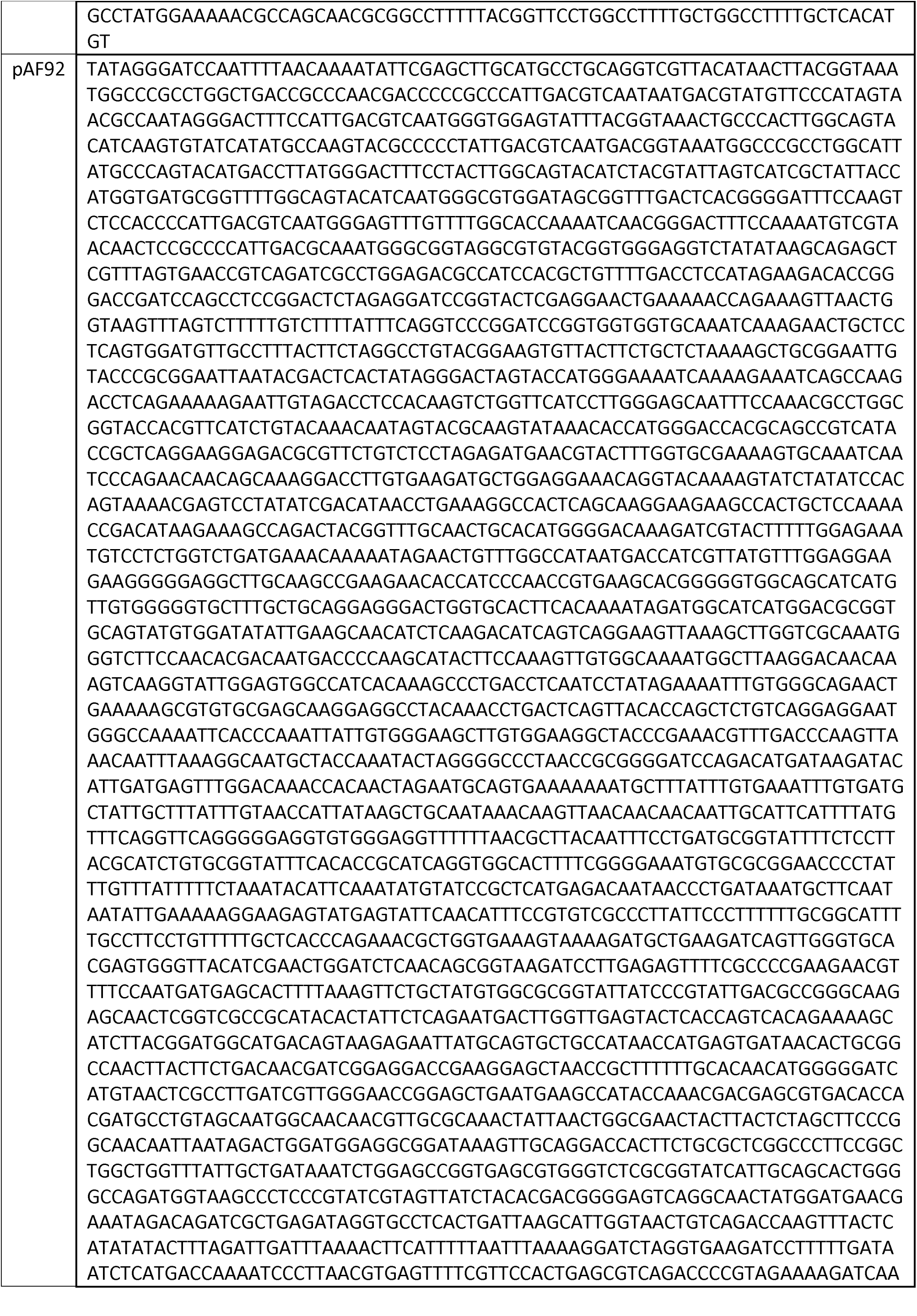
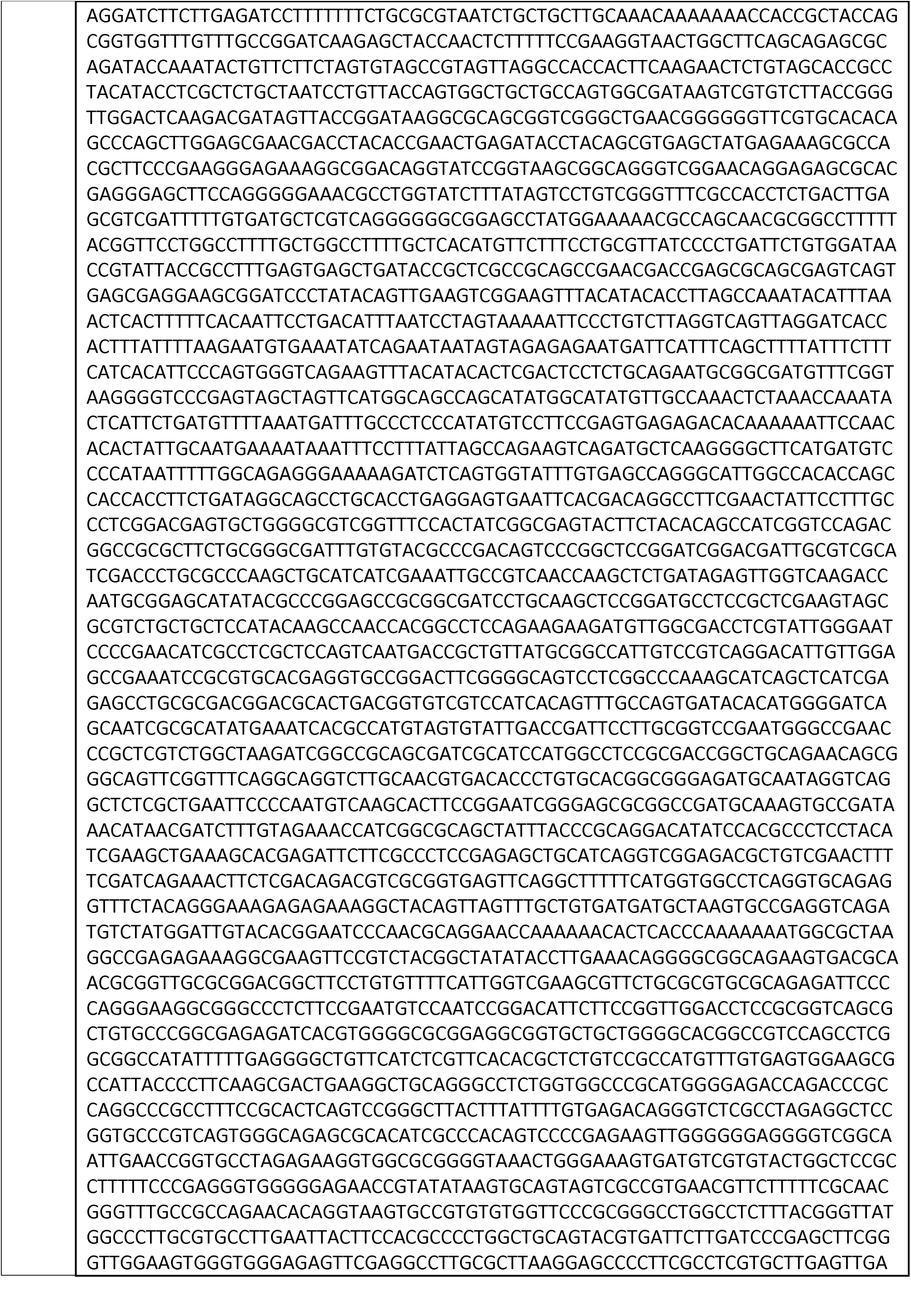
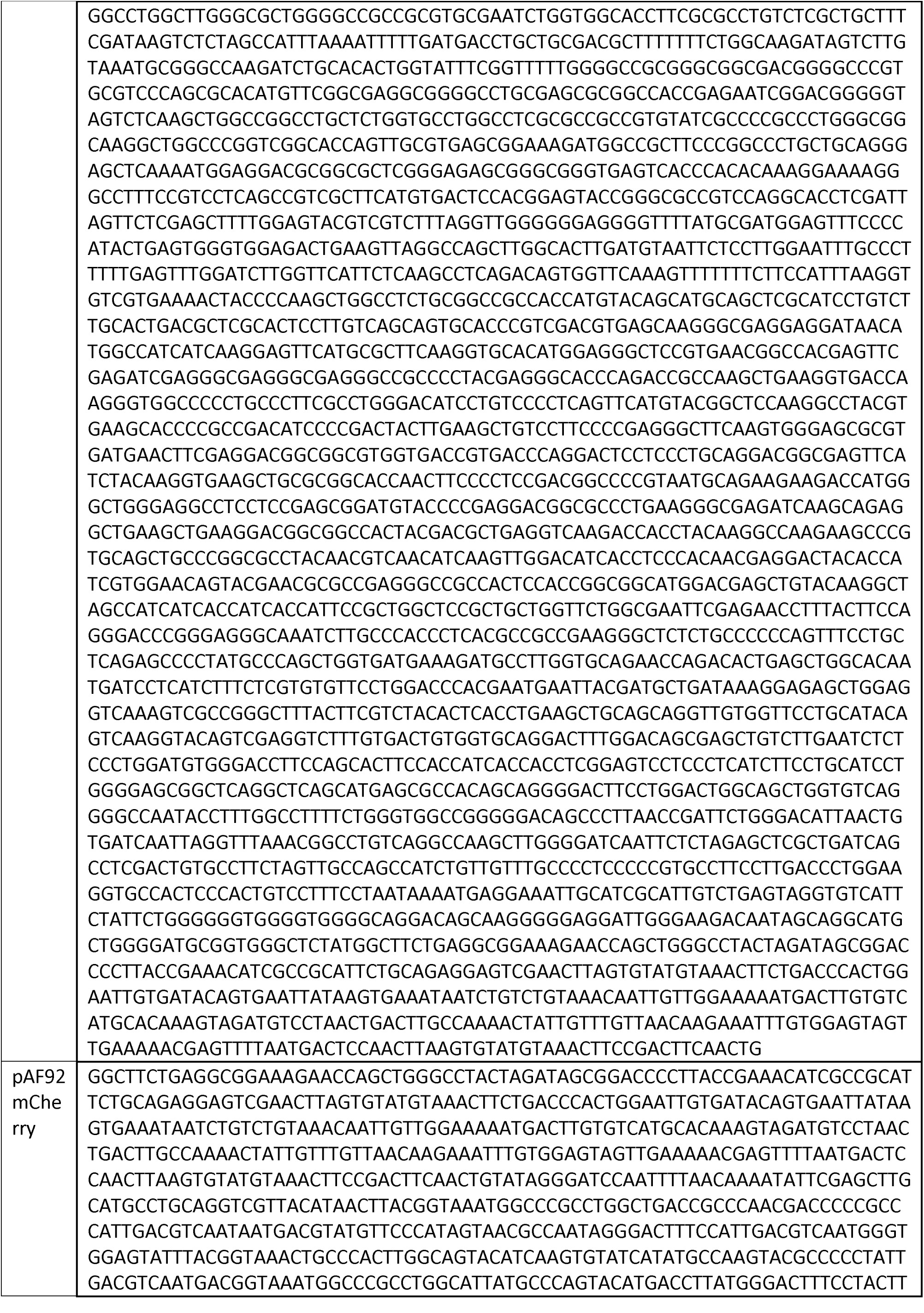
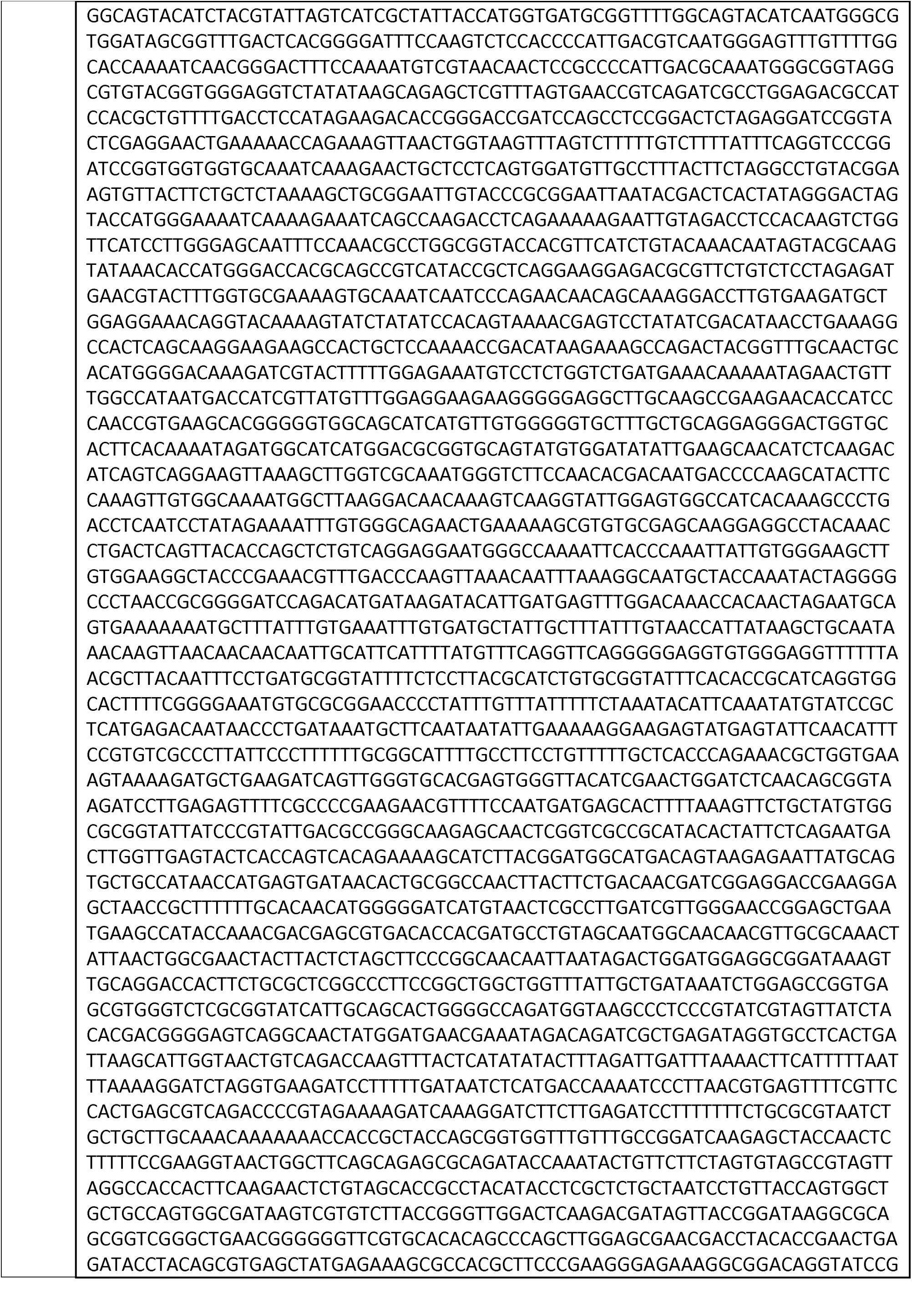
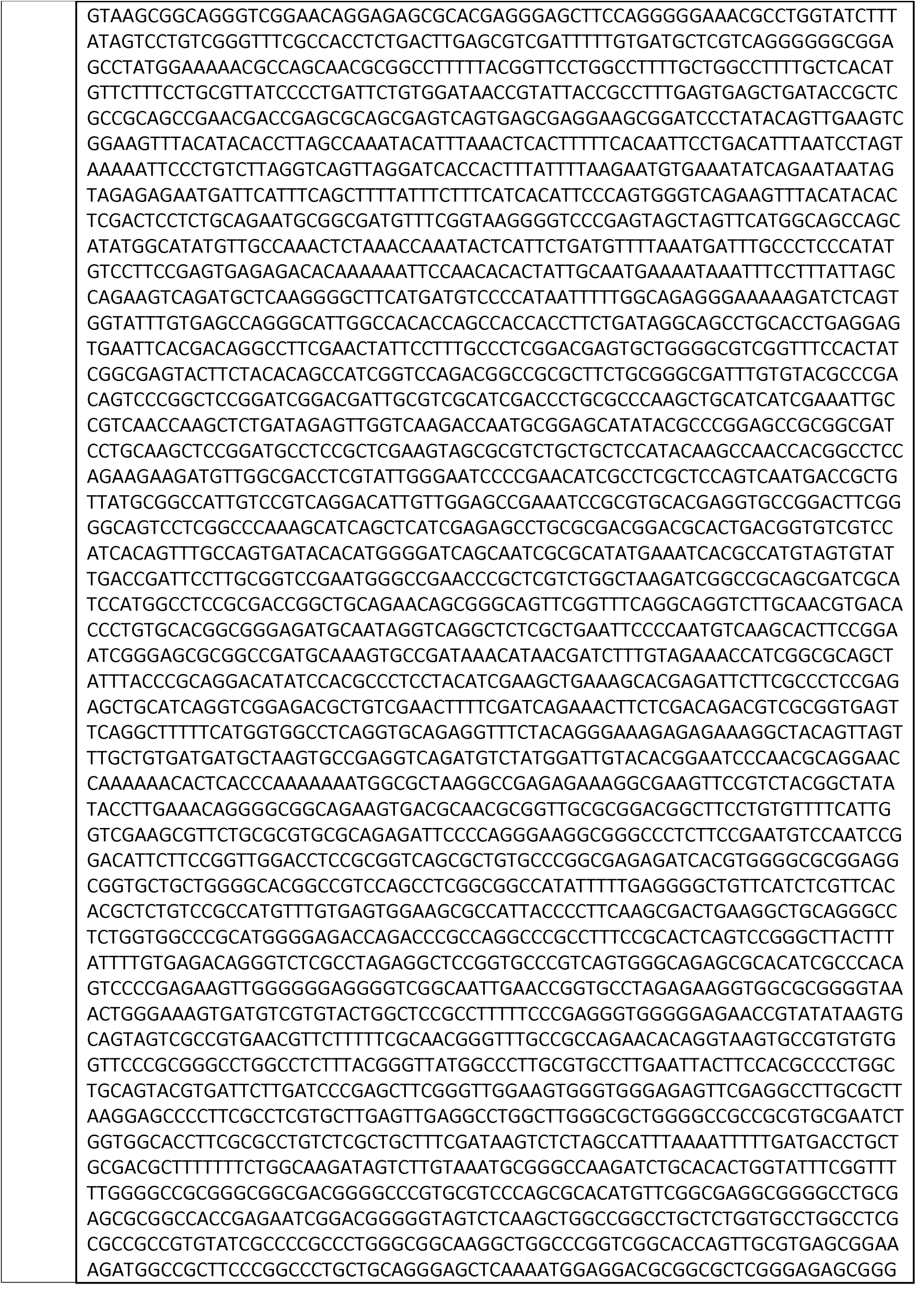
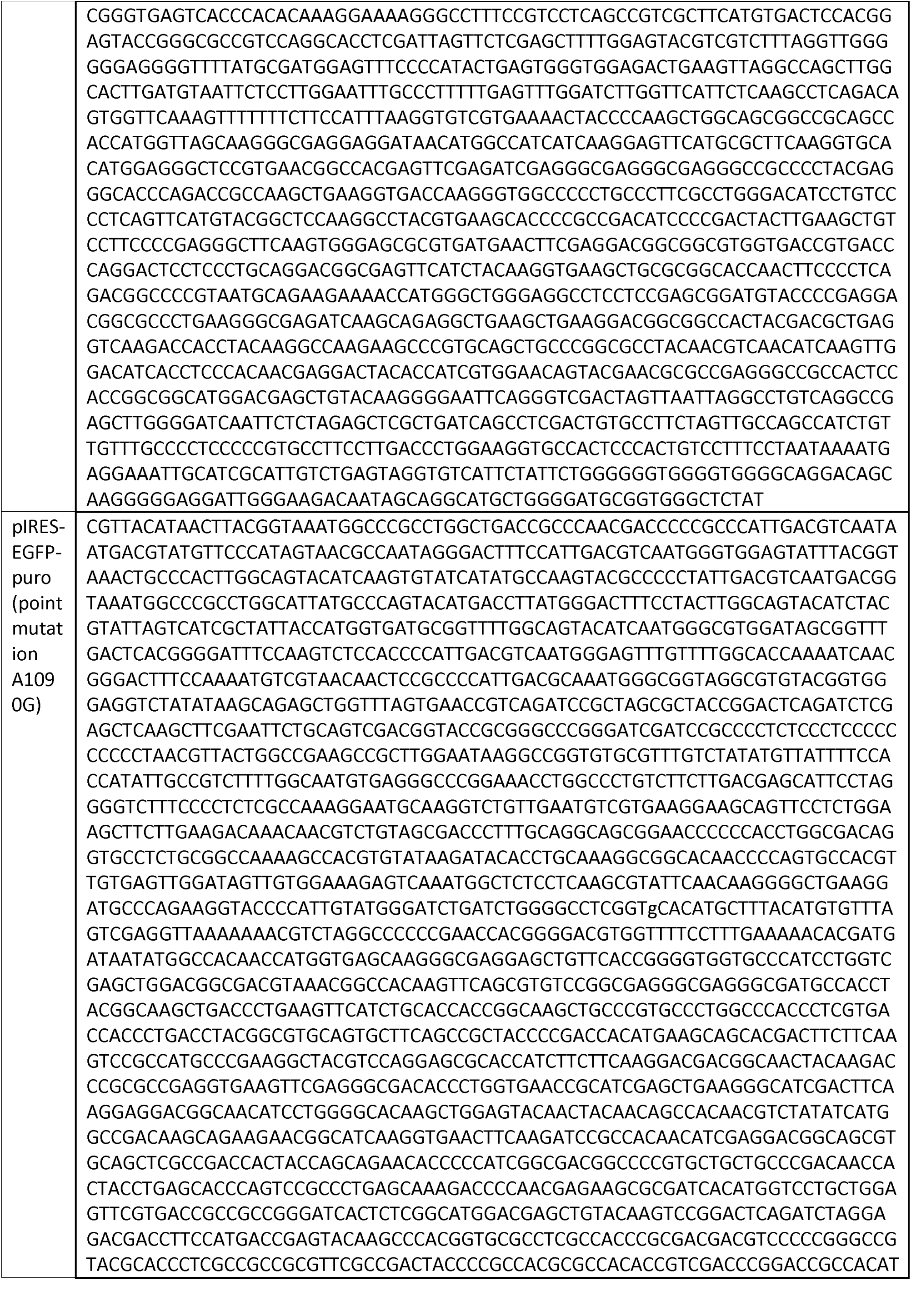
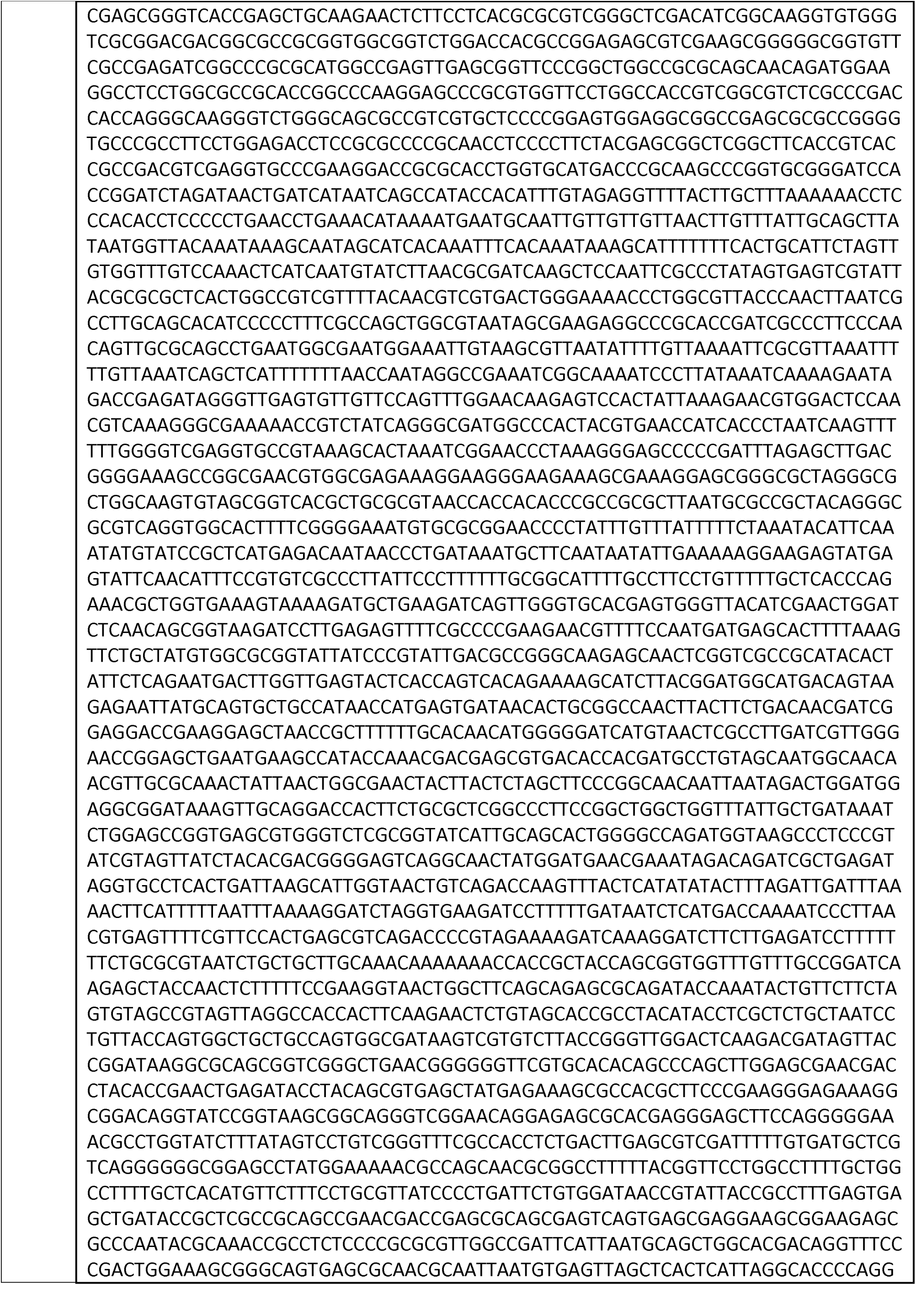
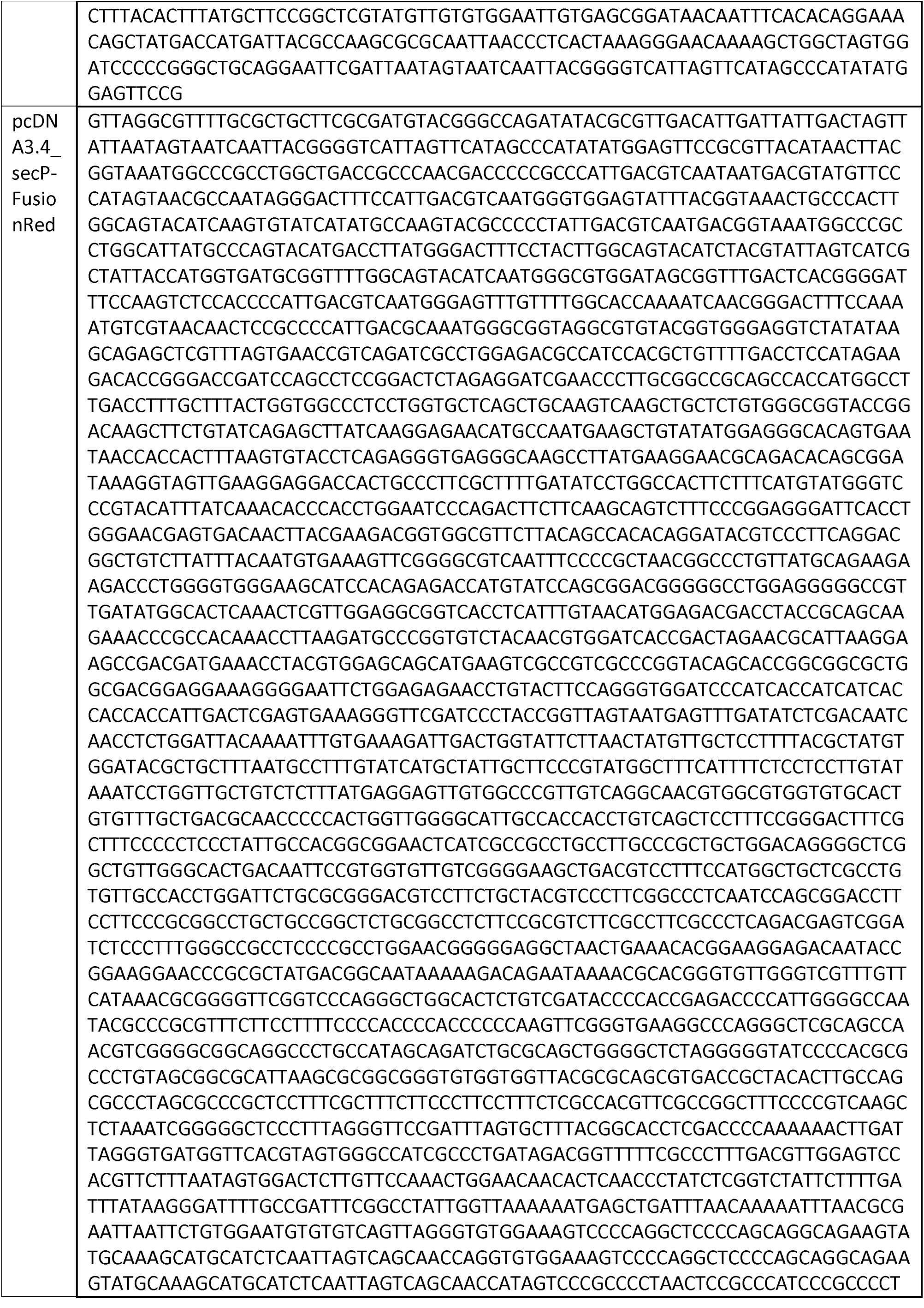
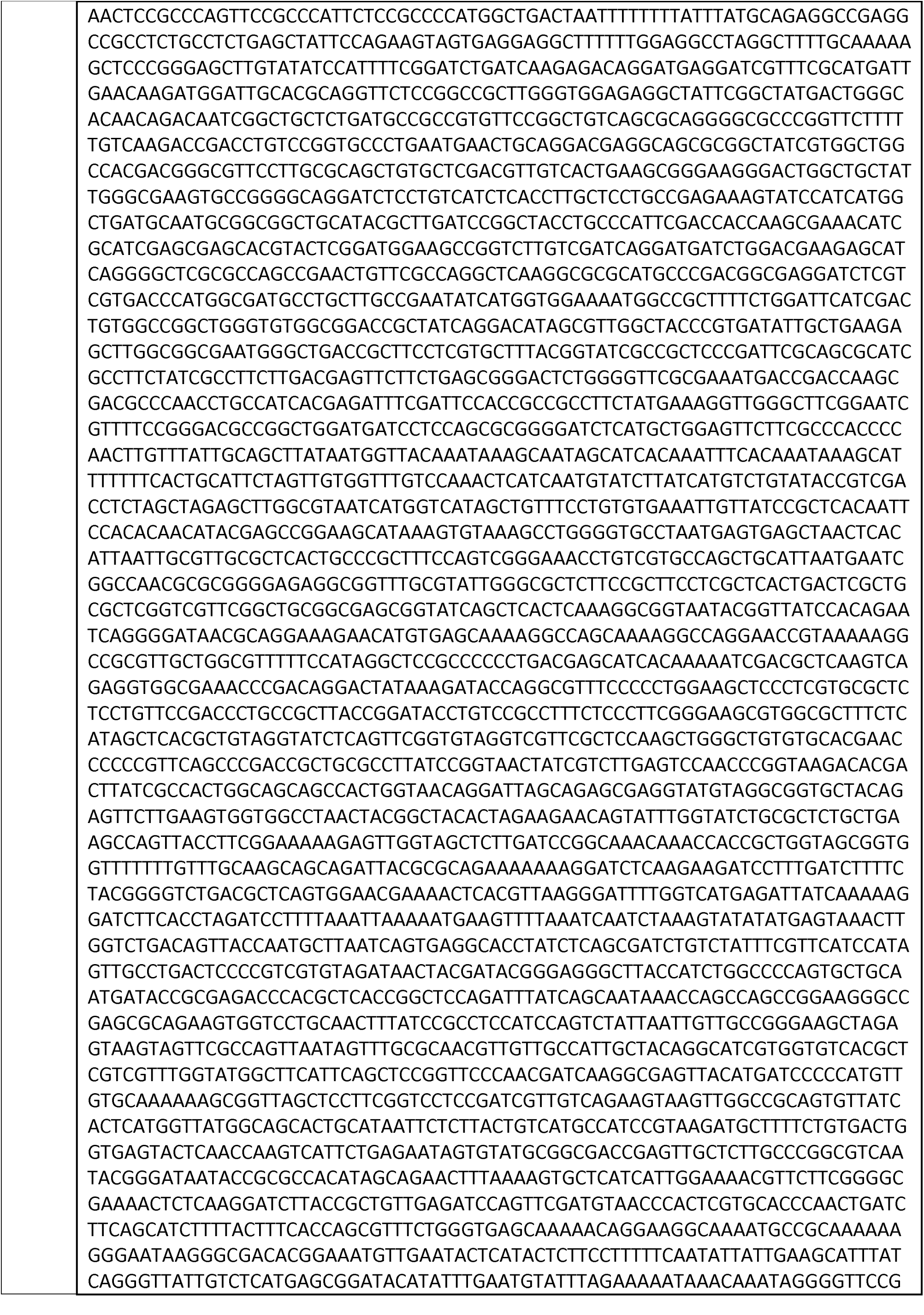
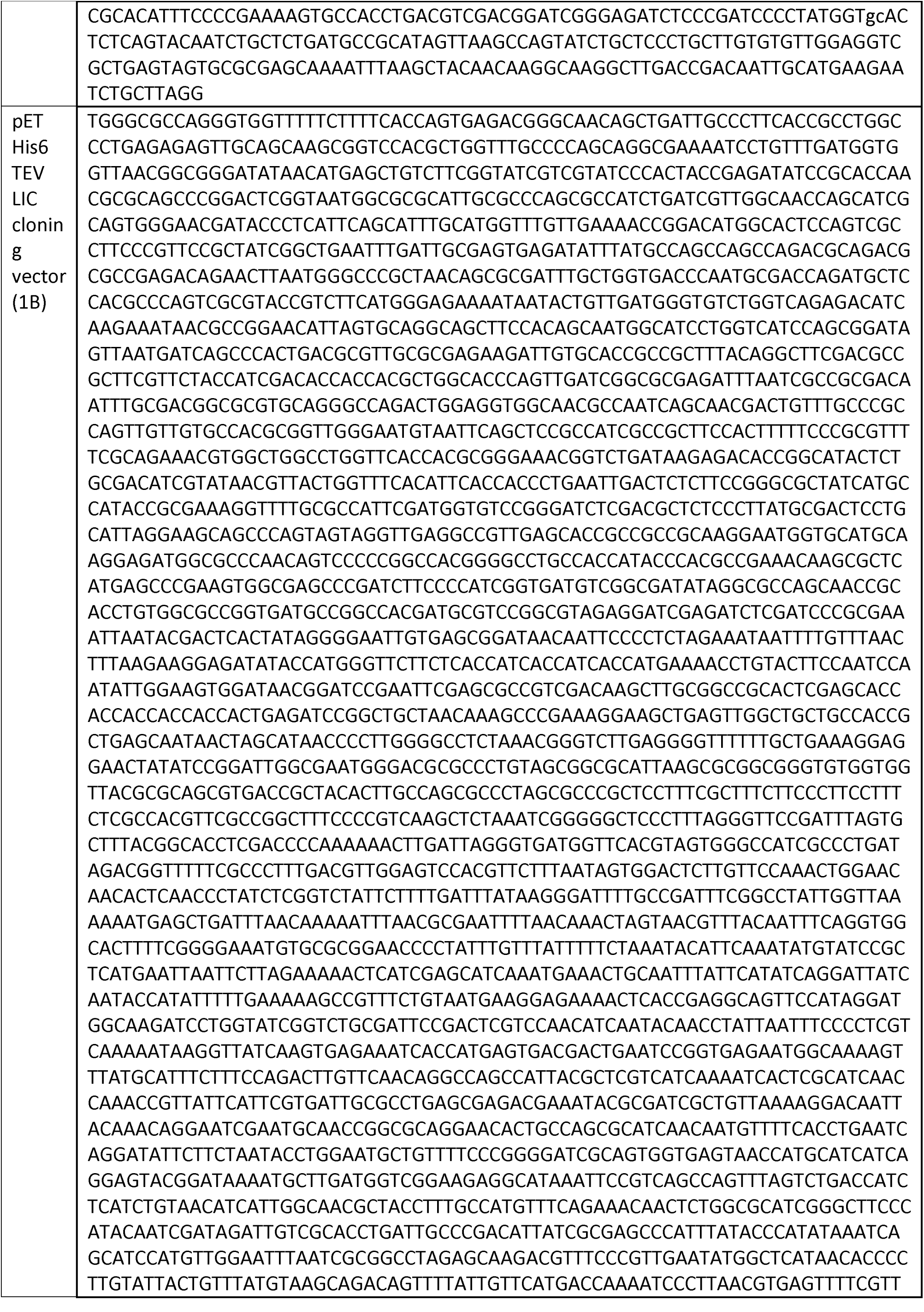
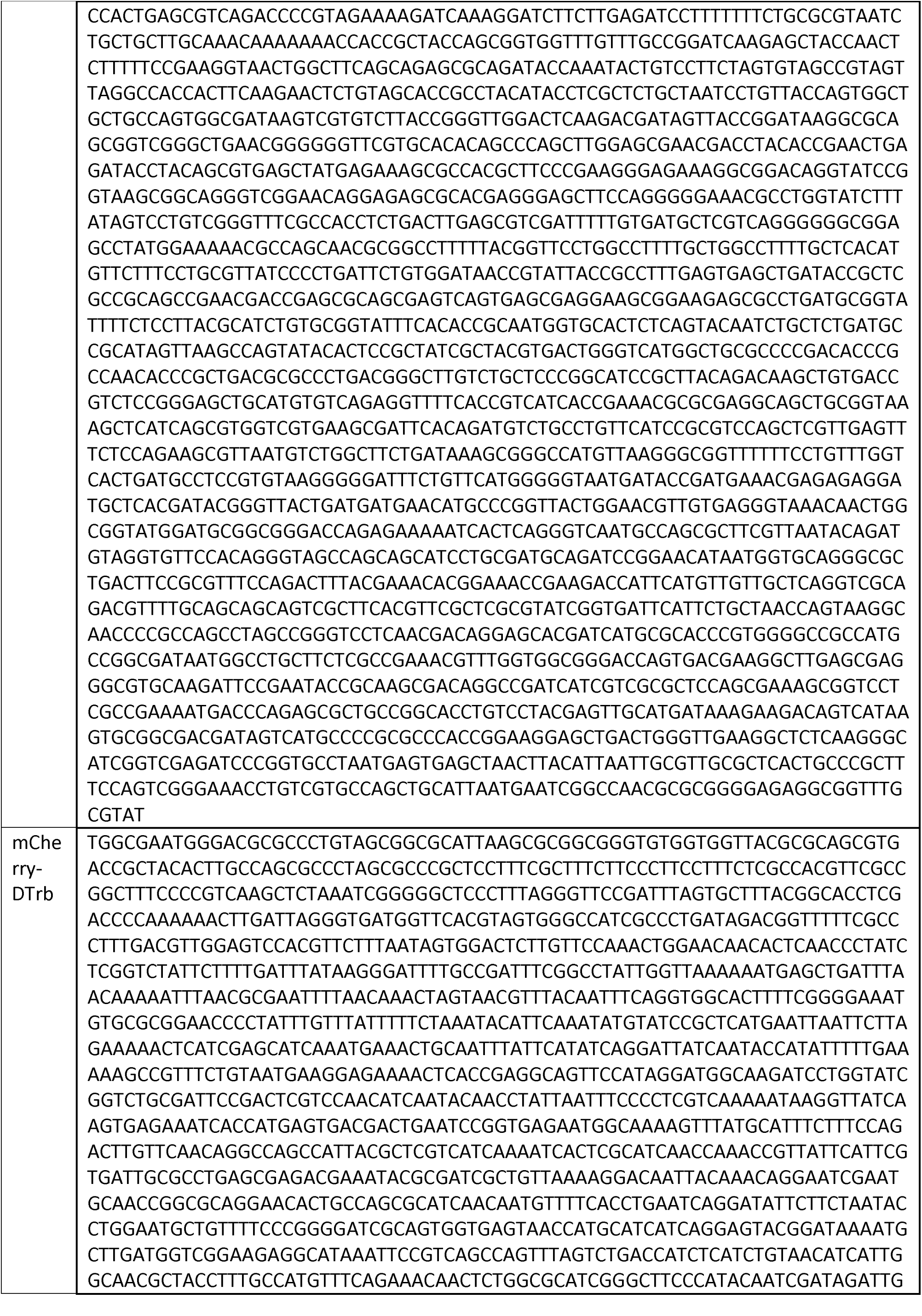
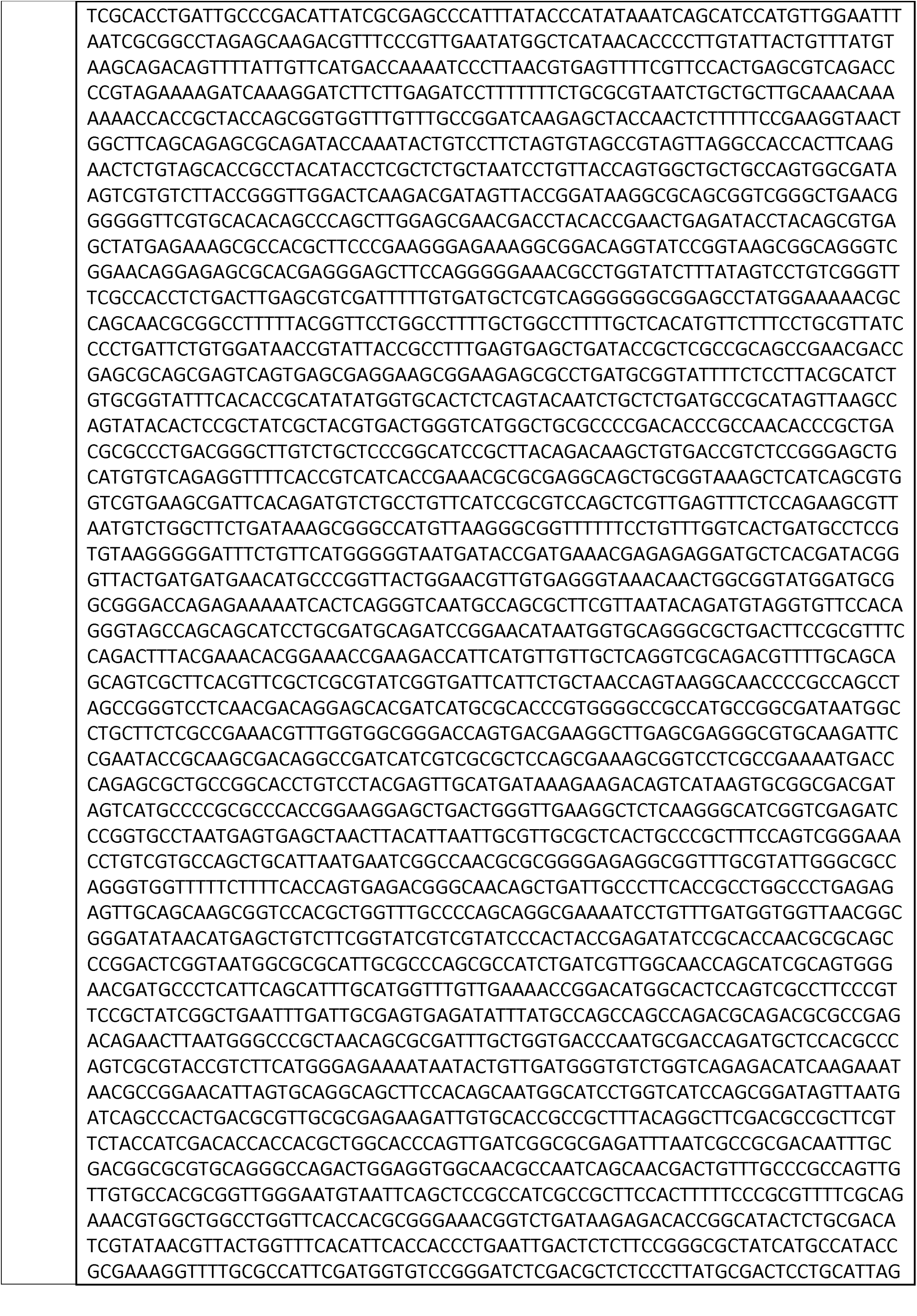
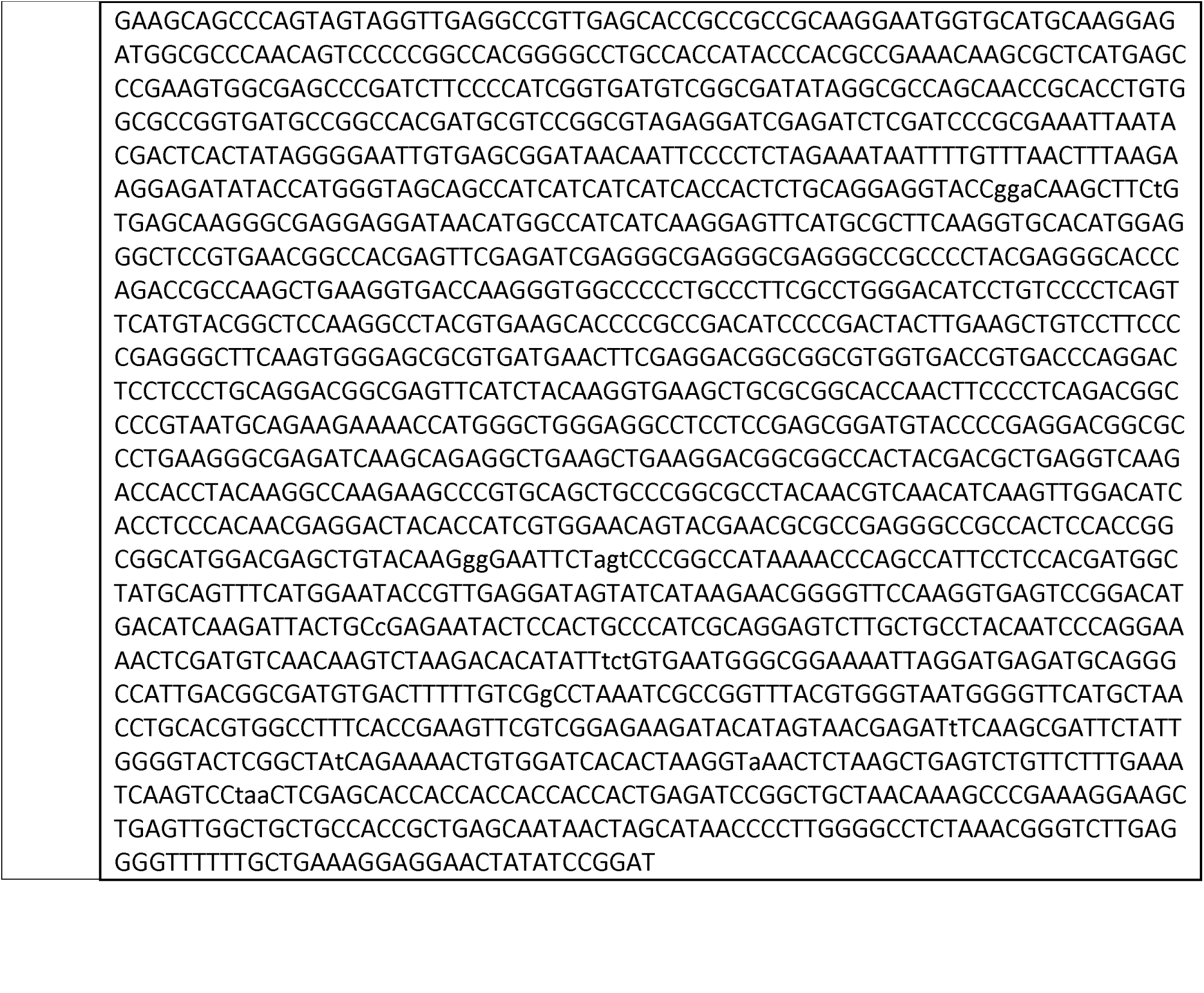

## Notes

### Competing Interest Statement

The authors have declared no competing interest.

